# Evolution of promoter-proximal pausing enabled a new layer of transcription control

**DOI:** 10.1101/2023.02.19.529146

**Authors:** Alexandra G. Chivu, Brent A. Basso, Abderhman Abuhashem, Michelle M. Leger, Gilad Barshad, Edward J. Rice, Albert C. Vill, Wilfred Wong, Shao-Pei Chou, Gopal Chovatiya, Rebecca Brady, Jeramiah J. Smith, Athula H. Wikramanayake, César Arenas-Mena, Ilana L. Brito, Iñaki Ruiz-Trillo, Anna-Katerina Hadjantonakis, John T. Lis, James J. Lewis, Charles G. Danko

## Abstract

Promoter-proximal pausing of RNA polymerase II (Pol II) is a key regulatory step during transcription. Despite the central role of pausing in gene regulation, we do not understand the evolutionary processes that led to the emergence of Pol II pausing or its transition to a rate-limiting step actively controlled by transcription factors. Here we analyzed transcription in species across the tree of life. Unicellular eukaryotes display a slow acceleration of Pol II near transcription start sites that transitioned to a longer-lived, focused pause in metazoans. This event coincided with the evolution of new subunits in the NELF and 7SK complexes. Depletion of NELF in mammals shifted the promoter-proximal buildup of Pol II from the pause site into the early gene body and compromised transcriptional activation for a set of heat shock genes. Our work details the evolutionary history of Pol II pausing and sheds light on how new transcriptional regulatory mechanisms evolve.

## Introduction

The evolution of new mechanisms to control the transcriptional cycle were required for the specialization of distinct cell types from a single genome sequence^1–3^. These additional stages of transcriptional control enabled the regulatory sophistication required for multicellularity, embryonic and developmental regulatory states, and the evolution of sophisticated body plans^1,4–8^. However, some transcriptional regulatory steps found in animals are not present in bacteria, or even in unicellular eukaryotes^9,10^. This raises a fundamental question about how regulatory complexity evolves when changes affect an essential cellular process like transcription.

Promoter-proximal pausing is a key regulatory stage during transcription by RNA Polymerase II (Pol II), where Pol II pauses 20-60 bases downstream of the transcription start site (TSS). This promoter-proximal pausing (herein, a pause) is ubiquitous in *Drosophila* and mammals and can disrupt the continuous flow of transcription for durations of 1-15 minutes (**Fig. 1A**)^11–14^. The rate at which polymerases are released from a paused state into productive elongation is actively regulated by transcription factors^15^, and is therefore essential for proper development in most animal species^16–18^. Proteins in the Negative Elongation Factor (NELF) and DRB Sensitivity Inducing Factor (DSIF) complexes are sufficient to establish pausing^14,19^. In addition, SPT6 has recently been argued to play an even more important role in establishing a pause *in vivo*^20^. Pause release is mediated by the P-TEFb protein complex, specifically, the kinase CDK9 which phosphorylates multiple targets including NELF, DSIF, and the Pol II C-terminal domain^21^. Another player in the duration Pol II spends paused is the 7SK snRNP complex (HEXIM, LARP7, MEPCE and its noncoding 7SK RNA binding partner), which inhibits P-TEFb to prevent pause release^22^. However, despite the importance of pausing and pause-related proteins in mammals, certain model organisms such as yeast, do not have a pause^7,8,23,24^. This phylogenetic disparity raises the question of how and when the pause evolved, as well as which factors mediated the original pause.

**Figure 1:**
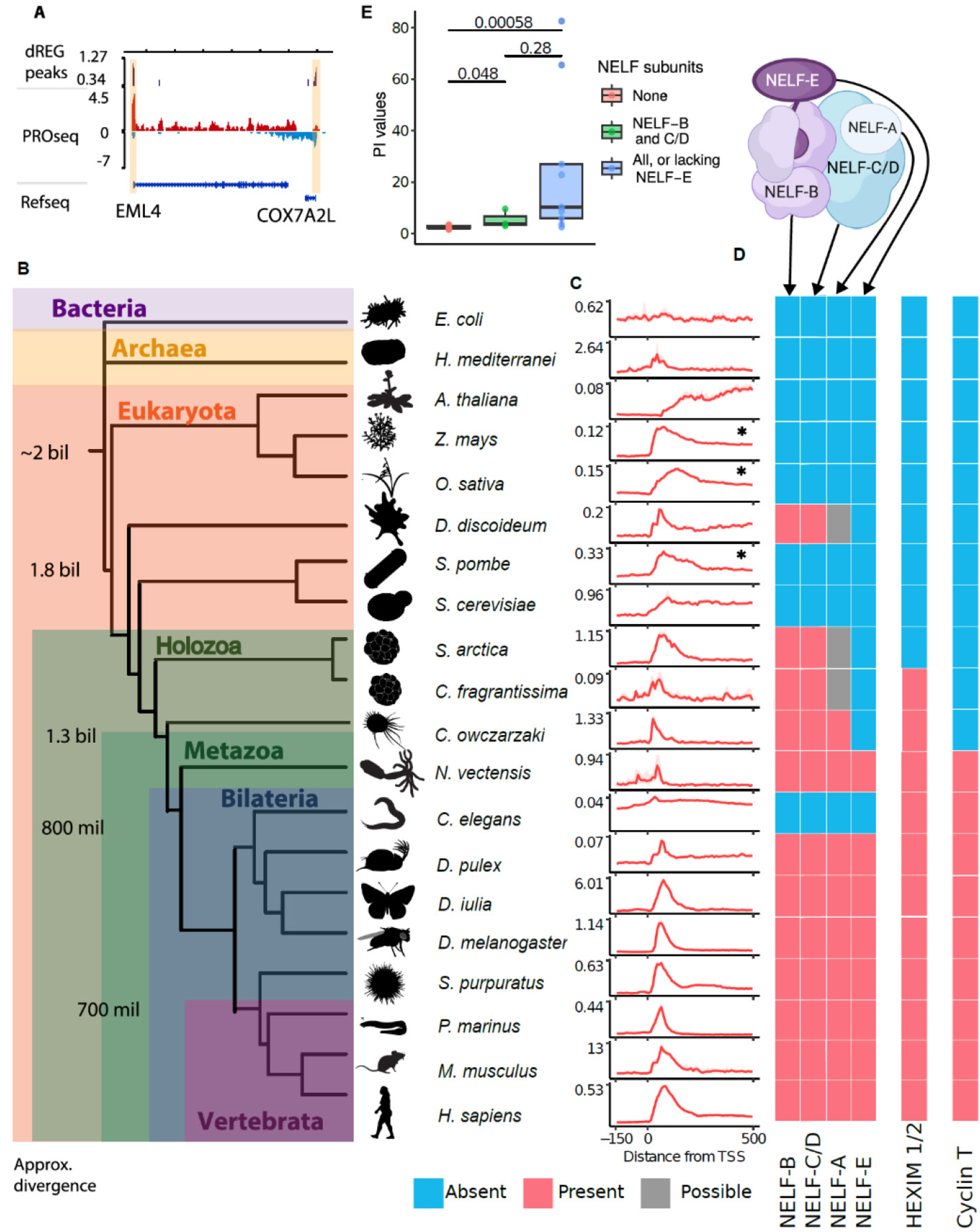
The evolution of NELF subunits is associated with pausing. (A) Depiction of a PRO-seq track where red and blue represent reads mapped to the plus and minus strands, respectively. dREG peaks are marked in purple and pause regions are highlighted in yellow. (B) Schematic depicting the relationships between the 20 species included in this study. Divergence times were taken from Parfrey *et al.*^81^ and Roger *et al.*^82^. (C) Metaprofiles of PRO-seq data were collected in each species. The dotted line marks the position of TSSs. The 25-75% confidence intervals are depicted in transparent red. Species which show proto-pausing indicated by *, vertical axis represents the mean PRO-seq signal. (D) Cartoon depicting internal interactions in the NELF complex based on the crystal structure in ^35^ above colored blocks denote the presence (red), absence (blue), or possible presence (gray) of the human orthologues of NELF subunits in each species as inferred from reciprocal blast searches. (E) Box and whiskers plot of pausing indexes in each species. Boxes are clustered by the number of NELF subunits in each species. A Mann-Whitney test was used to compute p-values.

Here, we studied the evolution of Pol II promoter-proximal pausing by performing Precision Run-On and Sequencing (PRO-seq) to examine the transcription profiles of twenty organisms. Integration of PRO-seq data and phylogenetic analysis revealed that proteins in the NELF complex were crucial for the emergence of pausing early in animal evolution, highlighting their primacy over alternative pausing factors such as DSIF and or SPT6, which we found to be conserved throughout Eukarya. To understand what the pause contributed to gene regulation in animal species, we employed the dTAG system to degrade two NELF subunits, NELF-B and NELF-E. We found that degradation of either NELF subunit prevented signal-regulated gene activation by transcription factors targeting gene expression by pause release, as demonstrated for heat-shock factor 1 (HSF1). Collectively, our findings show how pausing evolved in Metazoa and provided a new rate-limiting step during transcription that can be regulated to fine-tune mRNA production.

## Results

### Promoter-proximal Pol II residence time increased during eukaryote evolution

We used PRO-seq^25^ to study transcription in 20 extant organisms that represent two billion years of evolution (**Fig. 1A-B**), including multiple early branching metazoans and non-metazoan eukaryotes. Our atlas of organisms adds new data representing two prokaryotic organisms (*Escherichia coli* and *Haloferax mediterranei*, representing the bacterial and archaeal domains, respectively), single-celled eukaryotes including close relatives of animals - two ichthyosporeans (*Creolimax fragrantissima,* and *Sphaeroforma arctica*), a filasterean (*Capsaspora owczarzaki*) - and the model social amoeba (*Dictyostelium discoideum*). The metazoans studied represent major taxa, including the cnidarian (*Nematostella vectensis*), the sea urchin (*Strongylocentrotus purpuratus*), the water flea (*Daphnia pulex*), the butterfly (*Dryas iulia*), and the cyclostome (*Petromyzon marinus*). For each new organism, we prepared PRO-seq libraries from at least two independent biological replicates, which were well correlated with each other (**Supplemental Fig. 1**). Additionally, we augmented our PRO-seq atlas by integrating published PRO-seq, GRO-seq, or ChRO-seq data from a fly (*Drosophyla melanogaster*^25^), a nematode (*Caenorhabditis elegans*^26^), yeast (Saccharomyces *cerevisia*e and *Schizosaccharomyces pombe*^23,24^), model plants (*Arabidopsis thaliana*, *Oryza sativa*, and *Zea mays*^27–29^), and mammals (*Homo sapiens* and *Mus musculus*^17,30^). Throughout the manuscript, we examined the sensitivity of major results to the data source and/ or details of the run-on and sequencing assay library prep, finding no association with any technical factors (**Supplemental Fig. 2**). These species occupy key phylogenetic positions relative to animals, allowing us to investigate most major transitions in the animal stem lineage.

To evaluate the presence of a promoter-proximal pause in a way that is agnostic to the quality of genome assembly and gene annotations in each organism, we developed a computational strategy to refine the transcription start sites using PRO-seq data (see Methods; **Supplemental Fig. 3**). As expected, most metazoan organisms exhibited a pileup of RNA polymerase 30-100 base pairs downstream of the TSS indicative of Pol II promoter-proximal pausing (**Fig. 1C**). Conversely, the bacterium *E. coli* lacked a prominent RNA polymerase peak in our data. Plants and unicellular eukaryotes exhibited more diverse pause variation: the plant *Z. mays* showed a focused peak, while the plant *O. sativa* and the yeast *S. pombe* displayed a more dispersed peak downstream of the TSS. The plant *A. thaliana* and yeast *S. cerevisiae* showed no evidence of pausing.

To reveal more subtle differences in the dynamics and gene-by-gene variation at the pause site than observed in meta profiles, we computed pausing indexes which quantify the relative duration Pol II spends in a promoter-proximal paused state^13,31^. Pausing indexes revealed that the residence time of Pol II at the pause site is, on average, 1-2 orders of magnitude higher in metazoans than unicellular eukaryotes or plants (**Supplemental Fig. 4**). Consistent with the meta plots, we also noted wide variation in pausing indices even in species without evidence of a focal pause. These major differences across taxa were evident when examining either PRO-seq, GRO-seq, or ChRO-seq datasets separately from the remaining samples, indicating that differences between taxa were not driven by technical differences between experimental approaches (**Supplemental Fig. 2**). Taken together, our results suggest that an ancestral slowdown in Pol II transcription near the TSS may have arisen in unicellular eukaryotes and become longer in duration and more focused during the early evolution of metazoans.

### NELF and 7SK subunits associated with increased residence time of paused Pol II

Pausing is established by interactions between Pol II and proteins in the DSIF and NELF complexes^32,33,34^. In addition to these proteins being necessary for establishing a pause, the duration Pol II spends in a paused state is further influenced by the p-TEFb and 7SK snRNP complexes^21,22^. To determine which proteins were associated with pausing in eukaryotes, we used sequence similarity and profile hidden Markov model protein domain searches to identify potential orthologs of the human subunits in public sequence repositories.

The NELF complex consists of multiple subunits, including NELF-A, -B, -C (or its isoform -D), and -E (**Fig. 1D**). We found that NELF-B and -C/-D are widely distributed in eukaryotes (**Fig. 1D**; Supplemental Table 1). Both subunits appear to have been secondarily lost in yeast, *S. pombe* and *S. cerevisiae*, as well as in the nematode *C. elegans* and in land plants. CryoEM studies revealed that NELF-B and -C/-D interact with one another forming a core structure that holds the entire NELF complex together^35–37^(**Fig. 1D**). Our results indicate that this core NELF structure was most likely present in a shared common ancestor of all eukaryotes.

Phylogenetic analysis revealed that the NELF-A and NELF-E subunits were more recently derived (**Supplemental Fig. 5**). Apparent homologs of NELF-A are present in some unicellular eukaryotes. However, the NELF-A hepatitis delta antigen (HDAg)-like domain, crucial for RNA Polymerase II binding and transcriptional pausing^36,38,39^, was identified only in animals and *C. owczarzaki* (**Supplemental Fig. 6**). NELF-A gene annotations in apicomplexan parasites, such as *Plasmodium* species, do not appear to have this domain (Supplemental Table 1; **Supplemental Fig. 6**). These results are potentially consistent with a model in which NELF-A evolution occurred through the addition of an HDAg-like domain in an existing protein of unknown function. We found strong evidence for NELF-E only in metazoans (**Supplemental Fig. 5**); the only similarity in unicellular eukaryotes was with an RNA-binding domain shared with many other proteins (**Supplemental Fig. 7**). The most parsimonious model is therefore that NELF-E evolved early in metazoan evolution or just before the emergence of Metazoa.

In contrast to NELF, the DSIF elongation complex (SPT4 and SPT5), which also has a role in establishing paused Pol II, were universally conserved across all eukaryotes (**Supplemental Table 1**), and in some cases in archaea and/or bacteria^40,41^. Likewise, SPT6, which was recently proposed as a primary protein responsible for promoter-proximal pausing^20^, was also conserved across Eukarya.

In addition to changes in NELF composition, we also identified changes in protein complexes that establish the duration that Pol II spends in a paused state. The 7SK snRNP complex (HEXIM, LARP7, MEPCE and its noncoding 7SK RNA binding partner), which binds and inhibits P-TEFb^42^, also underwent recent evolutionary changes. Orthologs of HEXIM proteins (due to a recent duplication, HEXIM1 and HEXIM2 in mammals^43^) were found in choanoflagellates, ichthyosporeans and possibly *Capsaspora*, as well as patchily distributed HEXIM-like sequences in a small number of other eukaryotes (**Supplemental Table 1; Supplemental Fig. 5**). However, functionally important threonine, phenylalanine and tyrosine residues involved in P-TEFb inhibition^22^ were absent from many of these sequences (**Supplemental Table 1; Supplemental Fig. 8**). The most parsimonious model is that HEXIM proteins evolved in a shared ancestor of metazoans and their closest single celled relatives.

The cyclin dependent kinase family, responsible for pause release, have a conserved role phosphorylating the Pol II C-terminal domain in most eukaryotes^24,44^. Gene duplications in CDKs, likely present in the last eukaryotic common ancestor, led to the emergence of separate families representing CDK9 and closely related paralogs CDK12/ CDK13, possibly at the onset of opisthokonts (the clade sharing a most recent common ancestor with fungi and animals) (**Supplemental Table 1; Supplemental Fig. 5**). In *A. thaliana*, CDKC;2, an orthologue of either CDK9 or CDK12/13, is reported to phosphorylate the RNA polymerase C-terminal domain ^44,45^, demonstrating a consistent molecular function of CDK9 paralogues back at least as far as the divergence of animals and plants. Thus, the Pol II kinase activity of p-TEFb paralogs appears to have emerged early during eukaryotic evolution. Cyclin-T, however, most likely originated relatively recently in a shared ancestor of choanoflagellates and animals^46^ through duplication of an ancestral Cyclin-T/K gene, coinciding roughly with the appearance of NELF-E and HEXIM (**Supplemental Table 1; Supplemental Fig. 5**). Thus, several independent cyclin dependent kinases which phosphorylate Pol II have apparently evolved to act together in p-TEFb, perhaps enhancing regulatory control of this positive elongation factor. These results illustrate how p-TEFb evolution was linked to changes in NELF and 7SK protein composition.

Our analysis led us to suspect that progressive changes to NELF and HEXIM protein subunits, perhaps coupled with the addition of Cyclin-T, may have resulted in incremental alteration of the residence time of paused Pol II along the animal stem lineage. To determine how the addition of multiple new proteins affected the strength of pausing, we compared pausing indexes between species with a different complement of NELF or HEXIM subunits. Species containing NELF-B and -C/-D (which we refer to as the “core” NELF complex) have higher pausing indexes than species without any NELF subunits (**Fig. 1E; Supplemental Fig. 4**). Thus, the core NELF subunits are sufficient to induce some promoter-proximal pause on their own. The addition of NELF-A, -E, and HEXIM increased pausing indexes to the same order of magnitude observed in flies and mammals. Thus, NELF-B and -C/-D are sufficient for pausing, but the addition of NELF-A, NELF-E, and HEXIM proteins correlate with higher pausing indices and suggests the derived proteins may act together with the core NELF complex to fine-tune the function of paused RNA polymerase (**Supplemental Fig. 9A-B**).

### Nucleosome positioning and DNA sequence correlate with the position of a proto-pause

In some unicellular organisms, we observed a dispersed pause-like behavior through the first 200 bp after the TSS. We hypothesized that this “proto-pause” may serve as an ancestral substrate pre-dating the highly focused, long-duration Pol II pausing observed in extant metazoans, perhaps symptomatic of slow moving Pol II as observed in Booth et al.^24^. The pronounced proto-pause in *Z. mays*, *S. pombe*, and *O. sativa* appears in a similar location to a canonical pause (or just downstream), but does not have the same magnitude of pausing index or focal peak (**Fig. 1C, E**).

To explore why some extant species have a consistently positioned proto-pause despite not containing any of the NELF subunits, we examined the DNA sequence under the strongest pause-like position in each organism. Consistent with previous work^25,47–51^, we found that in metazoan organisms, Pol II tends to pause on a C nucleotide after a relatively G-rich DNA sequence (**Fig. 2A**). A similar sequence preference was also present in three organisms that show a proto-pause, including *Z. mays, S. pombe,* and *O. sativa* (**Fig. 2B**). The candidate pause motif has a low information content, yet its consistent positioning in pause and proto-pause regions in many organisms suggests biological significance. To explore the importance of the pause motif, we analyzed PRO-seq data in three tissues isolated from F1 hybrid mice, bred from CAST and B6 parents^48^. We tested whether DNA sequence variation in the pause motif between CAST and B6 alleles impacted either the pausing index or pause position, given a shared primary TSS. Our findings revealed that stronger pause motifs on the maternal allele were associated with a higher pausing index on the maternal allele in all three tissues (and vice versa; **Fig. 2C, Supplemental Fig. 10B**). Thus, DNA sequence changes in the pause motif were correlated with changes in the pausing index.

**Figure 2:**
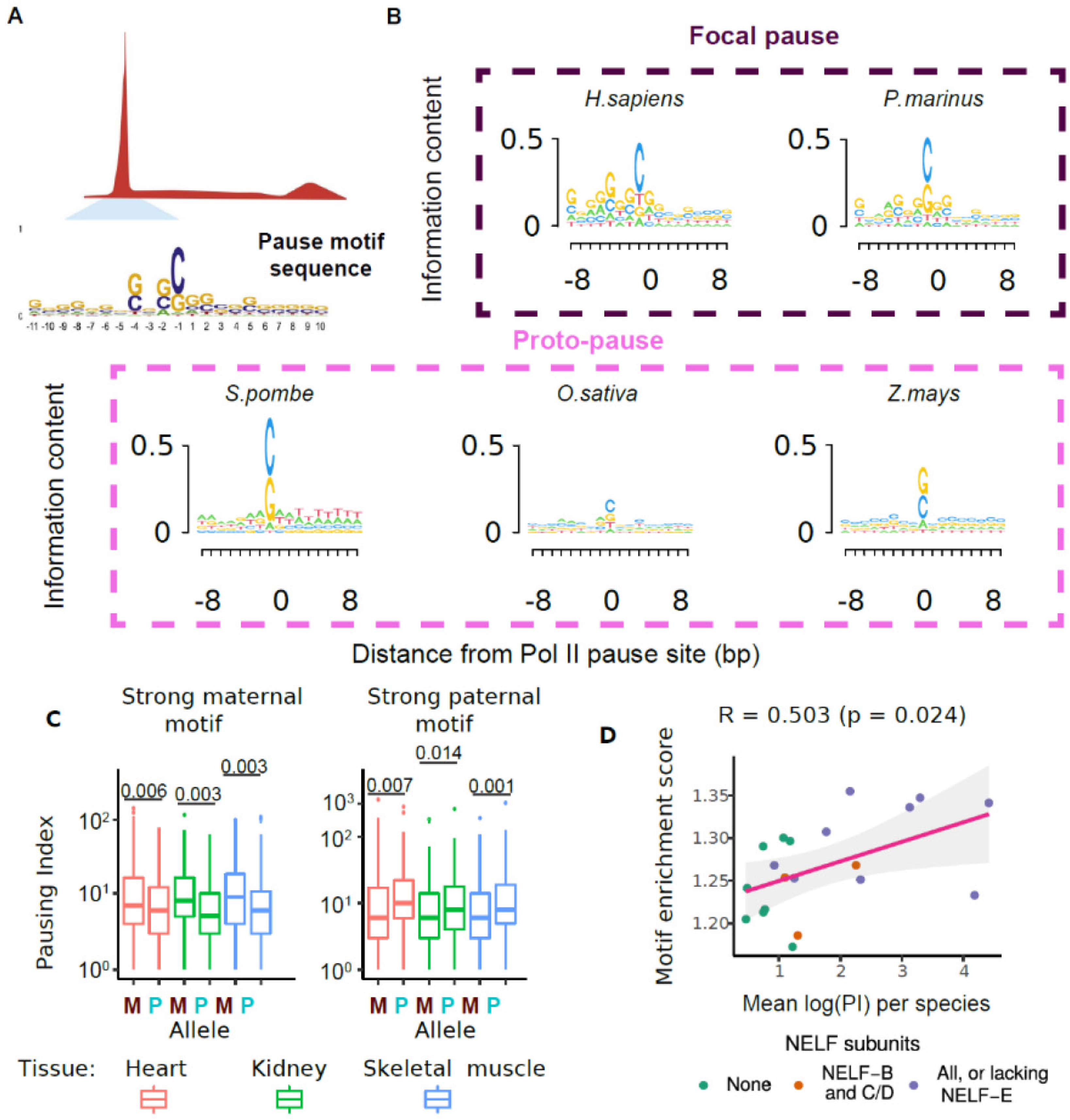
Genomic features are associated with pausing. (A) Schematic of a motif search at the Pol II pause site including the human pause motif sequence as published in Watts *et al*.^49^ (B) DNA sequence motif under the active site of paused Pol II in organisms with a focal pause or a proto-pause. The size of each base is scaled by information content. (C) Boxplot of pausing indexes for genes with stronger pause motif on the maternal (left) or paternal (right) alleles in F1 hybrid mice. (D) Scatter plot denotes the enrichment of the motif score relative to flanking DNA and the mean pausing index in each species. Each dot is colored by the number of NELF subunits found in each sample. We fit a linear regression to derive the R^2^ and the p-value.

To determine whether the pause motif was associated with pausing index variation across all 20 species, we examined the enrichment of the pause motif near the pause position (**Fig. 2D, Supplemental Fig 10A**). Despite the pause motif being derived from humans^49^, we nevertheless found that it explained over 25% of the variation in pausing index across all organisms (Pearson’s R = 0.503, *p* = 0.024). Conversely, the DNA sequence motif of the Initiator, a highly conserved and precisely positioned promoter element^52,53^, was not correlated with pausing index (**Supplemental Fig. 10C-D**). Altogether, our data support the idea that a pause sequence motif, featuring a C in a G-rich background precisely located downstream of the TSS correlated with pausing index, suggesting that it serves as an ancestral cis-regulatory DNA sequence that may contribute to the formation of proto-pausing.

Next, we examined the relationship between +1 nucleosome positioning and pausing using MNase sequencing data in model organisms^54–57^ with different pause status. Most strikingly, the position of the +1 nucleosome dyad was consistently over 100 bases downstream of the TSS in two metazoan organisms with a focal pause and a high pausing index (**Fig. 3A-B**). Conversely, the +1 nucleosome was positioned closer to the TSS in species with a proto-pause or no pausing. In the proto-paused organism *S. pombe*, the +1 nucleosome dyad was located in the same position as the pause, approximately 75 bp downstream of the TSS, while in the unpaused *S. cerevisiae*, the nucleosome dyad was found even closer to the TSS^58^ (**Fig. 3C**). The nucleosome dyad provides more resistance to Pol II elongation than transcription through other parts of the nucleosome^59^, suggesting that the observed slow-down of Pol II at this position in *S. pombe*, and perhaps other proto-paused organisms, is due in part to a nucleosome barrier. Furthermore, these results are consistent with a model in which the pause functions in part to exclude a nucleosome from encroaching upon the TSS^56^.

**Figure 3:**
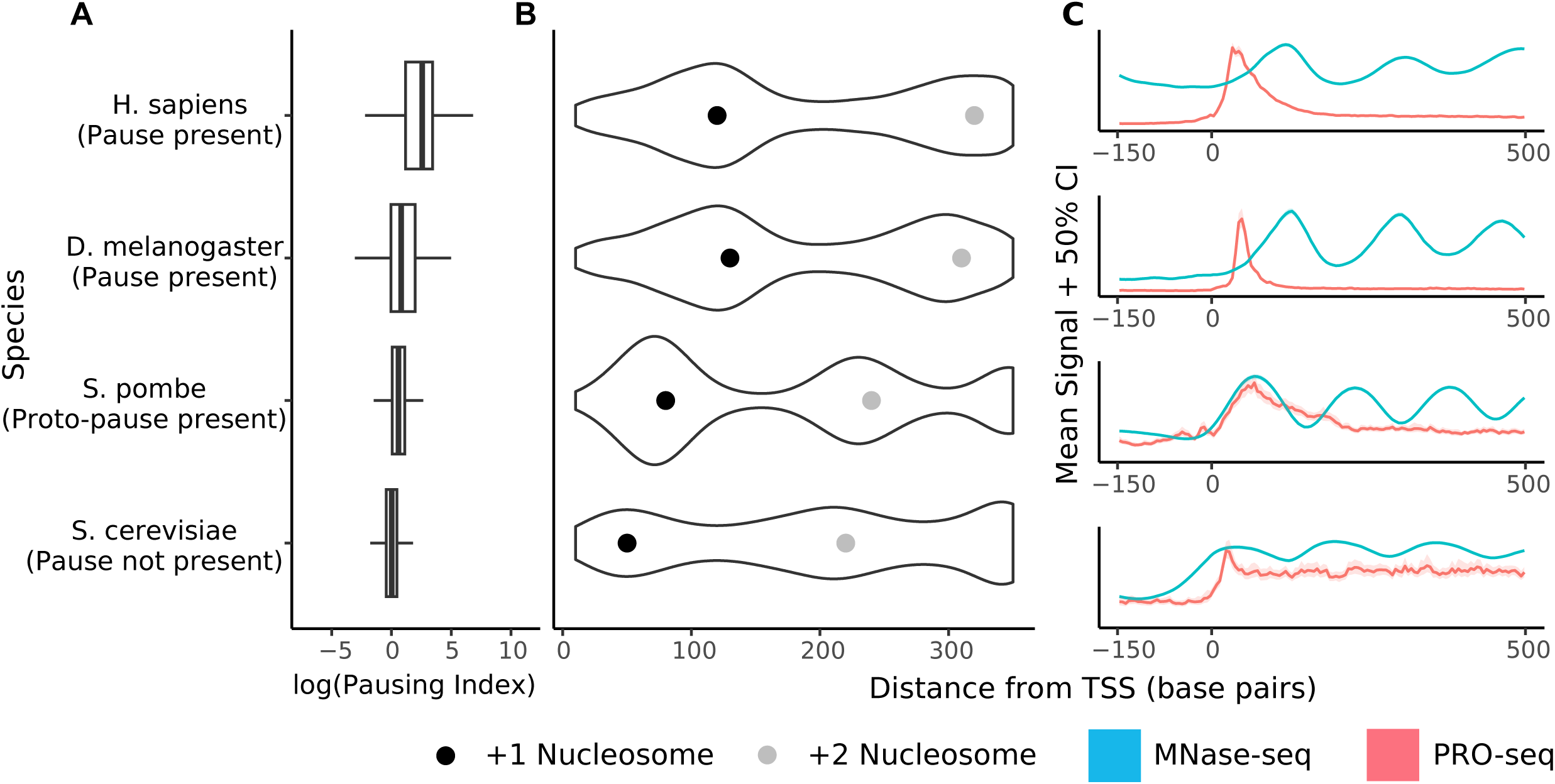
Nucleosome positioning is associated with pausing. (A) Boxplot of pausing indexes for selected species representative of paused (*H. sapiens*, *D. melanogaster*), proto-paused (*S. pombe*), and unpaused (*S. cerevisiae*) (B) Violin plot of MNase-Seq data with the most common +1 (black) and +2 (gray) nucleosome positions marked with a dot. Species with a high pausing index show downstream +1 nucleosome positioning. Species with no pausing or proto-pausing show further upstream +1 nucleosome. (C) Metaplot of PRO-seq signal (red) alongside MNase signal (blue). The pause in H. sapiens and D. melanogaster occurs at a position distinct from the +1 nucleosome while the position of a proto-pause coincides with the +1 nucleosome. In *S. cerevisiae*, the +1 nucleosome overlaps with the TSS.

These results, and previous studies^23–25,60,61^, suggest a model in which multiple factors contribute to the formation of the proto-pause. In all eukaryotic organisms, newly initiated Pol II is not yet ready to move efficiently through barriers to transcription, such as difficult to transcribe DNA sequences or nucleosomes. In paused organisms, NELF serves as a strong checkpoint, providing a position close to the TSS at which Pol II acquires competence to surmount these barriers, either through the phosphorylation of the Pol II CTD or associated elongation factors (DSIF), or the addition of new protein complexes (PAF, FACT, SPT6). Taken together, our results show that in organisms without NELF, these barriers, in particular the +1 nucleosome and a G-C rich DNA sequence motif, become critical impediments to Pol II transcription, and therefore their impact on residence time is more apparent.

### Loss of core NELF-B impacts chromatin localization of NELF-E and alters Pol II pausing

Our analysis of NELF evolution shows that the core NELF subunits, NELF-B and -C/D, evolved earlier than the ancillary subunits, NELF-A and -E. To test the functional impact of core and ancillary subunits in mammalian cells, we generated FKBP12-homozygously tagged mouse embryonic stem cell (mESC) lines that rapidly degrade either NELF-B or NELF-E after treatment with the small molecule dTAG-13 (**Fig. 4A-C**^17,62^). The NELF-B dTAG was reported and validated in a recent paper^17^, while the NELF-E dTAG cell line is novel here. We verified that the FKBP12-tagged NELF-E protein was properly localized and that NELF-E was nearly undetectable within 30 min after the addition of 500 nM dTAG (**Supplemental Fig. 11; Supplemental Fig. 12**). We also verified that the rapid depletion of both NELF subunits decreased Pol II levels at the pause site following 30-60 min of dTAG-13 treatment, as measured by PRO-seq (**Fig. 4D; Supplemental Fig. 12A**).

**Figure 4:**
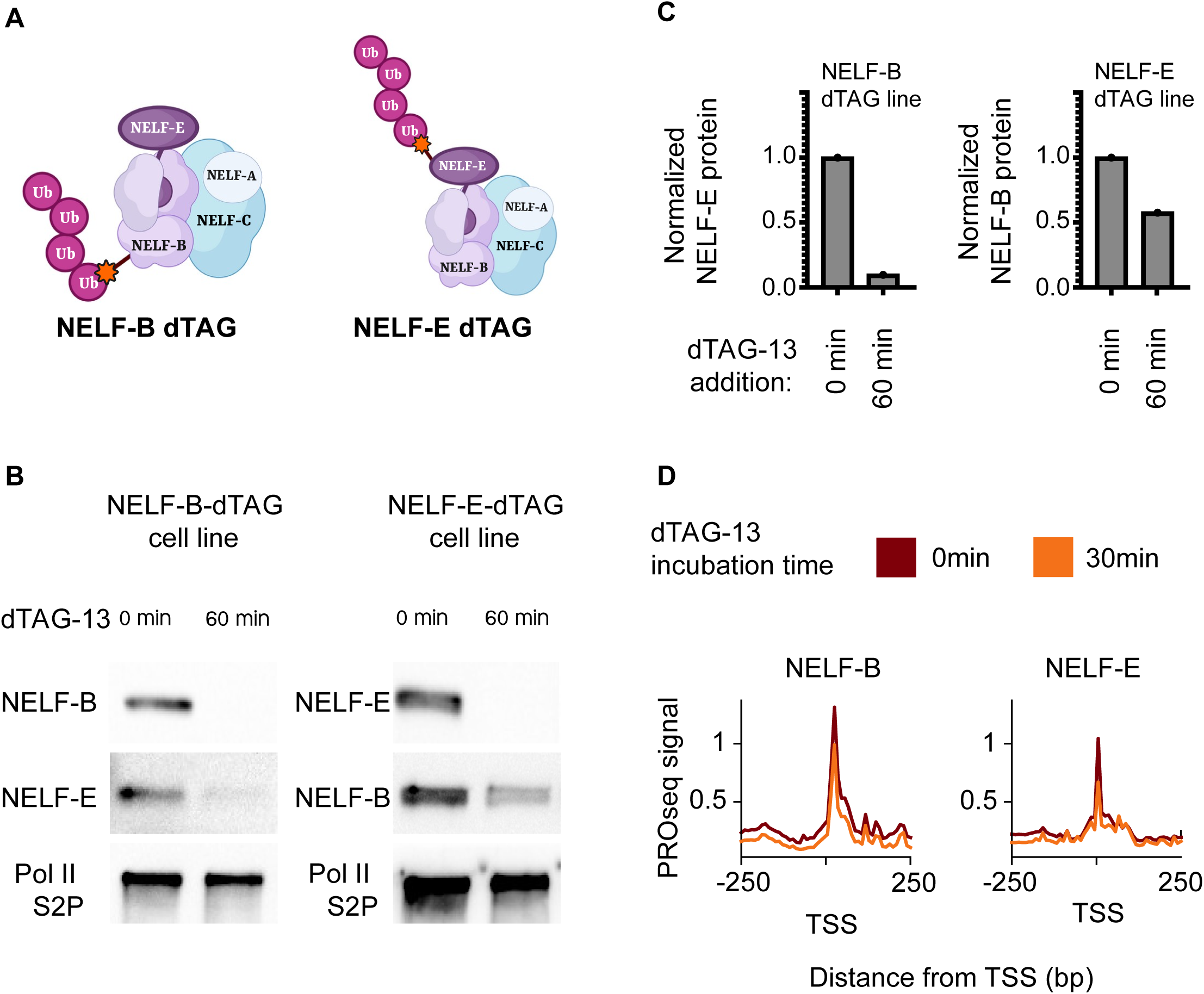
NELF degradation destabilized RNA Pol II pausing. (A) Schematic of NELF-B or NELF-E degradation mESC cell lines. (B) Western blots depict NELF-B, NELF-E and Pol II after the degradation of either NELF-B (left) or NELF-E (right) using 500nM dTAG-13. (C) Quantification of NELF-E western blot signal after NELF-B degradation (left) and NELF-B after NELF-E degradation (right). (D) Meta profiles of PRO-seq signal at 0min (red) and 30min (orange) after the degradation of NELF-B (left) or NELF-E (right).

We hypothesized that loss of NELF-B would have a greater impact on NELF complex assembly on chromatin than loss of NELF-E due to the central role of NELF-B in the complex (**Fig. 4A**^35^). Consistent with our hypothesis, we observed that loss of NELF-B led to a decrease of the entire NELF-B/E sub-complex from chromatin, while a loss of NELF-E resulted in only a moderate reduction of 40% in NELF-B protein levels (**Fig. 4B-C, left panel; Supplemental Fig. 11A-B, 12**). Consistent with changes in the NELF proteins themselves, we found that 30 min degradation of NELF-B decreased the quantity of promoter-proximal Pol II in PRO-seq data more than degradation of NELF-E (**Fig. 4D**, **Fig. 5A (3 clusters), Supplemental Fig. 13A-C (unclustered data)**). These findings confirm that the functions of NELF-B and -E in mESCs mirror the structure and evolutionary history of these NELF subunits.

**Figure 5:**
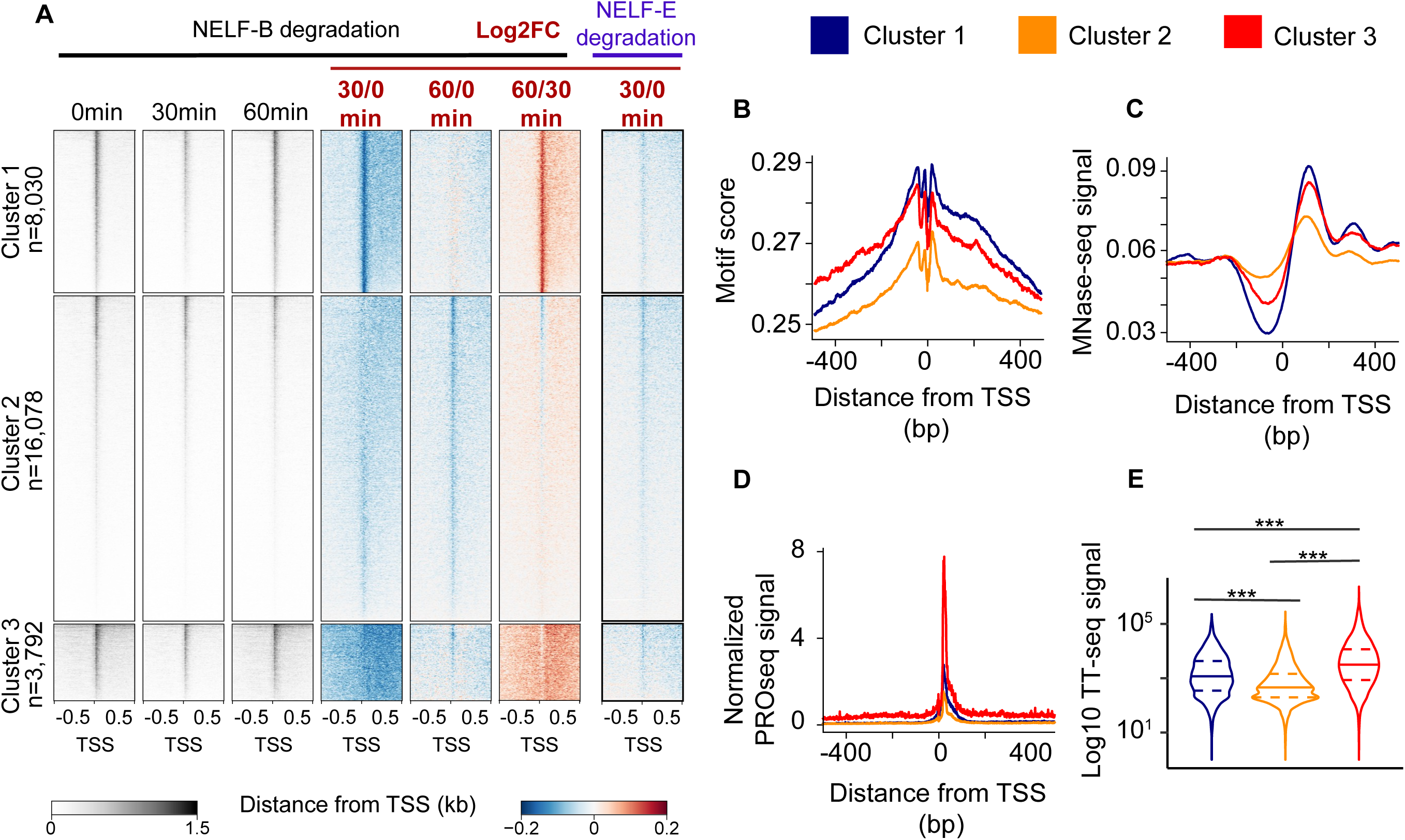
NELF degradation leads to different pause recovery profiles. (A) Heat maps of spike-in normalized PRO-seq signal after NELF-B or NELF-E degradation (left). Log2 fold changes of normalized PRO-seq signal relative to untreated controls are also depicted. (B-D) Pause motif enrichment scores (B), Meta profile of MNase-seq signal (C) and normalized PRO-seq signal (D) are depicted for three clusters of genes defined in panel (A). (E) Violin plots of log10 TT-seq signal in the three clusters defined in (A). A two-sided Mann-Whitney test was used to compute p-values. (***) defines p-values < 2.2e-16

### Pol II recovery after prolonged NELF-B degradation is in a similar position as a proto-pause

Our PRO-seq data in the NELF-B depleted cell line showed that many genes partially recovered Pol II near the pause site following 60 min of treatment (**Supplemental Fig. 13A-C**). To investigate the observed Pol II signal recovery, we first clustered genes based on their changes in Pol II loading between 30 and 60 min of dTAG treatment (**Fig. 5A**, clusters 1, 2, and 3). Cluster 1 showed a localized recovery of PRO-seq signal near the position of the canonical pause. Cluster 2 showed no indication of recovery and, relative to the other clusters, it was enriched in transcribed enhancer sequences (**Supplemental Fig. S14A**). Cluster 3 exhibited a recovery of Pol II further into the gene body (**Supplemental Fig. 13D, 15**), reminiscent of the slowdown of Pol II observed in *S. pombe* and *O. sativa*, potentially near the location of the +1 nucleosome as reported by a previous study^63^, and consistent with our hypothesis that the +1 nucleosome may act as a proto-pause positioning factor.

We hypothesized that after the depletion of NELF, DNA sequences associated with the pause in organisms without NELF-B may be sufficient to re-establish some paused Pol II. We looked for enrichment of the DNA pause motif at loci that exhibited recovery of the paused state after NELF depletion. We found both higher enrichment of the pause motif and better positioning of the +1 nucleosome in clusters 1 and 3, which show evidence of pause signal recovery, when compared to cluster 2, which does not (**Fig. 5B-C**). Interestingly, the main difference between clusters 1 and 3 was that genes in cluster 3 had higher effective initiation rates, as determined by both TT-seq^64^ and a computational modeling approach analyzing steady-state PRO-seq data^31^ (**Fig. 5D-E**). Genes in cluster 3 also had much higher binding of some components of the pre-initiation complex and more clearly defined DNA sequence motifs that specify transcription initiation^65^ (**Supplemental Fig. 14B-E**), potentially consistent with higher initiation rates. Based on these results, we propose that clusters 1 and 3 partially recover Pol II near the pause due to a combination of DNA sequence and interactions with well-positioned nucleosomes. We also speculate that genes in cluster 3 recover in a more downstream position as a result of a higher rate of initiation. Greater initiation rates at these genes may lead to an accumulation of Pol II at the start of the gene that causes polymerases to be pushed downstream due to interactions between newly incoming Pol II, or may cause a trickle of many incompletely phosphorylated elongation complexes in the absence of a focal checkpoint at which CDK9 can act.

In sum, we found that after NELF depletion, paused Pol II is temporarily and partially depleted on many genes, and has a recovery in signal at the same location as the proto-pause in multiple eukaryotic organisms that have lost the core NELF subunits. Furthermore, this pause-like recovery state is associated with the same functional features enriched in organisms with a proto-pause, and may be symptomatic of slow moving or non-productive Cdk9-mediated transcription^24,60^.

### Pol II pausing allows transcription factors to regulate pause release

Cyclin dependent kinases, which are responsible for pause release in metazoans, are conserved across eukaryotes, pointing to the critical role of phosphorylation of the Pol II CTD and DSIF in most eukaryotic organisms. In metazoans, sequence-specific transcription factors can tune the level of gene expression by modulating the recruitment of p-TEFb^15,16,66–69^. Thus, a previously necessary step during transcription was changed at some point during eukaryotic evolution into a rate-limiting step that provides a new means of transcriptional control. How this new regulatory step evolved is not known.

We hypothesized that introducing pause-release as a regulatory step in gene regulation required a focal pause, because the focal pause meant that the substrate of Pol II CTD phosphorylation was located in a fixed position relative to transcription factor binding sites. We previously showed that blocking CDK9 in *S. pombe* results in inefficient transcription across the early gene body, indicating that the substrate for CDK9 phosphorylation is distributed across several hundred bases of DNA^17,24,63^. Parallel to this observation, depletion of NELF in our mESC system, caused Pol II to “creep” across the beginning of the gene body^17^ (**Fig. 5A; Supplemental Fig. 15**). This suggests that Pol II, which needs to be phosphorylated by p-TEFb for efficient transcription elongation, may no longer be readily accessible in a fixed location after NELF depletion.

We hypothesized that the downstream redistribution of Pol II after NELF depletion would prevent the targeted regulation of gene expression by transcription factors acting to release paused Pol II into productive elongation. To test this hypothesis, we turned to the well-studied heat shock system, where the transcription factor heat shock factor 1 (HSF1) activates transcription of a core group of a few hundred genes following heat stress by the release of paused Pol II^67,70,71^. We asked whether HSF1 could release paused genes as efficiently following the depletion of NELF-B and -E in mESCs (**Fig. 6A**). We first identified genes that were up- and down- regulated using a regular heat shock experiment in mESCs. Our analysis confirmed the induction of a core group of heat shock-responsive genes^67,70,71^ (**Supplemental Fig. 16, 17A-D**). We found a clear defect in the HS-induction of up-regulated genes, consistent with a model in which HSF1 failed to adequately release Pol II after NELF depletion (**Fig. 6A**). The up-regulation defect was more prominent following the depletion of NELF-B than NELF-E (unpaired Mann-Whitney, p-value = 2.8e-4) (**Fig. 6B**), consistent with a larger impact of NELF-B depletion on paused Pol II. We also note that NELF depletion did not affect the HS-repression of down-regulated genes, indicating that HS-repression is independent of NELF.

**Figure 6:**
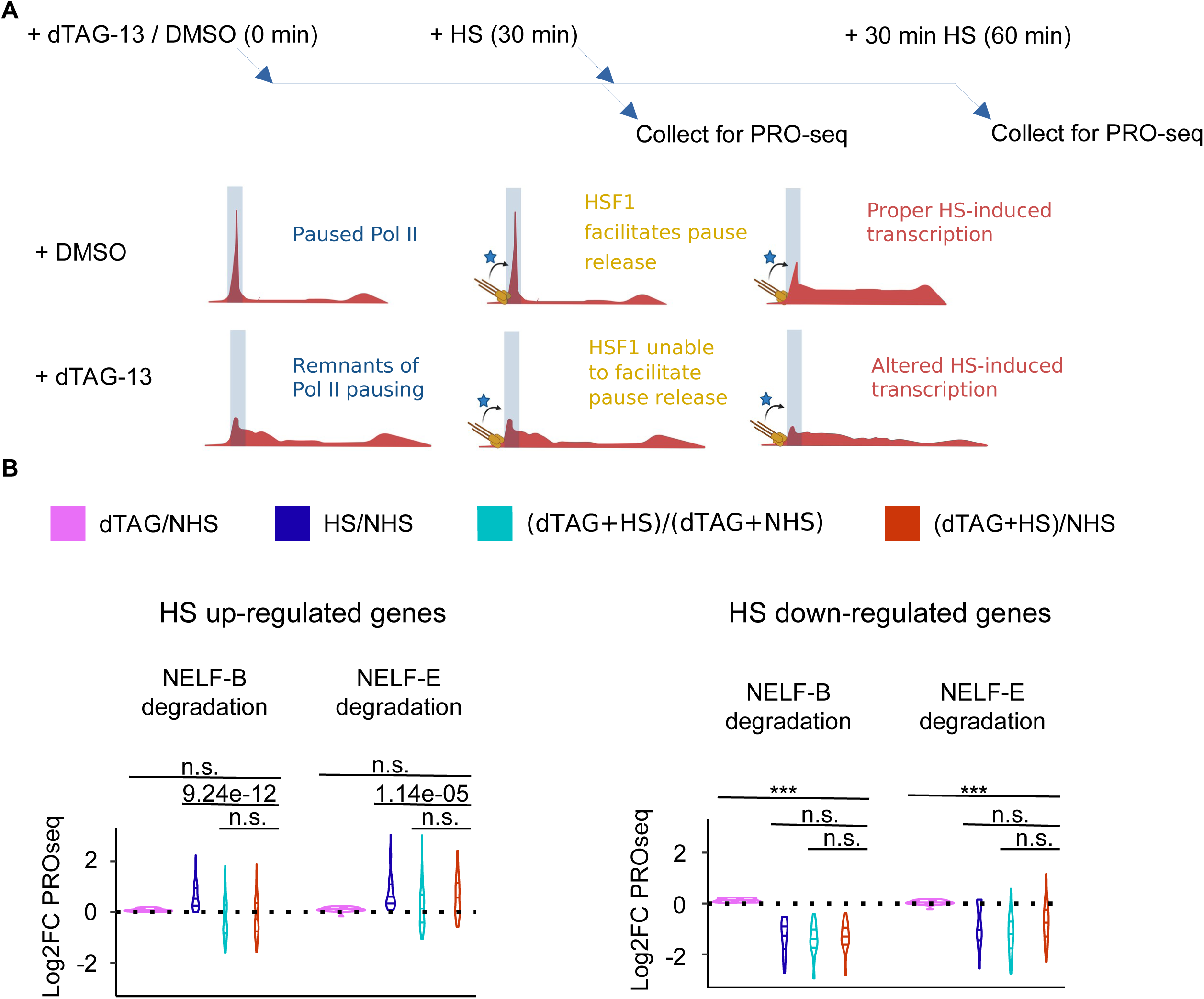
Removing paused Pol II prevents activation of genes by HSF1 after HS stimulation. (A) Time course of dTAG-13 drug treatment followed by heat shock (HS) (left) and cartoon depicting mechanisms of HSF1 action on Pol II pausing (right). HSF1 is depicted in yellow, while other cofactors that assist in pause release are depicted by the blue star. (B) Violin plots of log2 fold changes in PRO-seq signal in the gene-bodies (TSS +300, TES -300) for HS-upregulated (left) and downregulated (right) genes. A two-sided Mann-Whitney test was used to compute p-values, where n.s. Defines non-significant p-values, and (***) p-values < 2.2e-16.

Interestingly, many of the most highly HS-induced genes did not show a large defect in up-regulation as seen here (e.g. *Hspa1b*, *Hsp1h1*; **Supplemental Fig. 17**) and in a previous study^63,72^. The high rate of firing at these genes may be associated with the known high concentration of p-TEFb^73^, resulting in a higher probability of releasing Pol II in the right location before it creeps away from the promoter. In contrast, the more moderately induced and highly paused HS genes are firing less frequently and the creeping of paused Pol II to more downstream locations may prevent their proper activation (**Supplemental Fig. 18**). Alternatively, another interpretation is that highly induced HS genes unaffected by NELF depletion have a relatively short transcription unit, which could allow an incompletely modified and relatively inefficient elongation complex to nevertheless produce mRNA. Altogether, these findings support our model in which the evolution of pausing facilitated the ability of transcription factors to act on pause-release, providing an additional step to more tightly control gene expression.

## Discussion

Our work offers mechanistic insights into the evolution of gene regulation by promoter-proximal pausing from a preexisting step in the transcription cycle. We propose that the recruitment of P-TEFb, which causes pause-release in metazoans, actually serves a more general role that is necessary at all Pol II transcribed genes in all eukaryotic organisms, regardless of whether the organism or gene has a long-lived focal pause. The evolution of a “focal” pause collapsed the substrate for this step in transcription to a single location at each gene and increased the pause residence time. The degree to which Pol II slows down at the pause position, which progressively increased in unicellular Holozoan groups, appears to be affected first by the gain of a GC rich promoter proximal region which decreases the processivity of Pol II near the promoter and then by the evolution of NELFB-C/D, HEXIM, NELF-A, and, finally, NELF-E. This focal pause was necessary prior to the evolution of targeted regulation of pause release by transcription factors. The evolution of additional regulatory complexity may have helped to enable the evolution of complex multicellular organisms, which may explain how the loss of NELF, and in turn focal pausing, were tolerated in organisms which have lost multicellularity^8^ (yeasts) or developmental complexity (*C. elegans*).

As expected based on previous studies^17,63^, we found that depletion of NELF-B and NELF-E using the dTAG system alters pausing behavior directly. However, we note that many genes recover signal near the Pol II pause site, even in the absence of NELF-B. Why this recovery occurs remains unclear. Some of this appears to be explained by nucleosome positioning and/ or DNA sequence motifs associated with the pause. However, NELF-C and -A form a separate subcomplex and may retain some weak association with Pol II even in the absence of NELF-B. We also highlight the advent of a key tyrosine residue in HEXIM proteins, part of the 7SK complex which prevents pause release by P-TEFb^74^, as correlating with pausing index across eukaryotes. Collectively, our work supports models in which the NELF complex, alongside new innovations in the 7SK complex, coordinate to establish high residence time promoter-proximal paused Pol II characteristic of metazoan organisms. These innovations add structural features to the transcriptional regulatory nodes that control development in metazoans.

Our work shows that gain and loss of the NELF complex subunits correlates with the advent of a high Pol II residence time, consistent with classical work which argues that NELF is the critical factor responsible for promoter-proximal pausing^14,19^. However, one surprise in our work is that the impact of NELF-B and NELF-E degradation is small and Pol II is able to pause with relatively high residence times even after the prolonged degradation of NELF, consistent with other degron experiments^17,20^. Perhaps at the most stably paused genes Pol II retains its association with NELF so strongly and for so long^14,75,76^ that this population of NELF complexes are protected from degradation, or the remaining NELF subunits are sufficient for the retention of pausing at these subsets of genes. Altogether, the NELF complex on its own does not explain all of the variation in promoter-proximal pausing that we observe across eukaryotic organisms.

Recent studies have proposed that SPT6 plays a large role in pausing^20^. However, we found that SPT6 was present in nearly all eukaryotes regardless of whether they have paused Pol II (Supplemental Table 1). Moreover, we recently showed that Spt6 is only recruited after positive transcription elongation factor (P-TEFb)-mediated phosphorylation and RNA Pol II promoter-proximal pause escape^77^. Therefore we believe Spt6 is a poor candidate for contributing a pause. Instead, our work provides two alternative explanations: First, we show that HEXIM proteins in the 7SK complex have a distribution that also correlates with differences in the pausing index across species. The 7SK complex sequesters P-TEFb^78–80^ and therefore these proteins are excellent candidates to explain some of the variation in pausing index across different species, perhaps synergistically with new subunits in NELF. Second, we found that DNA sequence motifs proposed to influence pausing^49^ explain ∼25% of the variation in pausing index between species. Together, NELF, HEXIM, weak DNA sequence preferences, and the position of the +1 nucleosome are all factors that likely contribute to Pol II pausing. In conclusion, we postulate that NELF and HEXIM protein complexes are evolutionary innovations that create promoter-proximal Pol II pausing, a new transcriptional regulatory step which helped enable the evolution of complex developmental gene regulatory programs.

## Supporting information

Other supplementary files.

## Acknowledgments

We thank members of the Danko and Lis labs for valuable discussions and suggestions throughout the life of this project, and Meritxell Antó Subirats for preparing samples from *C. owczarzaki*, *C. fragrantissima* and *S. arctica*. We acknowledge the Fundación Pública Galega Centro Tecnolóxico de Supercomputación de Galicia (CESGA), Spain, for access to the FinisTerraeIII supercomputer, and Victoria Shabardina for facilitating access. Work in this publication was primarily supported by a grant from the NASA exobiology program (17-EXO-17-2-0112). Additional funding was also available from NHGRI (R01-HG010346 and R01-HG009309) to CGD, from the NIGMS (R01 GM147731) to ILB and CGD, and from NIH (RM1-GM139738) to JTL. AA was supported by the NIH (T32GM007739, F30HD103398). MML was supported by an Ayuda Juan de la Cierva-Incorporación postdoctoral fellowship (IJC2018-036657-I) from the Spanish Ministry of Science and Innovation. Work in AKH’s lab is supported by the NIH (R01HD094868, R01DK127821, R01HD086478, and P30CA008748). Work in IR-T was supported by an European Research Council Consolidator Grant (ERC-2012-Co -616960). The content is solely the responsibility of the authors and does not necessarily represent the official views of the US National Institutes of Health. Some of the figures in this manuscript were created using BioRender. Data was deposited in Gene Expression Omnibus (GSE223913).

## Author Contributions Statement

J.J.L., C.G.D., and A.G.C. designed the study. E.J.R. and A.A. performed experimental research. A.G.C., M.M.L., W.W., J.J.L., and C.G.D. designed and interpreted protein sequence comparisons across the tree of life. A.G.C., B.A.B., G.B., W.W., J.J.L., J.T.L., and C.G.D. analyzed and interpreted sequencing data. A.G.C., J.J.L., J.T.L., and C.G.D. wrote the manuscript. A.A., A.V., J.J.S., A.H.W., C.A.M., I.L.B, I.R.T, A.K.H., R.J.B. collected cells or provided samples for experimental research. All authors have been involved in revisions and approved the final manuscript.

## Methods

### Data Availability

Tables in CSV format can be downloaded from: https://github.com/alexachivu/PauseEvolution_prj

Data generated in this study can be found in Gene Expression Omnibus at: GSE223913.

### Code Availability

Custom code for analyzing sequencing data can be found on GitHub under: https://github.com/alexachivu/PauseEvolution_prj/

### Experimental methods

#### Sample collection

*E. coli:* An overnight culture of E. coli MG1655 was subcultured in 50 mL LB and grown at 37°C to OD 600 = 0.95. 5 mL aliquots were pelleted by centrifugation at 3000 × g. Pellets were permeabilized, washed, and flash-frozen as described in ^85^.

*H. mediterranei:* ATCC 33500 was grown for 48 hours at 35 °C in ATCC Medium 1176. 12.5 mL culture was centrifuged, and the cell pellet was resuspended in 3 mL cold non-yeast permeabilization buffer. To increase permeabilization of archaeal cells, the cell suspension was split into 3 × 1 mL aliquots in screw-cap tubes and combined with 400 µL sterile 0.5 mm glass beads. Cells were subject to bead-beating for 3 cycles of 2 minutes vortexing, 2 minutes on ice. Supernatants were transferred to 1.5 mL tubes, centrifuged to collect cell contents, and washed twice by resuspension in 500 µL storage buffer. Cells were resuspended in a final volume of 50 µL storage buffer and snap-frozen. The permeabilization and storage buffers were the same as reported previously ^85^, and include: ATCC Medium 1176 recipe (1 L), 156 g NaCl, 13 g MgCl 2 × 6H 2 O, 20 g MgSO 4 × 7H 2 O, 1 g CaCl 2 × 2H 2 O, 4 g KCl, 0.2 g NaHCO 3, 0.5 g NaBr, 5 g yeast extract, 1 g glucose. After mixing components, the pH was adjusted to 7.0 and the buffer was autoclaved.

*Sea Lampreys (Petromyzon marinus)* were obtained from Lake Michigan via the Great Lakes Fisheries Commission and maintained under University of Kentucky IACUC protocol number 2011-0848 (University of Kentucky Institutional Animal Care and Use Committee). For tissue sampling, animals were euthanized by immersion in buffered tricaine solution (1.0 g/l), dissected, and tissues were immediately frozen in liquid nitrogen. We analyzed muscle samples taken from the flank of one male and one female.

*Sea urchin (S.purpuratus):* All of the animal rearing and downstream processing use the same protocols as ^86^. Biological replicates of 20 hour blastula embryos were raised at 15°C in 0.2 um filtered sea water.

*D. iulia:* Wing tissues were sampled from Day 3 pupae derived from Costa Rican stock following standard protocols (e.g. ^87^ and ^30^). Wing tissues were dissected from pupae in cold PBS, after which nuclei were extracted in cold PBS using a dounce homogenizer. Nuclei were spun down and resuspended in nuclei storage buffer before flash freezing.

*Capsaspora owczarzaki* (strain ATCC 30864) was cultured axenically at 23°C in ATCC medium 1034 (modified PYNFH medium) in tissue culture-treated flat-bottomed polyethylene tissue culture flasks. Confluent cells were harvested by centrifugation (5000 x g, 5 minutes), and the pellet flash-frozen and stored at −80°C. For the isolation of intact nuclei, cells were harvested as before; the pellet was washed twice with phosphate-buffered saline (PBS), resuspended in 1ml of 2x Lysis Buffer (for 2x buffer: 10mM Tris-Cl pH 8.0; 300mM sucrose; 10mM NaCl; 2mM MgAc2;6mM CaCl2; 0.2% NP-40) and incubated on ice for 18 minutes. The resulting lysate was centrifuged (5000 x g, 5 minutes), and the pellet containing nuclei was washed once with 1ml Wash Buffer (10mM Tris-Cl pH 8.0; 300mM sucrose; 10mM NaCl; 2mM MgAc2). The nuclei were pelleted once more, resuspended in 1ml Storage Buffer (50mM Tris-Cl pH 8.0; 40% glycerol; 5mM CaCl2; 2 mM MgAc2), and stored at −80°C. For each buffer, 2 PhosStop^TM^ phosphatase inhibitor tablets (Roche), 1mM PMSF, 50 µg Pepstatin A, 56 mg sodium butyrate, and 1 cOmplete^TM^ Protease Inhibitor Cocktail tablet (Roche) per 50ml buffer were added immediately prior to use.

*Creolimax fragrantissima* and *Sphaeroforma arctica* were cultured axenically in BD Difco™ Marine Broth 2216, in tissue culture-treated flat-bottomed polyethylene tissue culture flasks at 12°C; confluent cells were harvested by centrifugation, and the pellet flash-frozen and stored at −80°C.

*Dictyostelium discoideum* AX3 wildtype cells were cultured axenically in HL-5 (Formedium) on untreated polystyrene petri dishes at 22°C. Confluent cells were resuspended in fresh media and centrifuged at 300x*g* for 5 min. The pellet was flash frozen and stored at −80°C.

*Nematostella vectensis:* Adult Nematostella were reared in 1/3 strength artificial seawater at 18°C in dark conditions. Spawning was induced using the protocol described in ^88^. Adult males and females were induced to spawn in small glass bowls, and fertilized egg masses were removed and cultured in small glass bowls at 25°C. Swimming gastrula/early polyp stage animals were harvested for nuclei isolation.

*Mouse embryonic stem cell (mESC) cell culture:* E14 mESCs (ATTC) were cultured on 0.1% gelatin-coated (Millipore) tissue culture-grade plates in a humidified 37°C incubator with 5% CO2. The culture medium consisted of DMEM (Gibco) supplemented with 2 mM L-glutamine (Gibco), 1× MEM nonessential amino acids (Gibco), 1 mM sodium pyruvate (Gibco), 100 U/mL penicillin/100 U/mL streptomycin (Gibco), 0.1 mM 2-mercaptoethanol (Gibco), 15% fetal bovine serum (Gibco), and 1000 U/mL recombinant leukemia inhibitory factor (LIF). Genetic editing and experiments were performed using cells at passages 10-20.

*Generation of NELFB and NELFE mESCs:* both cell lines were generated using an identical approach to endogenously and homozygously tag the C-terminus of each protein with FKBPF36V tag. The NELFB line has been previously described, and the NELFE line was generated for this study. The methods below describe the NELFE line generation, for more details of NELB line, please refer to ^17^.

*Plasmid Generation:* To target the *nelf-e* genes, two plasmid constructs were generated:

1. Cas9 vector to target the C terminus of Nelfe gene: PX459 vector (Addgene 62988) was digested using BbsI-HF (NEB) and single guide RNA targeting Nelfe was annealed (Ran et al. 2013)..
2. Homology-directed repair (HDR) vector containing the insert FKBPF36V tag, 2× HA tag, self-cleaving P2A sequence, and puromycin resistance, flanked by 1-kb Nelfe HDR sequences: The insert was obtained from pCRIS-PITCHv2-dTAG-Puro (Addgene 91796) (Nabet et al. 2018). The plasmid backbone (pBluescript), Nelfe HDR sequences, and the insert were amplified using Q5 polymerase (NEB), and the plasmid was constructed using NEBuilder HiFi DNA assembly (NEB). All oligos used are available in the table below.

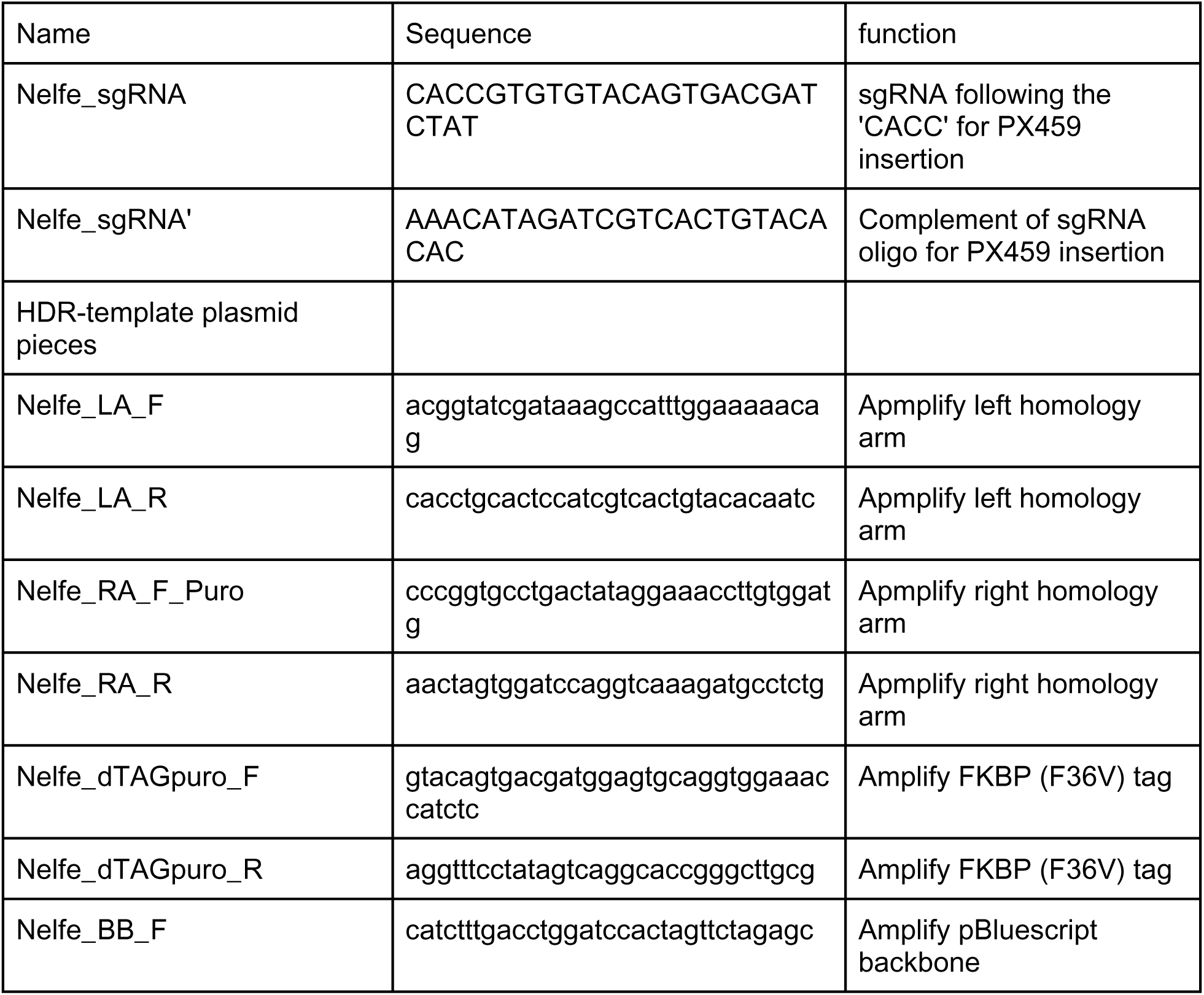
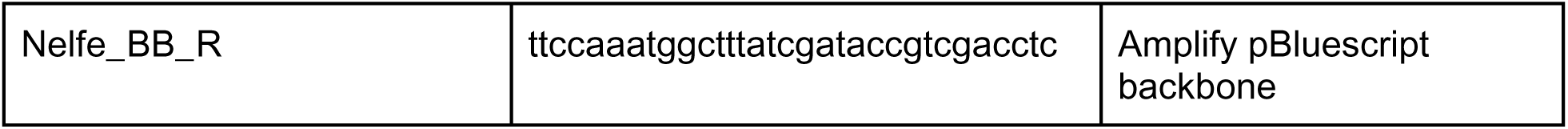

*Generation of Nelfe-dTAG mESCs:* 3 million cells were transfected with 10 µg of PX459-Nelfe_sgRNA and 10 µg of Nelfe_left-FKBPF36V-2xHA-P2A-PURO-Nelfe_right using the Lonza P3 primary cell 4D-Nucleofector X 100-µL cuvettes. The transfected cells were plated on a 10-cm dish coated with mouse embryonic fibroblasts (MEFs). 48 hours after transfection, correctly targeted cells were selected for in 6 µg/mL Blasticidin for 5 days. Surviving cells were split into 1000 cells per 10-cm dish and maintained for 9 days under puromycin selection. Surviving clones were picked and expanded under a stereomicroscope and genotyped for the insert.

*dTAG drug treatment:* The dTAG-13 reagent (Bio-Techne: https://www.bio-techne.com/p/small-molecules-peptides/dtag-13_6605) was reconstituted in DMSO (Sigma) to a final concentration of 5 mM. The dTAG-13 solution was diluted in culture medium to 500 nM and added to cells for the indicated time period during medium changes.

*Immunofluorescence:* Cells plated on u-Slide eight-well plates (Ibidi) were washed with PBS+/+ and fixed in 4% PFA (Electron Microscopy Sciences) in PBS+/+ for 10 min at room temperature. Cells were subsequently washed twice with PBS+/+, followed by wash buffer and 0.1% Triton X-100 (Sigma) in PBS+/+, and permeabilized in 0.5% Triton X-100 (Sigma) in PBS+/+ for 10 min. Then blocked with 3% donkey serum (Sigma) and 1% BSA (Sigma) for 1 h at room temperature. Cells were incubated with primary antibodies in blocking buffer overnight at 4°C (antibodies and concentrations are listed in Supplemental Table S1). Then, they were washed three times in wash buffer and incubated with suitable donkey Alexa Fluor (1:500; Invitrogen) for 1 h at room temperature. Finally, cells were washed three times with wash buffer, with the final wash containing 5 μg/mL Hoechst 33342 (Invitrogen), and imaged.

*Imaging:* Fixed immunostained samples were imaged using a Zeiss LSM880 laser scanning confocal microscope. An air plan-apochromat 20×/NA 0.75 objective was used. Images represent a 2D plane correlating to the monolayer of cells in culture. No further image processing was performed.

*Western blotting:* Cells were harvested and lysed by adding 350 µL of lysis buffer containing 1× cell lysis buffer (Cell Signaling), 1 mM PMSF (Cell Signaling), and cOmplete Ultra protease inhibitor (Sigma) to a 90% confluent six-well dish (Falcone) after washing with PBS−/−.

The harvested cells were incubated on ice for 5 min, scraped and collected then sonicated for 15 sec to complete lysis and then spun down at 12,000g for 10 min at 4°C. The supernatant was collected, and protein concentration was measured using Pierce BCA protein assay kit (Thermo). Samples were prepared by mixing 10 to 20 µg of protein with Blue loading buffer (Cell Signaling) and 40 mM DTT (Cell Signaling), followed by boiling for 5 min at 95°C for denaturation. Cellular compartment fractions were prepared using subcellular protein fractionation kit (Thermo) following the manufacturer’s instructions.

The samples were run on a Bio-Rad Protean system and transferred to a nitrocellulose membrane (Cell Signaling) using transblot semidry transfer cells (Bio-Rad) following the manufacturer’s instructions and reagents. The nitrocellulose membrane was briefly washed with ddH2O, stained with Ponceau S (Sigma) for 1 min, and washed three times with TBST (0.1% Tween 20 [Fisher] in TBS) to check for transfer quality and serve as a loading control. Then it got blocked with 4% BSA in TBST for 1 h at room temperature and incubated with primary antibodies diluted in blocking buffer overnight at 4°C. The membrane was then washed three times with TBST, incubated with secondary antibodies in blocking buffer for 1 h, and washed three times with TBST. Last, the nitrocellulose membrane was incubated with ECL reagent SignalFire for 1-2 min and imaged using a ChemiDoc (Bio-Rad).

The following antibodies were used in this paper:

| Antibody | Source | Identifier | Application | Conc. |
| --- | --- | --- | --- | --- |
| anti-Histone H3 | Cell Signaling | Cat# 4499, RRID: AB_10544537 | Western | 1:2000 |
| anti-RNA pol II S2P | Abcam | Cat# ab193468, RRID: AB_2905557 | IF | 1:500 |
| anti-NELFE | Abcam | Cat# ab170104, RRID: AB_2827280 | Western | 1:1000 |
| anti-COBRA1/NELFB | Cell Signaling | Cat# 14894, RRID: AB_2798637 | Western | 1:1000 |
| anti-HA | Abcam | Cat# ab130275, RRID: AB_11156884 | IF | 1:500 |
| anti-HA | Cell Signaling | Cat# 3724, RRID: AB_1549585 | Western | 1:1000 |
| anti-b-actin | Cell Signaling | Cat# 3700, RRID: AB_2242334 | Western | 1:2000 |
| anti-DSIF/Spt5 | BD Biosciences | Cat# 611106, RRID: AB_398420 | Western | 1:2000 |

*Heat shock experiments on mESCs:* Heat shock was administered as described in recent work from the Lis lab ^67,71^. We started the heat stress after 30 min of dTAG-13 treatment, which corresponds to the maximal depletion of paused Pol II based on PRO-seq data.

We performed the analysis of the HS data in two different ways:

- On a first analysis, by calling gene expression changes using DEseq (log2foldChange > or < 0, and padj < 0.05) between a regular HS and NHS experiment. Then, we plotted log2 fold changes of (HS+dTAG)/(NH) and (HS+dTAG)/(NHS+dTAG) at these pre-defined HS up-regulated or down-regulated genes coordinates.
- For a second approach, we considered the effect that NELF-B depletion has on Pol II trickling into gene bodies. To eliminate any biases from increased PRO-seq signal downstream from the TSS due to the dTAG-13 treatment alone, we re-analyzed our data after removing the first 3 kb downstream from the TSS and we focused on genes that remain unchanged following dTAG-13 treatment, but are either up or down-regulated after HS (**fig. S18F**). We confirmed a slight up-regulation of genes in the vicinity of the TSS after dTAG-13 treatment by plotting the correlation between the log2 fold change of dTAG-13 treatment after NELF-B degradation (**fig. S16**). We used deeptools to compute log2 fold change bigwigs and plot heat maps.

Both of these analyses confirmed a defect in up-regulation across many HS-dependent genes. In the second analysis, we observed that a significant number of genes that were meant to show HS-dependent upregulation failed to reach their full transcription potential in the absence of NELF-B (**Fig. 4C left; fig. S18A**). The same effect was also observed, though in far fewer genes in the absence of NELF-E (**Fig. 4C right; fig. S18B**). We noted no defect in down-regulated genes using this analysis approach.

*PRO-seq library prep:* PRO-seq or ChRO-seq ^25,89^ libraries were prepared from snap-frozen cell pellets following the protocol described in ^85^. All PRO-seq libraries were evaluated for data quality and sequencing depth using PEPPRO ^90^. Data and data quality are shown in (**fig. S1**).

### Computational analyses

*\*In this paper we refer to PRO-seq, GRO-seq, and ChRO-seq as **PRO-seq**. Mapping and processing PRO-seq data:*

Single and paired-end reads of PRO-seq data were aligned to its reference genome using the proseq2.0 pipeline from the Danko lab (https://github.com/Danko-Lab/proseq2.0) using the following parameters: -RNA5=R1_5prime --RNA3=R2_5prime --ADAPT1=GATCGTCGGACTGTAGAACTCTGAACG --ADAPT2=AGATCGGAAGAGCACACGTCTGAACTC --UMI1=4 --UMI2=4 --ADD_B1=6 --ADD_B2=0 --thread=8 --map5=FALSE. Library processing included adapter trimming using cutadapt, PCR deduplication (where UMIs are present) using printseq-lite.pl, followed by mapping to the reference genome using BWA. Mapped BAM files were then trimmed either to the 3’-end of the RNA (to map the location of RNA Pol II) or the 5’-end (to map the beginning of the RNA) and the 1bp position was converted to bedGraphs and BigWigs. PRO-seq libraries were also RPM normalized to account for differences in sequencing depth.

Data was mapped to the following reference genomes:

- E.coli:escherichia_coli_mg1655_01312020 (https://www.ncbi.nlm.nih.gov/nuccore/U00096.2)
- Haloferax: NZ_CP039139.1 (https://www.ncbi.nlm.nih.gov/nuccore/NZ_CP039139.1) and the plasmids included here: https://www.ncbi.nlm.nih.gov/genome/?term=txid523841
- D.discoideum: dicty_2.7 (https://www.ebi.ac.uk/ena/browser/view/GCA_000004695.1)
- A.thaliana: Arabidopsis_thaliana.TAIR10.dna.toplevel
- Z.mays:GCF_902167145.1_Zm-B73-REFERENCE-NAM-5.0 (https://www.ncbi.nlm.nih.gov/assembly/GCF_902167145.1/)
- O.sativa:GCF_001433935.1_IRGSP-1.0_genomic.fna (https://www.ncbi.nlm.nih.gov/assembly/GCF_001433935.1/)
- C.owczarzaki: Capsaspora_owczarzaki_atcc_30864.C_owczarzaki_V2.dna.toplevel
- S.pombe: Schizosaccharomyces_pombe.ASM294v2.dna.toplevel
- S.cerevisiae: Saccharomyces_cerevisiae.R64-1-1.dna.toplevel
- S.arctica: Sphaeroforma_arctica_jp610.Spha_arctica_JP610_V1.dna.toplevel.fa.gz (https://www.ncbi.nlm.nih.gov/assembly/GCF_001186125.1/)
- C.fragmatissima: Creolimax_fragrantissima.genome (ncbi.nlm.nih.gov/assembly/GCA_002024145.1/)
- N.vectensis: nemVec1
- C.elegans: ce6 (https://hgdownload.soe.ucsc.edu/goldenPath/ce6/chromosomes/)
- D.pulex: dpulex_jgi060905 (http://wfleabase.org/prerelease/dpulex_jgi060905/genome-assembly/)
- D.iulia: published assembly in ^87^
- D.melanogaster:dm3 (http://genome.ucsc.edu/cgi-bin/hgTracks?db=dm3&chromInfoPage=)
- S.purpuratus: Spur_5.0 (https://www.ncbi.nlm.nih.gov/assembly/GCF_000002235.5/)
- P.marinus: petMar2 (https://www.ncbi.nlm.nih.gov/assembly/GCA_000148955.1/)
- M.musculus: mm10 (GRCm38)
- H.sapiens: hg19 (https://www.ncbi.nlm.nih.gov/assembly/GCF_000001405.13/)

#### Reannotation of transcription start sites

We took PRO-seq mapped BAM files and ran it through the RunOnBamToBigWig tool developed in the Danko lab (https://github.com/Danko-Lab/RunOnBamToBigWig) to compute 5’prime mapped BigWigs (parameter for paired end data: --RNA5=R1_5prime; parameter for single end data: --SE_READ=RNA_5prime), or, for samples which used alternate sequencing chemistries, directly imported 5’ mapped ends of BAM files using the BRGenomics R package v1.10.0^91^

(https://github.com/alexachivu/PauseEvolution_prj/blob/main/Reannotate_TSSs). All following analyses were performed in R v4.2.1^92^, where range transformations were performed using rtracklayer v1.58.0^93^ and plyranges v1.16.0^94^ with BSgenome v1.66.1^95^ and tidyverse v1.3.2^96^. We used published gene annotations in each species, resized them to a 500bp window centered on the gene annotation start site, and computed the total number of 5’-prime mapped PRO-seq reads that fall within this interval using 10bp windows. With these signal counts, we performed a Wilcoxon test between two adjacent sliding windows of 20 counts, and selected the 10 base window which most significantly (lowest p-value) separated the two distributions of counts, conditioned such that windows to the left (upstream) of this point must have a lower median count than those to the right (downstream). Last, to reannotate gene start sites at single-base resolution, we took the maximum single-base 5’-PRO-seq signal within that 10bp window. We used these annotated TSSs for all further analyses. These operations were parallelized using the parallel (v4.2.1)^92^ and zoo (v1.8.11)^97^ R packages to decrease runtime. Only expressed genes with a length of at least 750bp were considered for reannotation. Exclusion criteria and search window size were selected to preserve the most possible annotations in all species. Some alternative TSS were collapsed into one by reannotation. In cases where this occurred we eliminated the duplicate genes before downstream analysis. The trustworthiness of these new annotations were assessed qualitatively in species where PRO-cap datasets are available (Supplemental Figure 15).

#### Computing Pausing indexes and generating PRO-seq metaprofile plots

Pausing indexes were computed as the ratio between Pol II density in the pause region and gene body region. We defined the pause region as the interval between [ TSS, TSS+150 ] bp and the gene bodies as [ TSS+200, TES ] bp (where TES = annotated transcription end site). Genes shorter than 400bp after reannotation were removed from this analysis. Bootstrapped metaprofiles including a 50% confidence interval around the mean PRO-seq signal were produced using the BRGenomics R package v1.10.0^91^ and ggplot2 v3.4.4^98^.

#### Heatmaps and metaprofiles for NELF degradation panels

We use DeepTools to functions (bigwigCompare, compute matrix, plotHeatmap, and plotProfile) to compute heatmaps and meta profiles of PRO-seq data. We also used DeepTools to cluster and compute correlations between the heat shock PRO-seq libraries as BAM files (bamCompare, and multiBamSummary, plotPCA, and plotCorrelation) ^99^.

Before running bigwigCompare, we generated combined BigWigs of the plus and minus PRO-seq data for each sample.

**bigwigCompare** --bigwig1 HS_plus.minus.bw --bigwig2 NHS_plus.minus.bw -o HS.NHS_log2FC_plus.minus.bw --outFileFormat bigwig --pseudocount 0.1 --operation log2 --skipNAs -p max/2 &

**computeMatrix** reference-point --referencePoint center -R regions.bed -S NHS_plus.minus.bw HS.NHS_log2FC_plus.minus.bw --samplesLabel “NHS” “HS / NHS” --sortUsingSamples 1 \ - b 1000 -a 50000 \ - -binSize 1000 \ - -skipZeros -o Counts_GB.increase_log2FCs_NELF_B.gz -p max/2

**plotHeatmap** -m Counts_GB.increase_log2FCs_NELF_B.gz \ - out Heatmap_Counts_GB.increase_log2FCs_NELF_B.pdf \ - -colorList “blue,white,red” \ - -heatmapHeight 10 \ - -heatmapWidth 3 \ - -zMin -2 --zMax 2 --missingDataColor 0 \ - -averageTypeSummaryPlot “mean”

**plotProfile** -m Counts_GB.increase_log2FCs_NELF_B.gz \ - out Metaplot_Counts_GB.increase_log2FCs_NELF_B.pdf

**multiBamSummary** bins \ - -bamfiles ./*bam \ - -minMappingQuality 10 \ - p max/2 \ - out allTSS_QC_readCount_corr.npz \ - -outRawCounts allTSS_QC_readCount_corrRaw.tab

**plotCorrelation** \
- in allTSS_Abood.PROseq_readCount_corr.npz \
- -corMethod spearman --skipZeros \
- -plotTitle “Spearman Correlation at annotated TSSs” \
- -whatToPlot heatmap --colorMap seismic --plotNumbers \
- o allTSS_QC.Spearman.heatmap_readCounts.png \
- -outFileCorMatrix allTSS_QC.PROseq_readCount_Spearman.tab

#### Identification of transcriptional machinery components

We first queried taxonomically broad databases of eukaryotic, archaeal and bacterial amino acid sequences using BLASTP with human queries (NELFA: NP_005654.4 ; NELFB: NP_056271.3 ; NELFC/D: NP_945327.3 ; NELFE: P18615.3 ; HEXIM proteins: NP_006451.1). For eukaryotic sequences, we queried a custom database consisting of the predicted proteome sequences in EukProt v.3^100^ from which most animal sequences had been removed, leaving a taxonomically representative subsection of animals (for a list of datasets included, see **Supplemental Table 1**). For archaeal and bacterial sequences, we queried the Genome Taxonomy Database^101,102^ release 207. Reciprocal best blast searches were conducted against the human predicted proteome (from EukProt v3, originally from the human genome assembly GRCh38.p12), and Pfam domains were predicted using the PfamScan tool with Pfam v.35^103^. Additional searches were conducted against databases containing sequences from organisms in which PRO-Seq studies were carried out (see Supplementary Methods). Sequences with reciprocal best blast hits to human proteins of interest were retrieved and clustered using CD-HIT 4.8.1^104,105^), and either used to construct phylogenies, or aligned with MAFFT L-INS-i 7.475^106–108^, and used to construct profile Hidden Markov Models (HMMs) with HMMer 3.3.2^109^with HMMer’s hmmsearch program (per-target e-value cut-off 0.01 and per-domain e-value cut-off 0.3).

For preliminary phylogenies, sequences were aligned using MAFFT v. 7.475 in automated mode and trimmed using trimAl v1.4.rev22^110^ in *gappyout* mode, and maximum likelihood trees were constructed in IQ-TREE 2.3.1 using models chosen by the automated ModelFinder algorithm. For final phylogenies, short or poorly aligned sequences were removed; sequences were aligned using MAFFT L-INS-i and trimmed using trimAl in *gappyout* mode. Maximum likelihood phylogenies were constructed in IQ-TREE using the LG substitution matrix, the C20 mixture model with weights optimised to each dataset (--mix-opt option), and gamma rate heterogeneity among sites.

For detailed search strategies used for each component, see Supplementary Methods.

Fasta files containing sequences identified as probable homologs of human transcriptional machinery components of interest, all HMMs and the alignments used to construct them, untrimmed and trimmed alignments used to construct phylogenies (Supplemental Fig. 5), and the phylogenies themselves in Newick format, are provided as Supplementary Datasets.

#### Defining DNA sequence motif under the pause

The position of the pause site in each species was defined as follows. First, we utilized our re-annotated TSS positions and created a 100bp window starting from the TSS: [ TSS, TSS+100 ] bp for the pause button motif or a 100bp window centered on the TSS for the initiator motif.

For motif enrichment analysis, we searched this window for the maximum match to the human consensus motif. We calculated the motif match score over each motif sized window in this pause region by averaging the prevalence of the recovered base at each position in the consensus motif. The maximum match score was normalized to the average match score over a 1kb window assigned randomly 1kb upstream or 1kb downstream of the target 100bp window. We chose to use the maximum match score in the target window because we expect to recover each motif exactly one time in proximity to any given TSS. We used the average match score over the background window to prevent spurious similarity to a low-information motif and to prevent recovery of genuine motifs at nearby alternative TSSs from affecting match score comparability between distantly related species.

For motif discovery, we searched the same 100bp window downstream of the TSS for the maximum 3’ PRO-seq signal and created a 17bp window around this base. We extracted the underlying DNA sequence in that window from the reference genome and created motif logos using Biostrings v2.64.1^111^ and the seqLogo R package v1.68.0^112^. Given that this analysis presumes the presence of a PRO-seq maximum in the pause region, it was not performed in species where a pause or proto-pause is not observable.

#### Differential analysis

We performed differential analysis to quantify changes after heat shock, dTAG-13, and the dual treatment of heat shock and dTAG-13. To accomplish this, we run DEseq2. We used the total number of dm3 spike-in reads (divided by the mean of the spike-ins) as scaling factors.

#### Figure design

We used BioRender to draw all of the illustrations and cartoons in this paper, with the exception of the schematic representing the relationships between species. The latter was prepared using Interactive Tree of Life (iTOL) v.6.7^113^ based on relationships depicted in^114^ and edited in InkScape and Illustrator. PRO-seq metaprofile plots and all pausing index box plots and regressions were generated using ggplot2 v3.4.4^98^ with ggsignif v0.6.4^115^ and RColorBrewer v1.1.3^116^ and were arranged using cowpolot v1.1.1^117^, InkScape, and Illustrator.

*An inventory of all functions used to process the PRO-seq data is deposited on GitHub at:*

https://github.com/alexachivu/PauseEvolution_prj/tree/main

## Supplemental Materials

**Figure S1:**
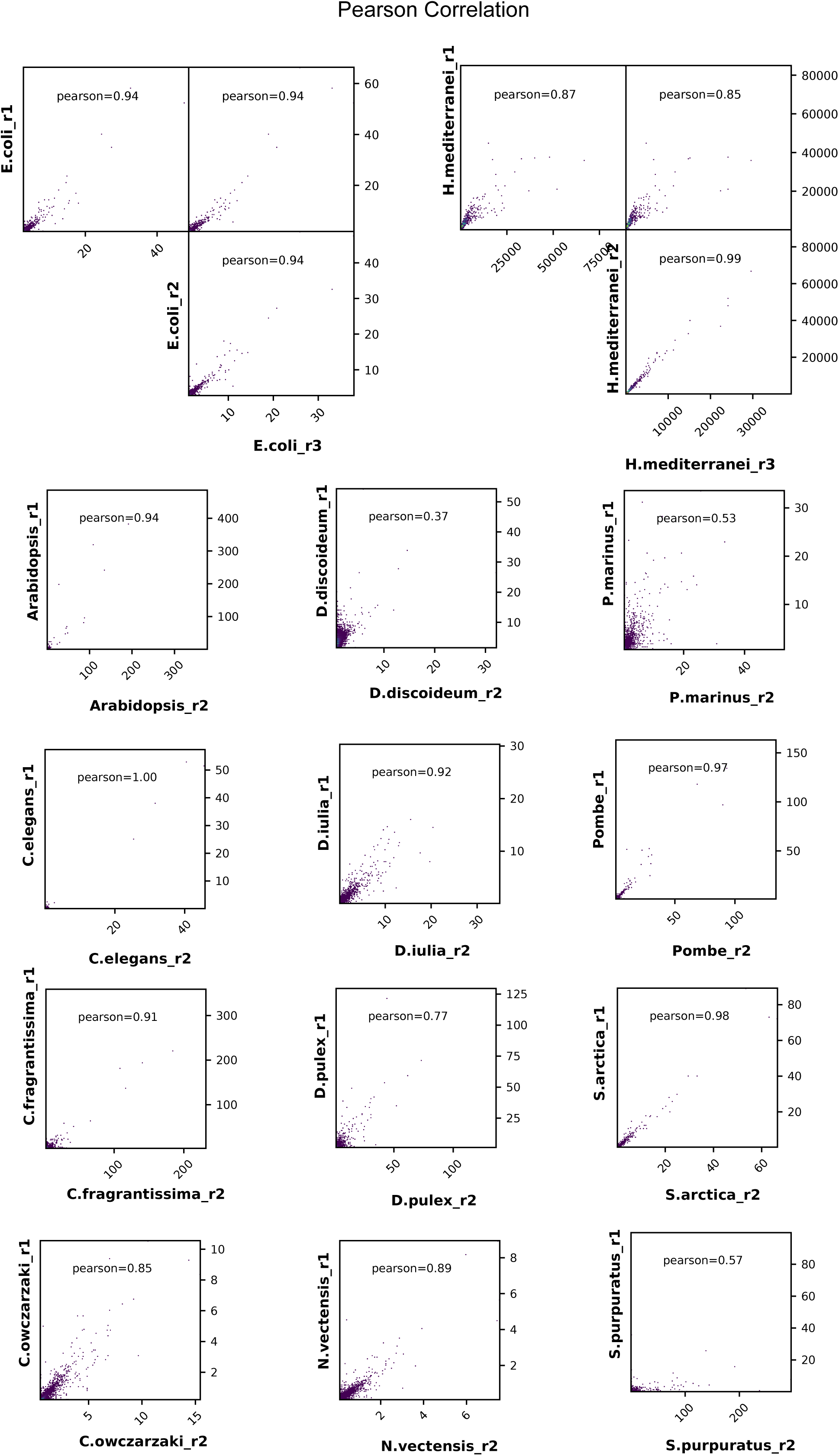
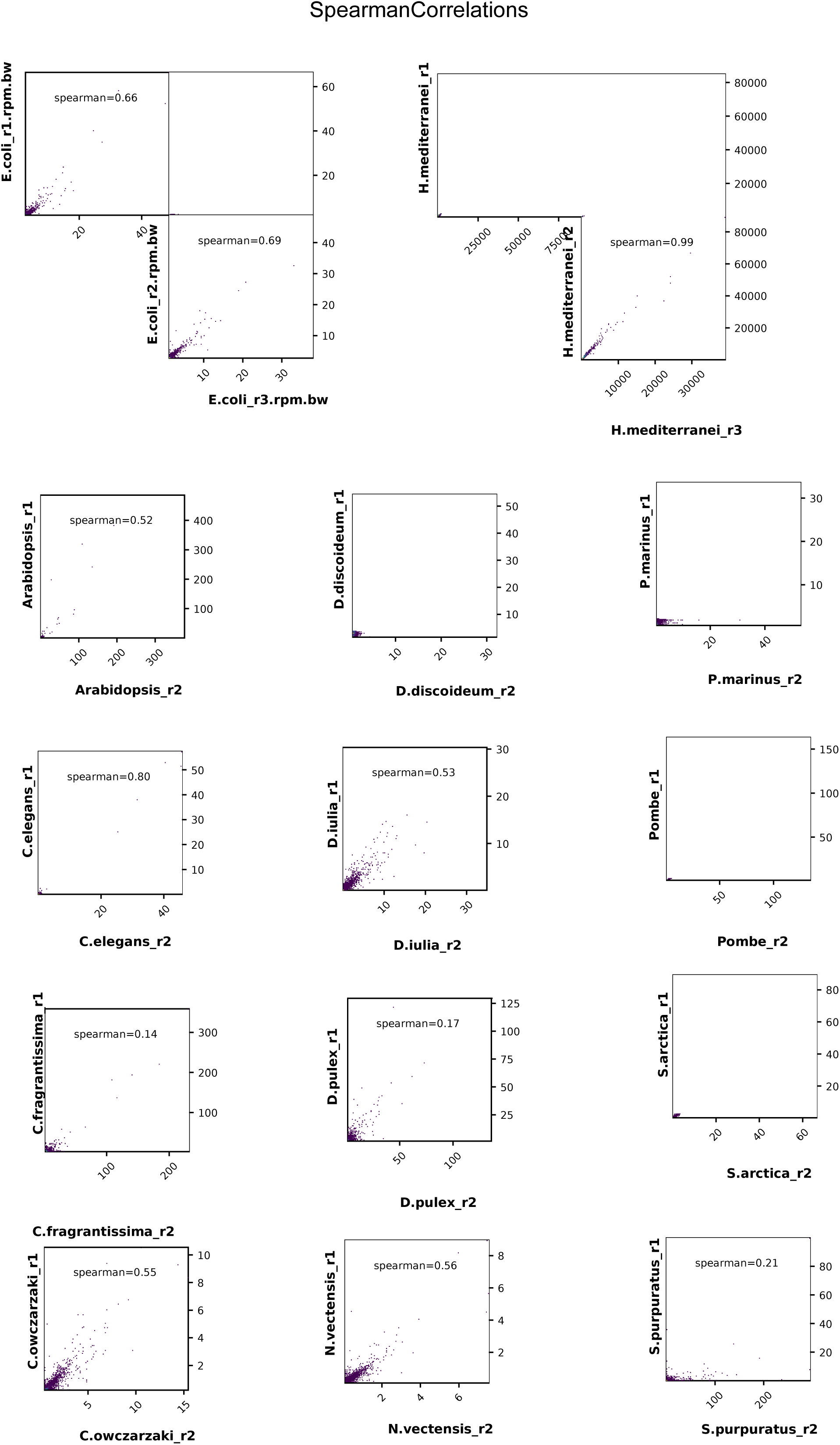
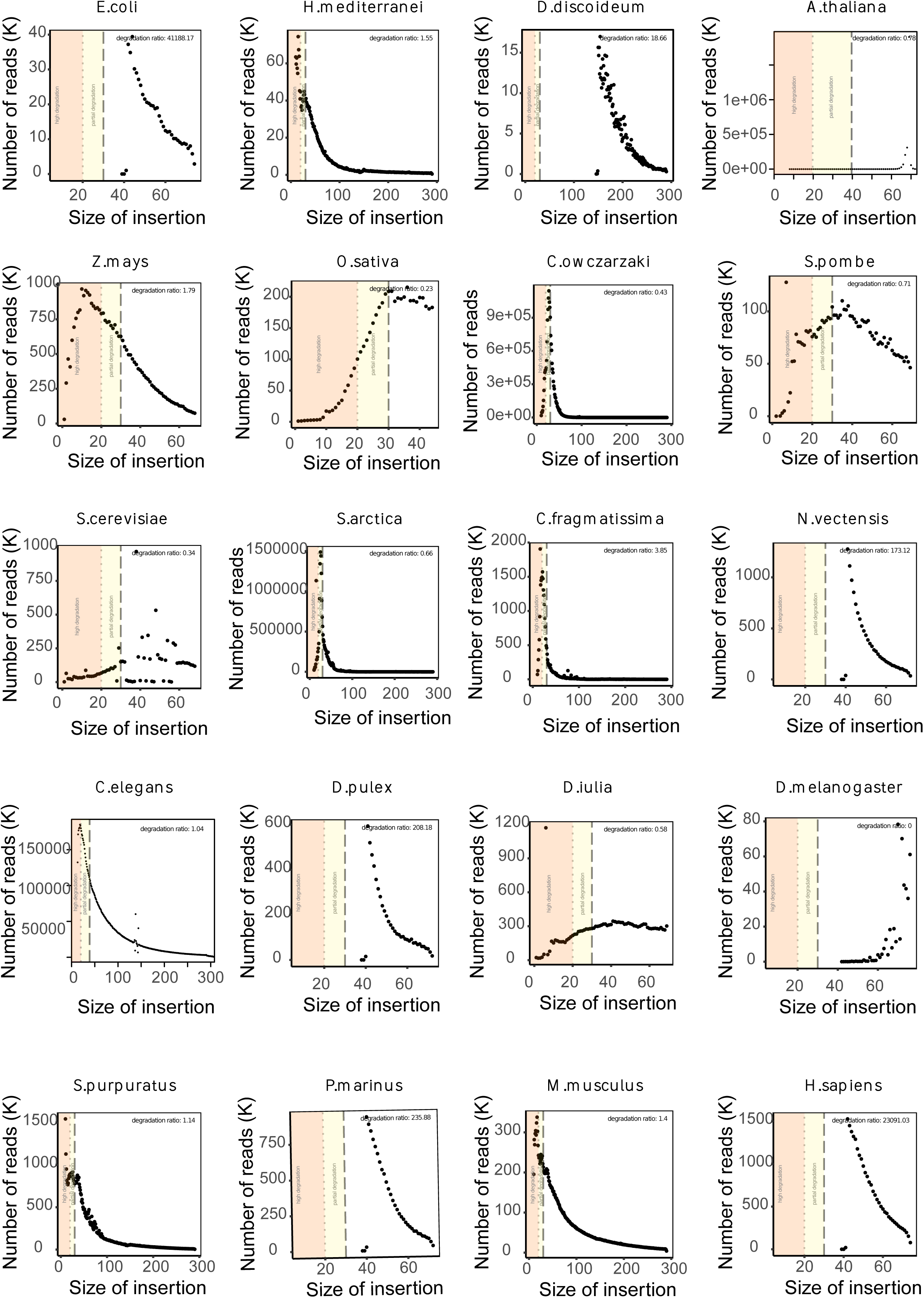
Correlation of experimental replicates. Pearson (A) and Spearman (B) correlation between biological replicates for ChRO-seq data generated for this project. In some cases biological replicates came from different tissues (D. iulia), sexes (P. marinus), or treatment conditions (D. discoideum, S. purpuratus). (C) Profiles show the number of PRO-seq reads per species is reported as a function of insert size. A color gradient from orange (depicting highly degraded RNA) to white (depicting lowly degraded RNA) marks the quality of each sample. A degradation ratio score is also reported at the top of each plot. Degradation ratios for *C.elegans* and *A.thaliana* were computed manually using the scripts in Scott *et al.*^83^.

**Figure S2:**
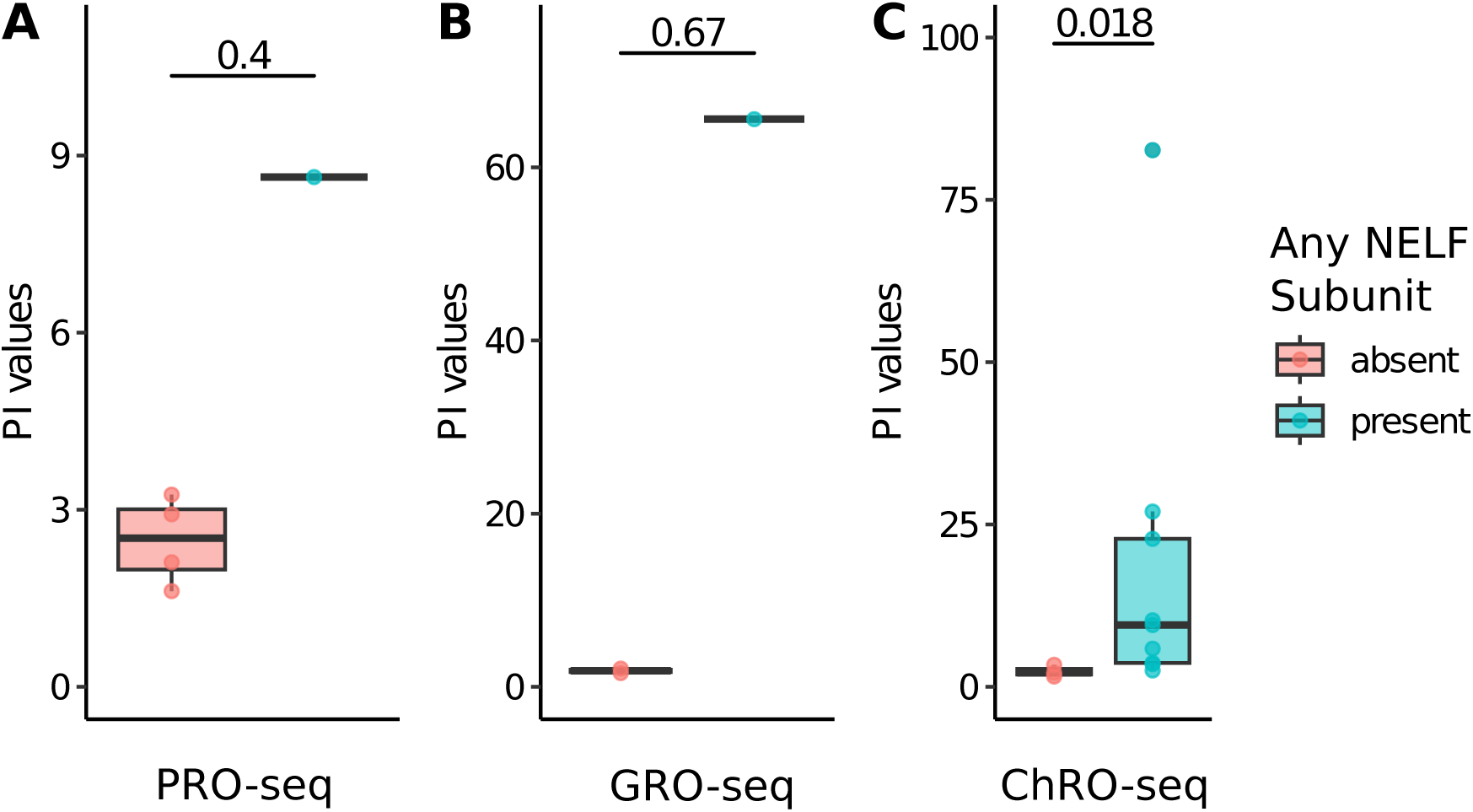
Comparability of PRO-seq, GRO-seq, and ChRO-seq. Box and whiskers plot of pausing indexes in each species. Boxes are clustered by whether any NELF subunits are present (at least one subunit present vs. no subunits present) in each species for which PRO-seq (A, n = 5), GRO-seq (B, n = 3), and ChRO-seq (C, n = 12), were used in this project. A Mann-Whitney test was used to compute p-values.

**Figure S3:**
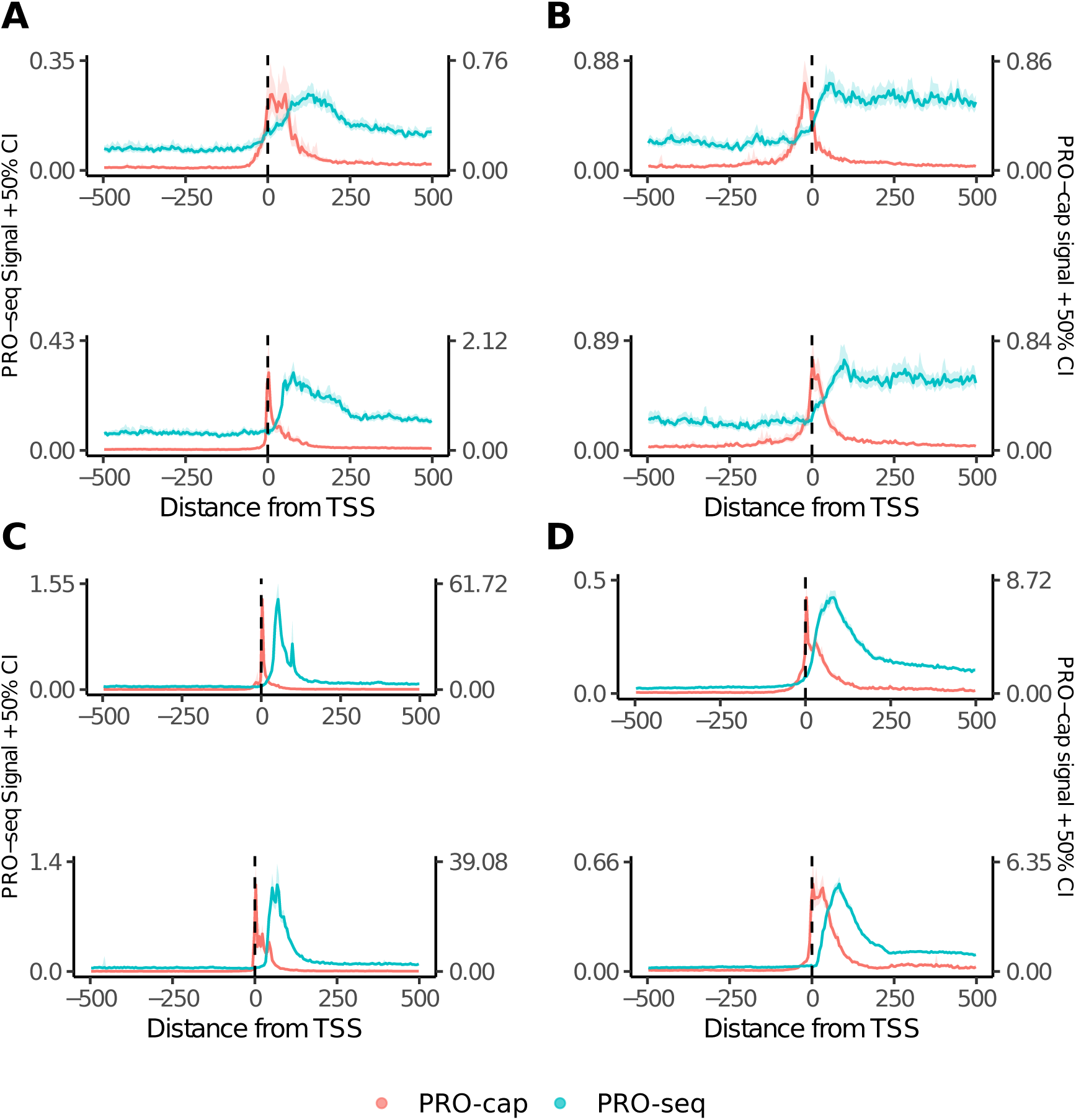
TSS reannotation. PRO-seq (blue) and PRO-cap^25^ (red) metaprofiles for *S. Pombe*^23^ (A), *S. cerevisiae*^23^ (B), *D. Melanogaster*^25^ (C), and *H. Sapiens*^84^ (D). Upper panels use published gene annotations, lower panels use reannotated genes. Note the relative depletion of PRO-cap reads upstream and downstream of the TSS and more focused pause in PRO-seq signal of *D. melanogaster* and *H. sapiens* in re annotated panels.

**Figure S4:**
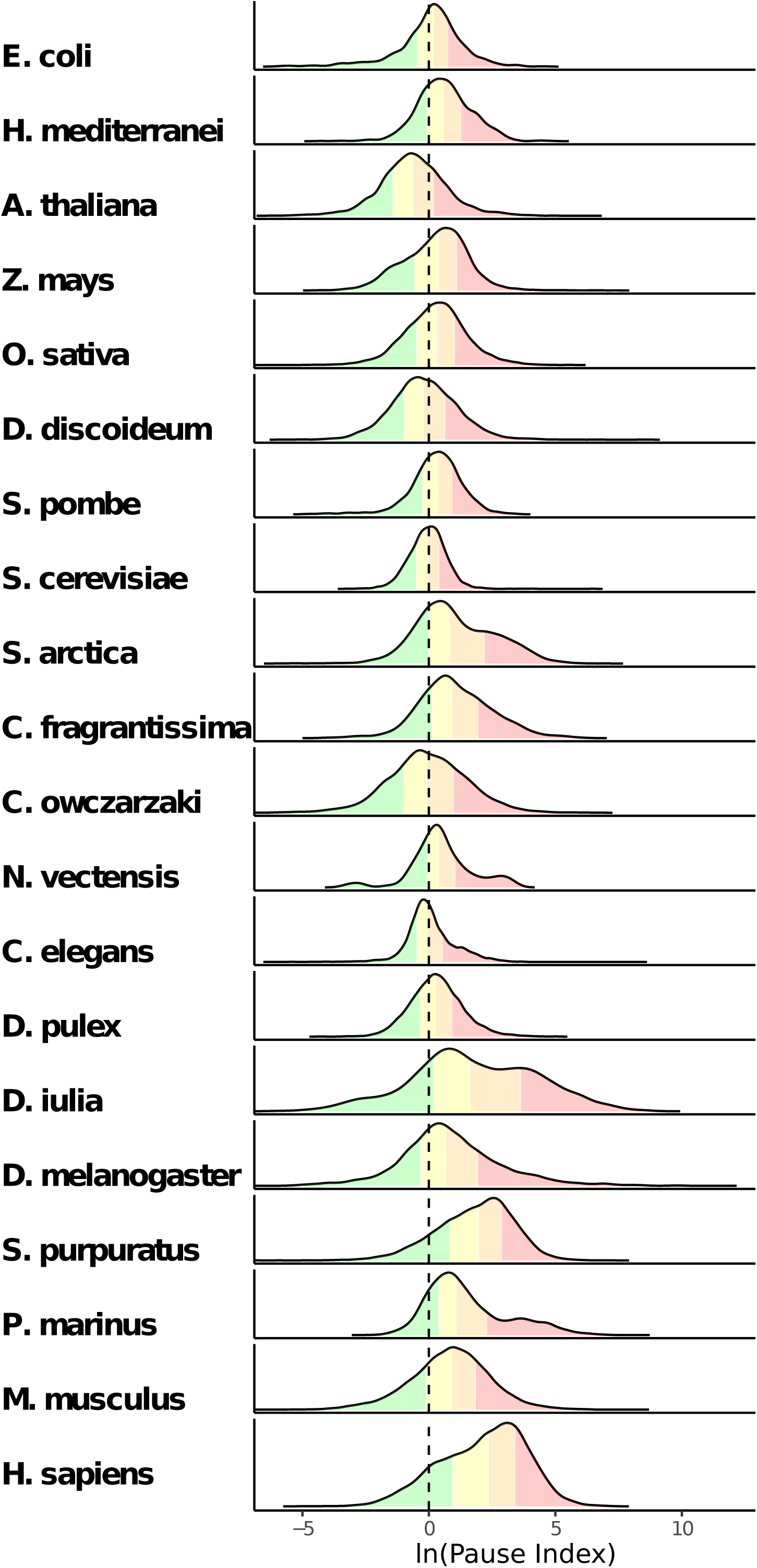
Clustering species by their pausing index values. Density maps of pausing indexes for each species. The plots are split into four quantiles and colored accordingly.

**Figure S5:** Maximum likelihood phylogenies of pausing machinery components. Red branches: opisthokonts, purple branches: amorpheans (opisthokonts and their sister lineages breviates, apusomonads and amoebozoans), cyan branches: other eukaryotes, black branches: archaea, grey branches: bacteria. For ease of viewing, only taxon names of opisthokonts, and only robust support values (higher than both 95% ultrafast bootstrap and 80% SH-aLRT supports), are shown. Raw phylogenies are provided as supplementary materials [Supplementary Files].

**Figure S6:**
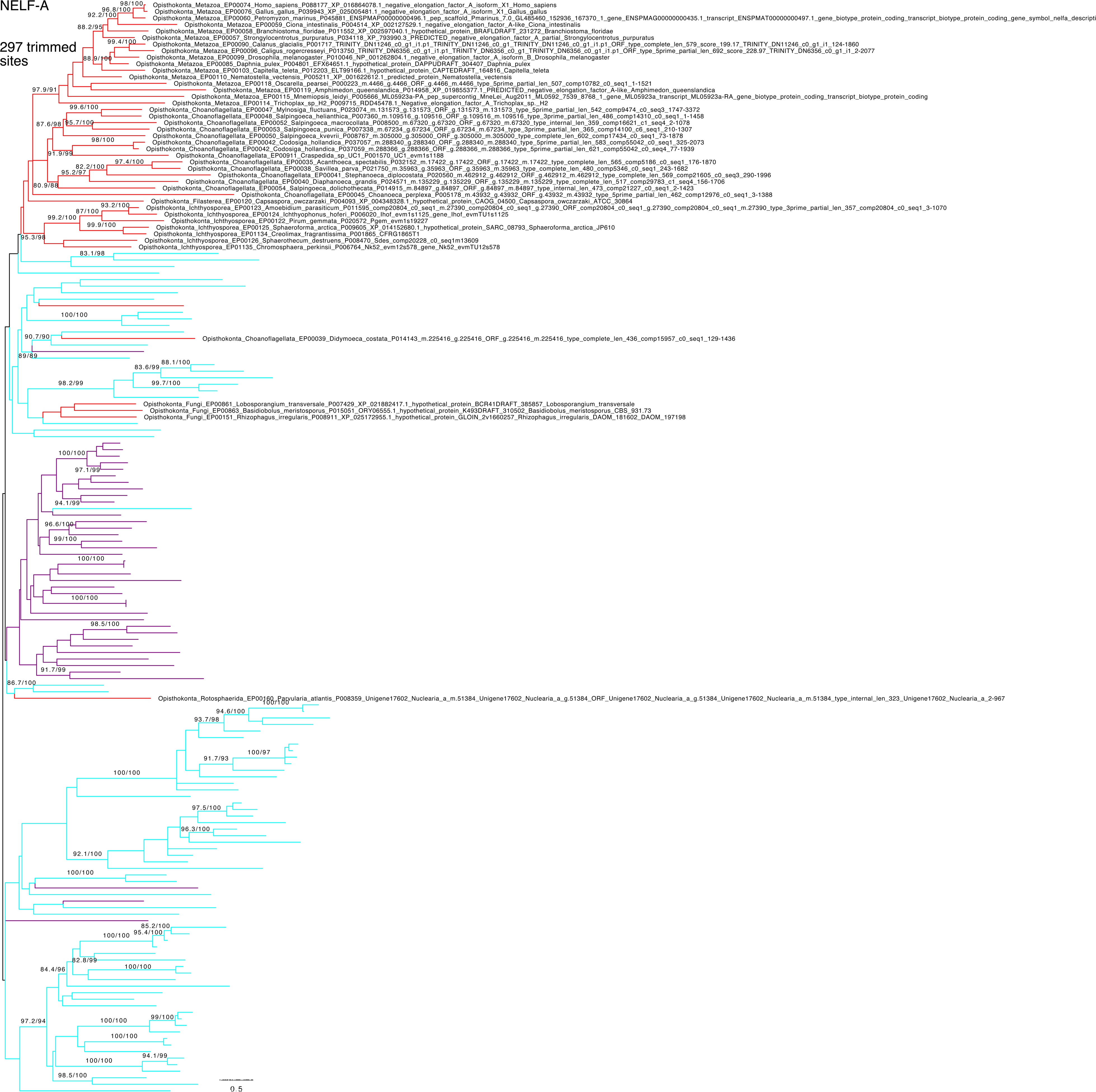
Selected positions of a multiple sequence alignment of NELF-A sequences from selected eukaryotes. Amino acids are shaded according to hydrophobicity value, where red is most hydrophobic and blue is the most hydrophilic, and numbered according to the Homo sapiens sequence. Locations of the Homo sapiens NELF-A NELFCD-binding domain (black) and NELF-A tentacle (grey) are shown above the sequence. HDAg domains predicted by InterProScan are underlined. Alignment constructed with MAFFT-L-INS-I, visualized with Geneious® 2023.2.1, and edited with Affinity Designer 2.

**Figure S7:**
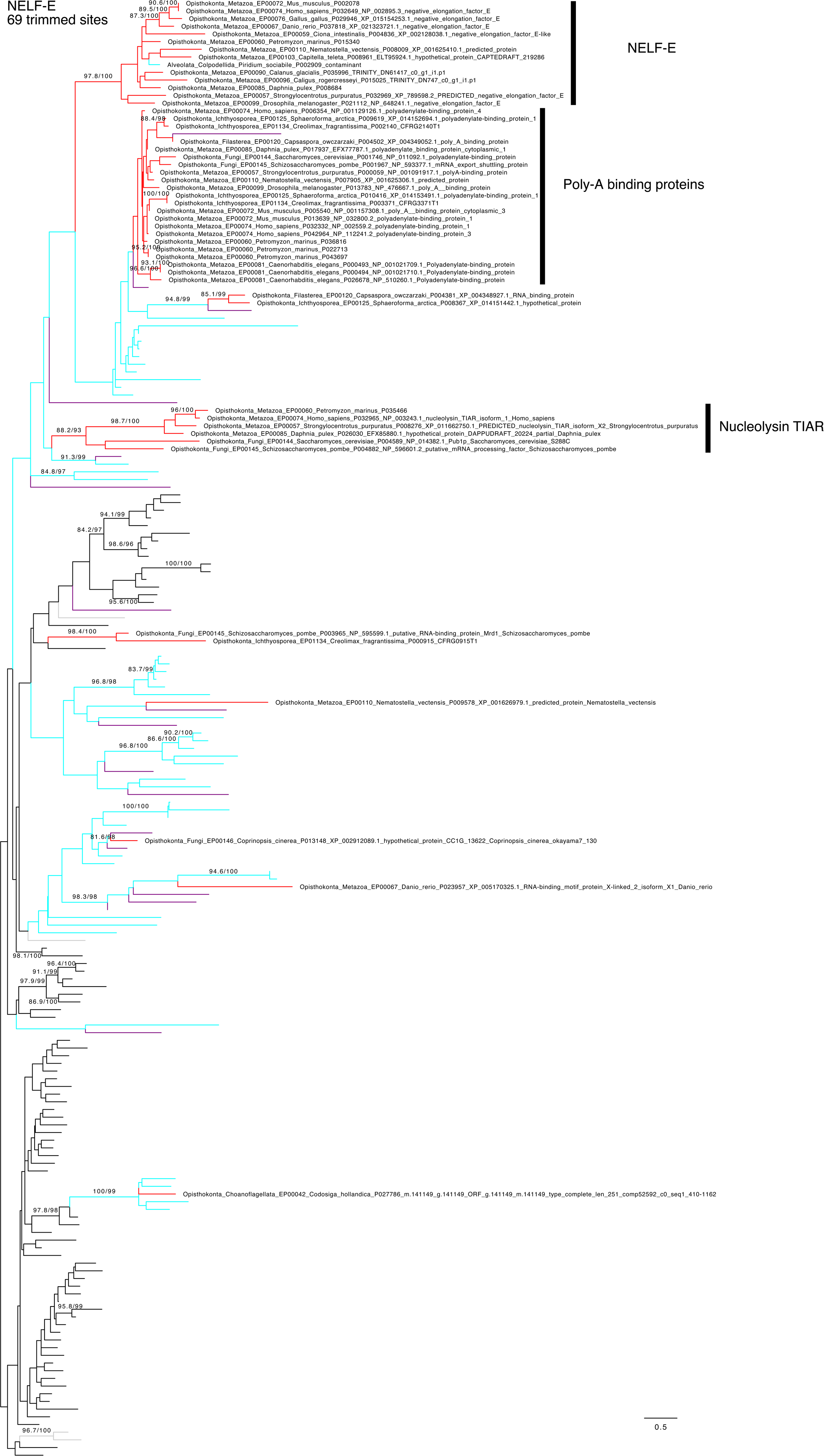
Selected positions of a multiple sequence alignment of NELF-E and paralogous sequences from selected opisthokonts. Paralogous sequences include polyadenylate-binding proteins and nucleolysin TIAR. Amino acids are shaded according to hydrophobicity value, where red is most hydrophobic and blue is the most hydrophilic, and numbered according to the Homo sapiens NELF-E (black numbers) and polyadenylate-binding protein 1 (red) sequences. The RNA Recognition Motif shared by these and other RNA-binding proteins is shown in black above the sequence. Alignment constructed with MAFFT-L-INS-I, visualized with Geneious® 2023.2.1, and edited with Affinity Designer 2.

**Figure S8:**
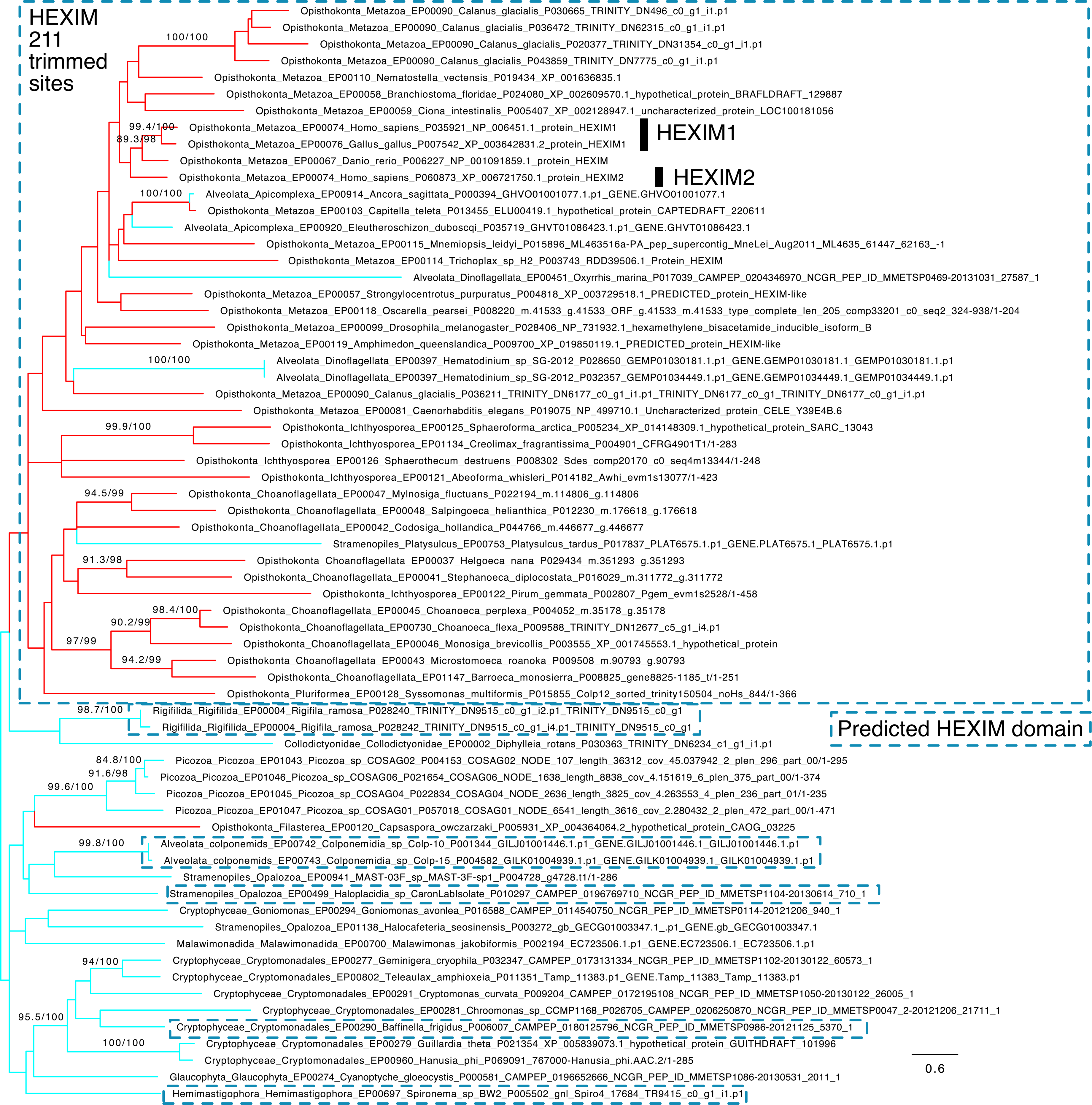
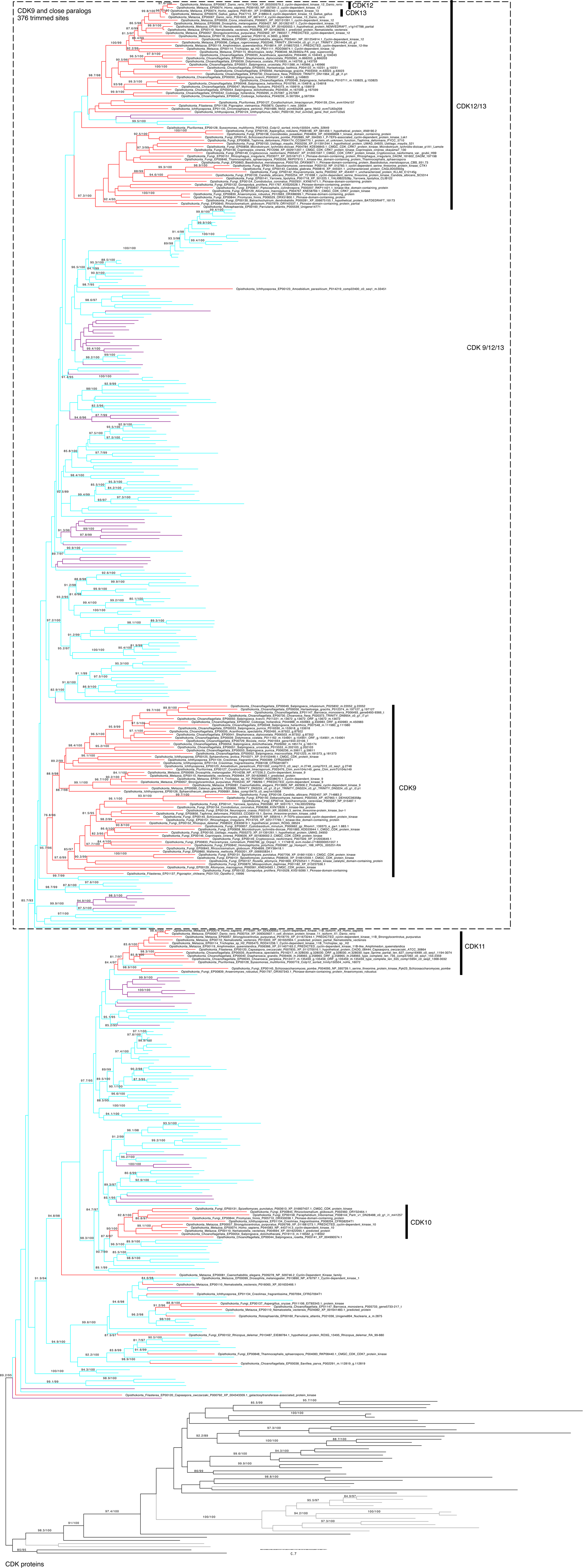
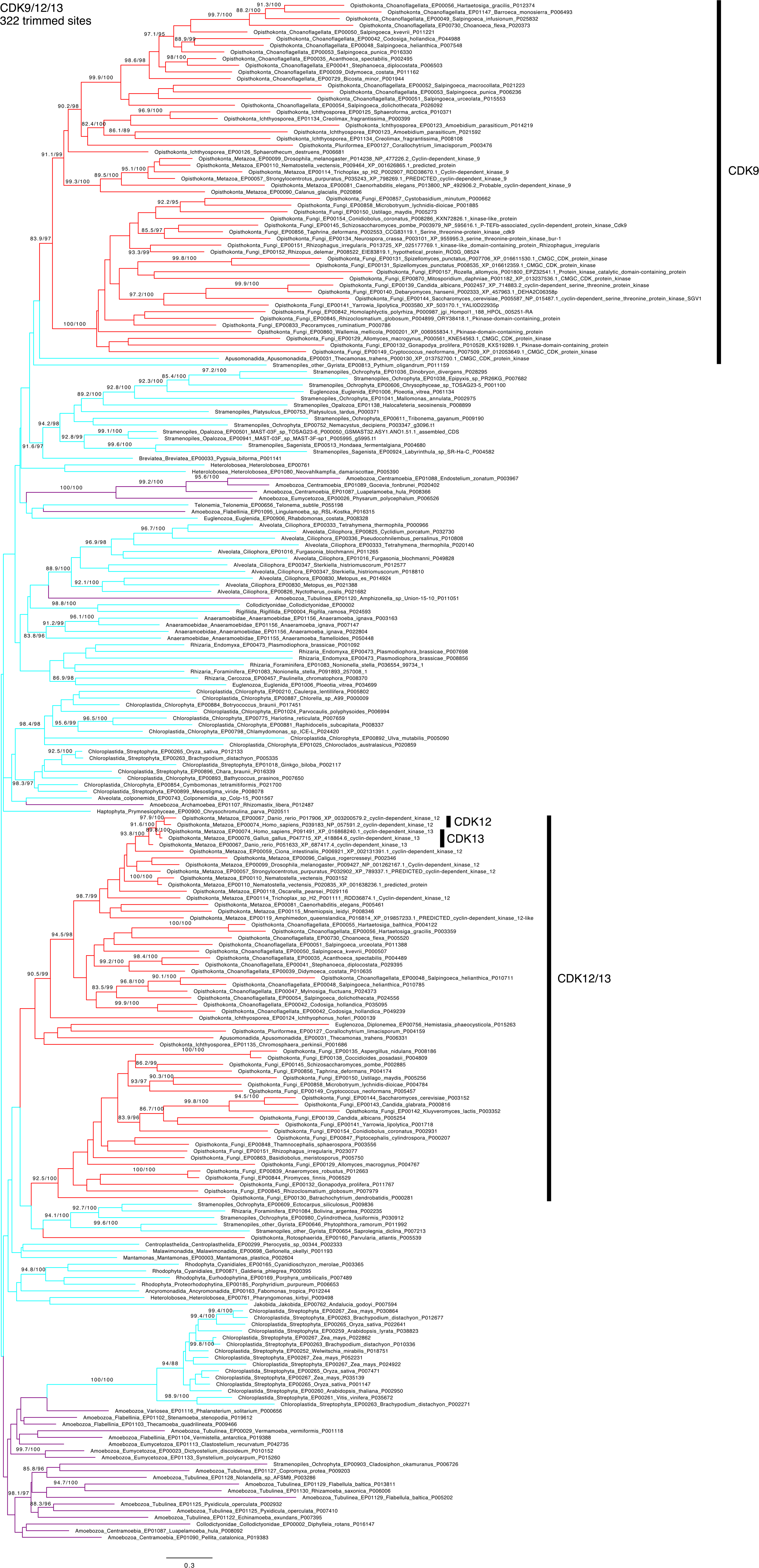
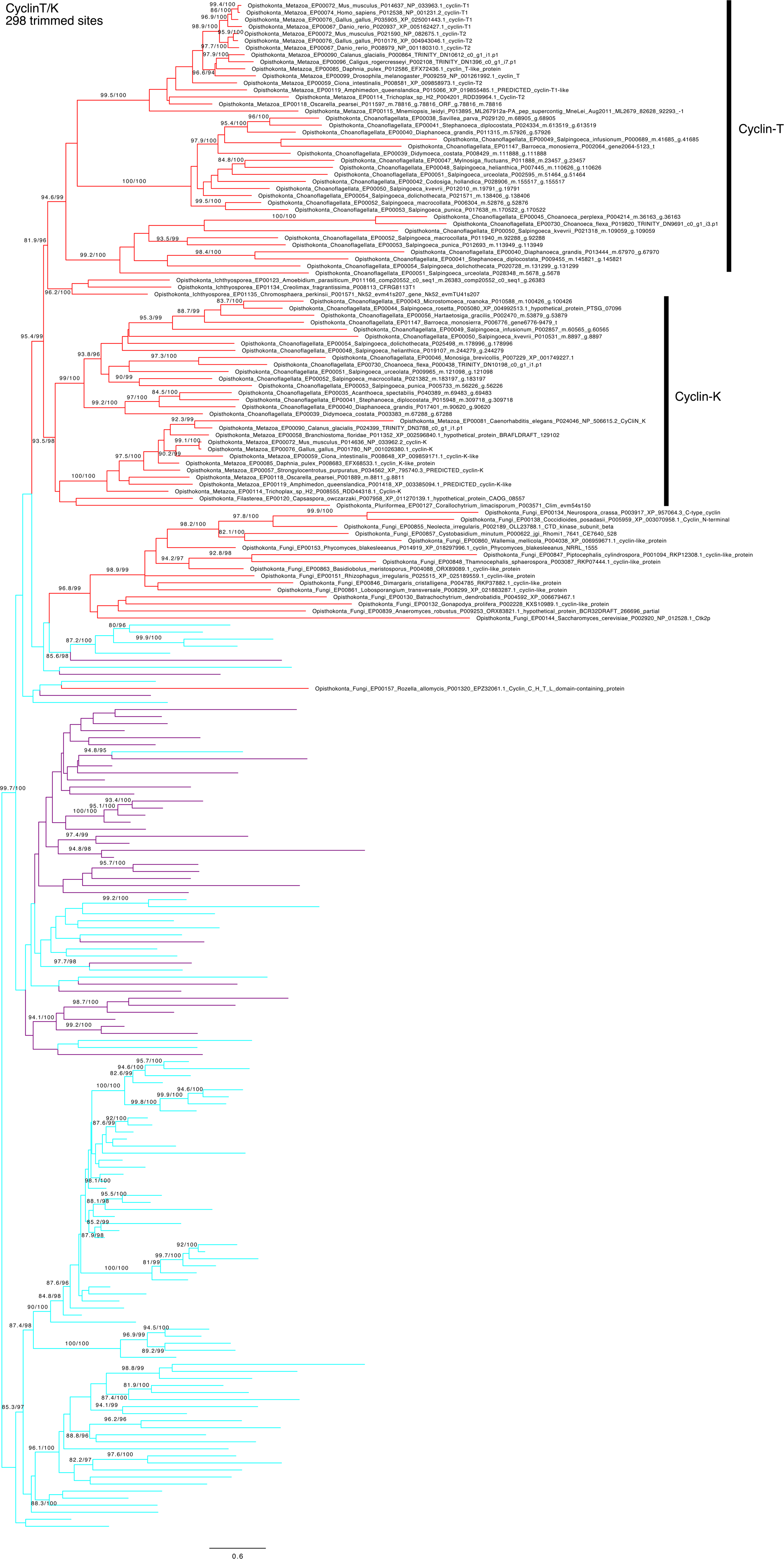
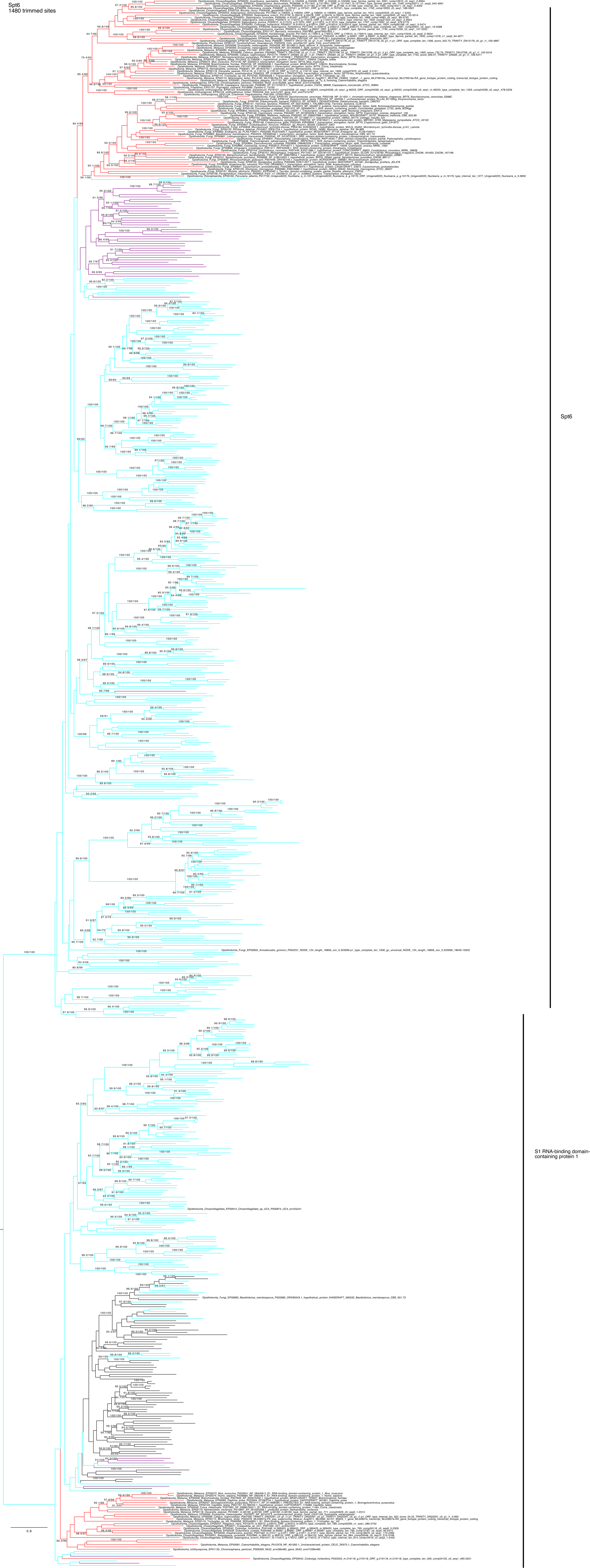
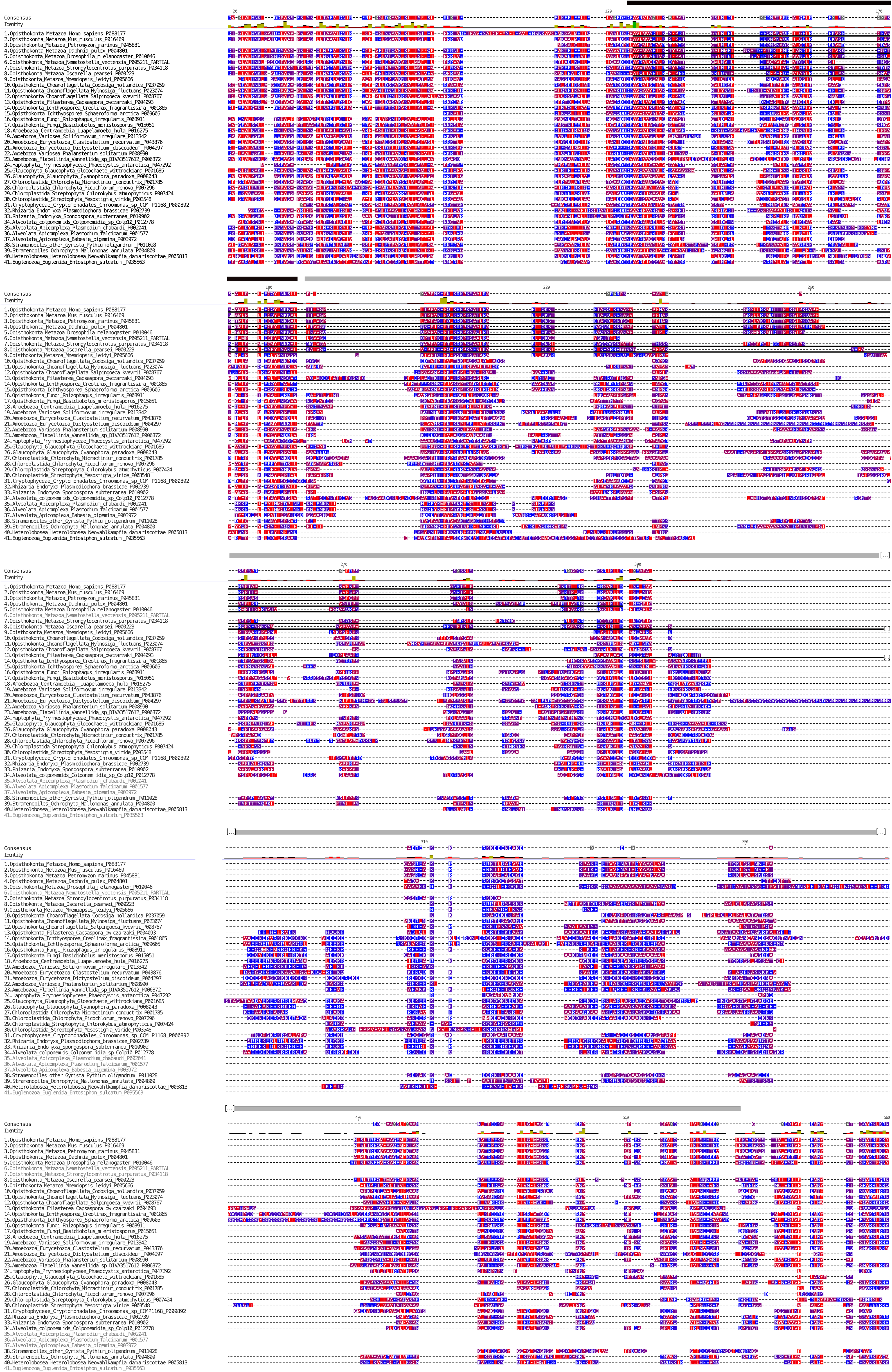
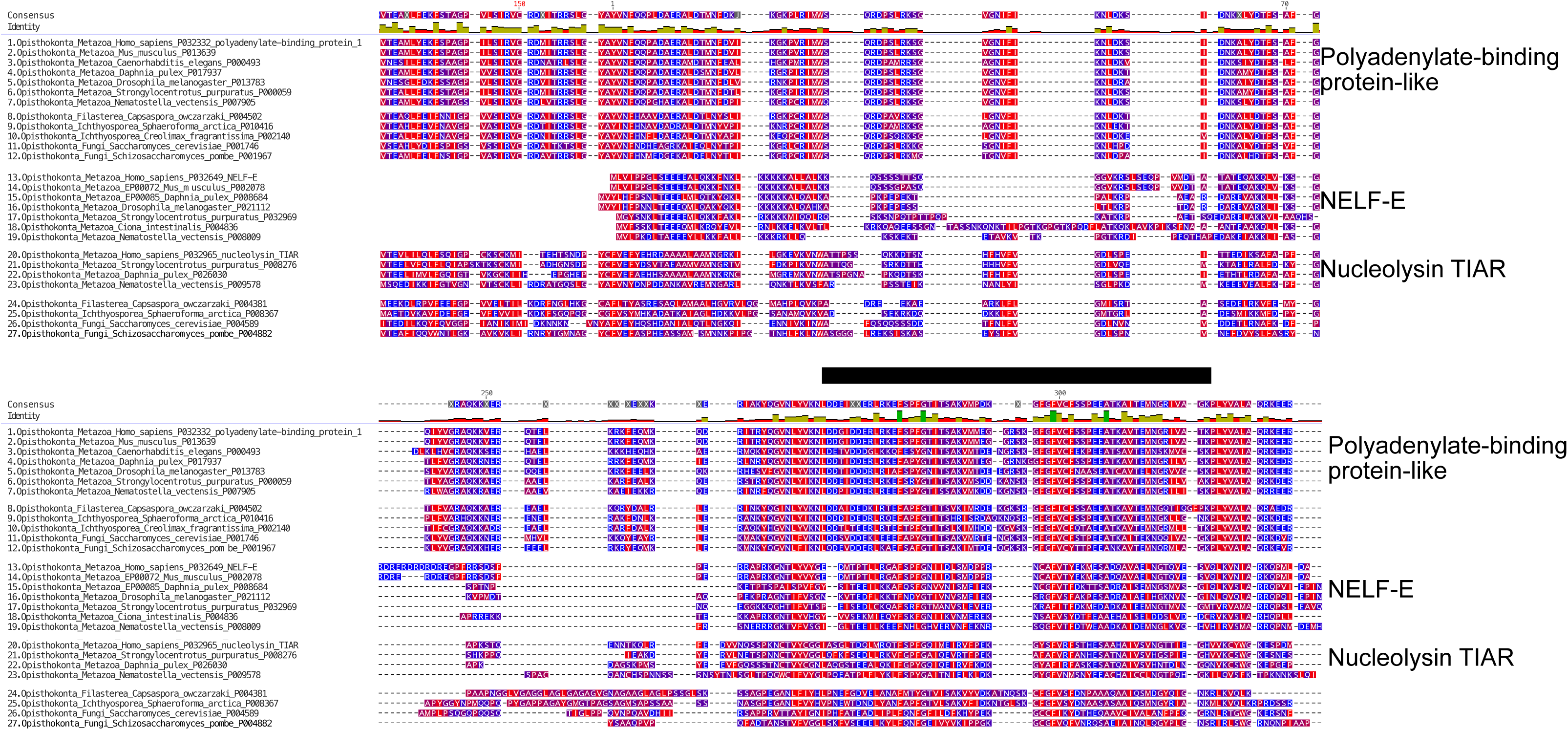
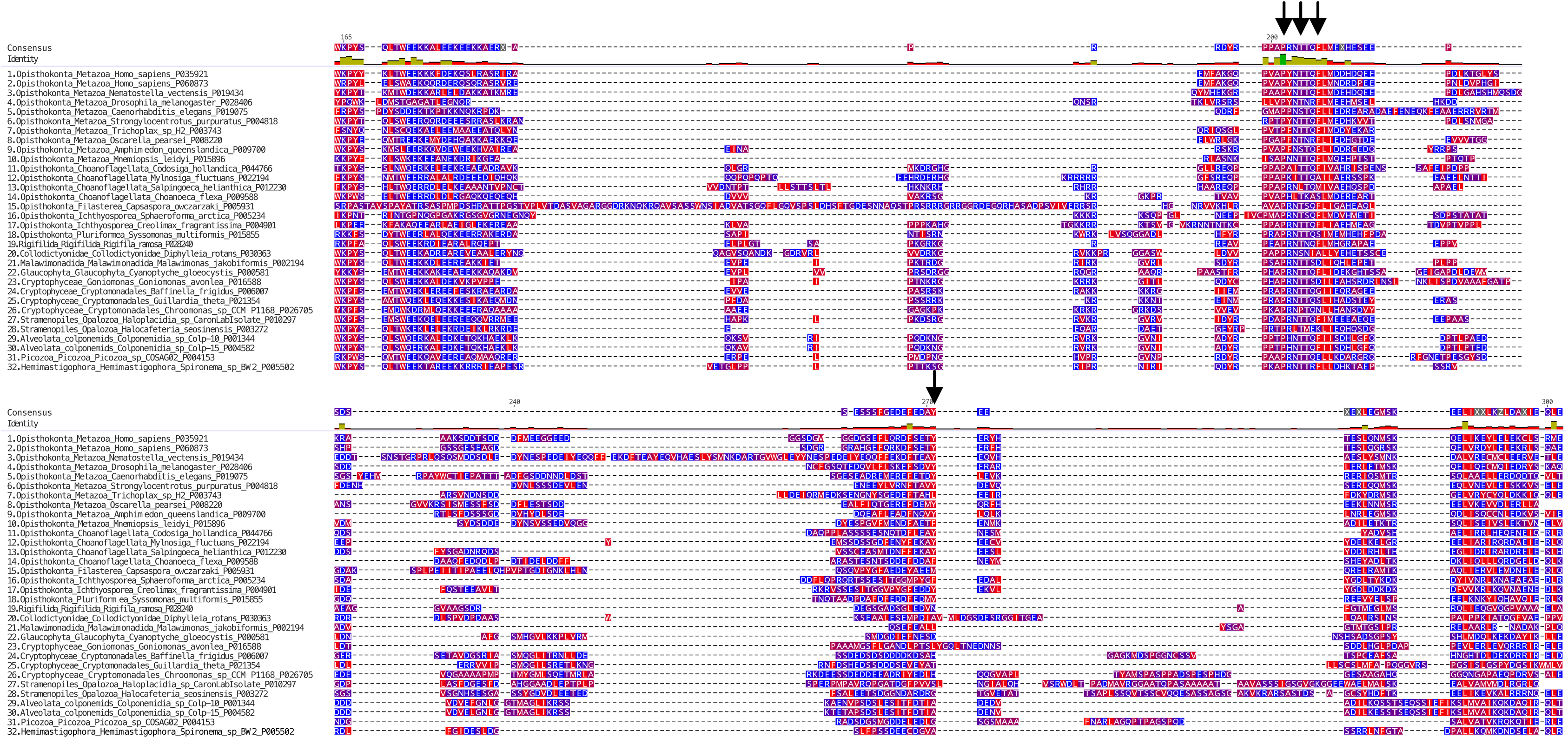
Selected positions of a multiple sequence alignment of HEXIM sequences from selected eukaryotes. Amino acids are shaded according to hydrophobicity value, where red is most hydrophobic and blue is the most hydrophilic, and numbered according to the Homo sapiens HEXIM1 sequence, and trimmed to the Homo sapiens HEXIM1 HEXIM Pfam domain. Arrows show Homo sapiens HEXIM1 residues Pro202, Thr205, Phe207 and Tyr271, crucial for the recruitment and inhibition of P-TEFb in human cells. Alignment constructed with MAFFT-L-INS-I, visualized with Geneious® 2023.2.1, and edited with Affinity Designer 2.

**Figure S9:**
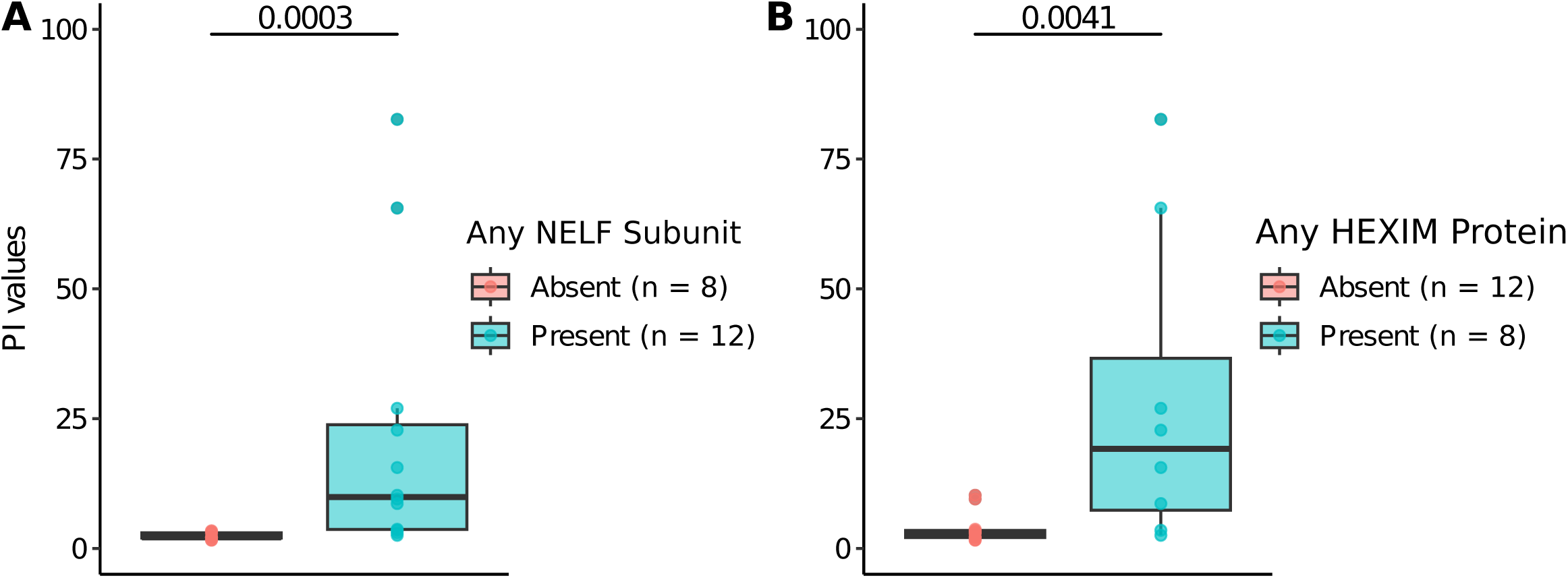
Association of NELF and HEXIM subunits with pausing index. Box and whiskers plots show pausing index values in each species. Samples are clustered by the presence or absence of any NELF subunits (A), and both HEXIM subunits (B). A two-sided Mann-Whitney test was used to compute p-values between the PI values.

**Figure S10:**
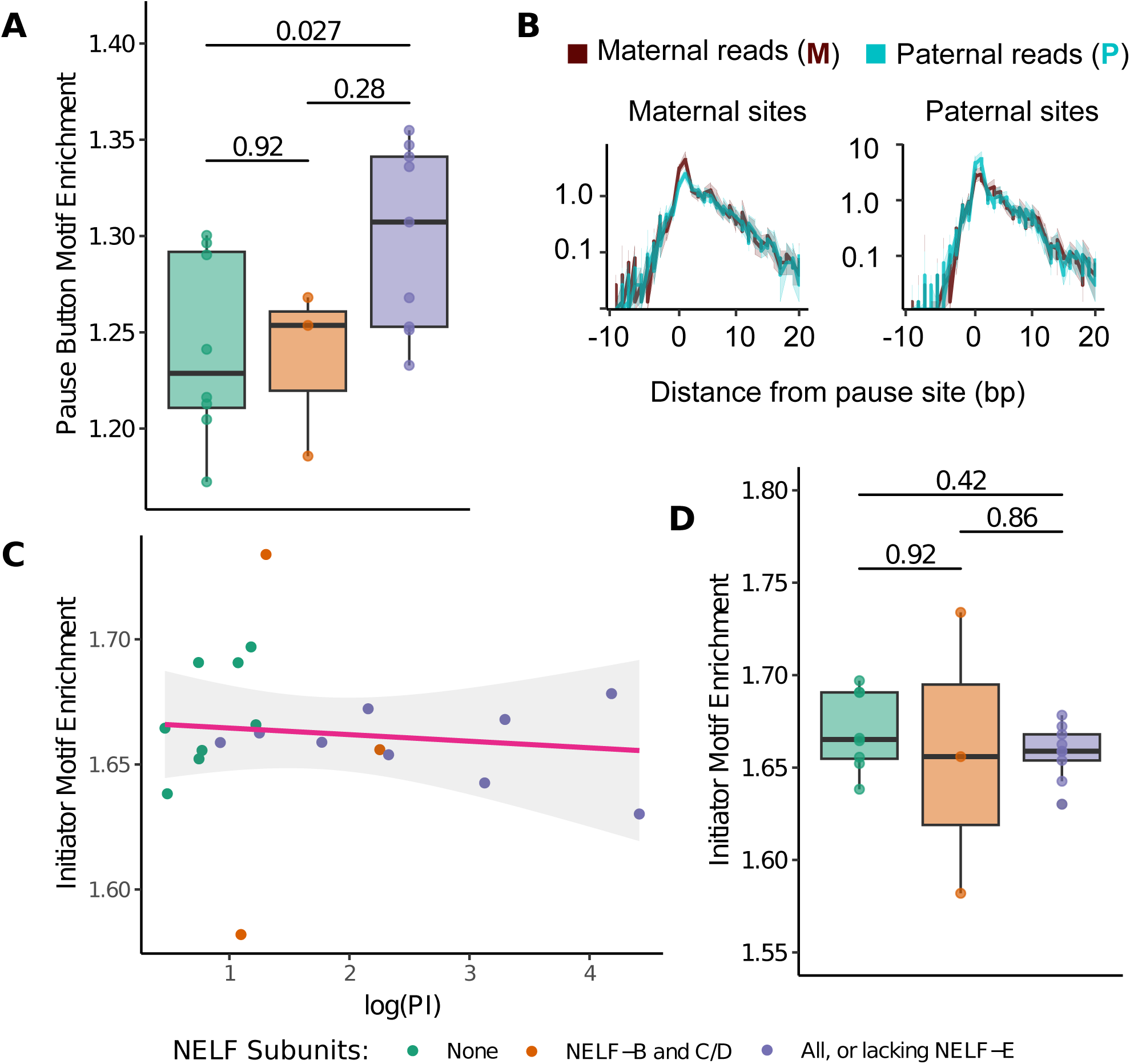
Pause motif search. (A) Box and whiskers plots depict enrichment of motif scores for the pause button in each species. Samples are clustered by the presence or absence of NELF-B, -C/D, and -A/E. A two-sided Mann-Whitney test was used to compute p-values. (B) Metaprofile plot of reads mapping to maternal and paternal alleles for genes with stronger pause motif on the maternal (left) or paternal (right) alleles. (C) Scatter plot of enrichment motif score of Initiator sequence plotted against the mean pausing index per species (R = 0.105, p = 0.661). Each dot is colored by the number of NELF subunits found in each sample. (D) Box and whiskers plots depict enrichment of motif scores for the initiator motif in each species. Samples are clustered by the presence or absence of NELF-B, -C/D, and -A/E. A two-sided Mann-Whitney test was used to compute p-values.

**Figure S11:**
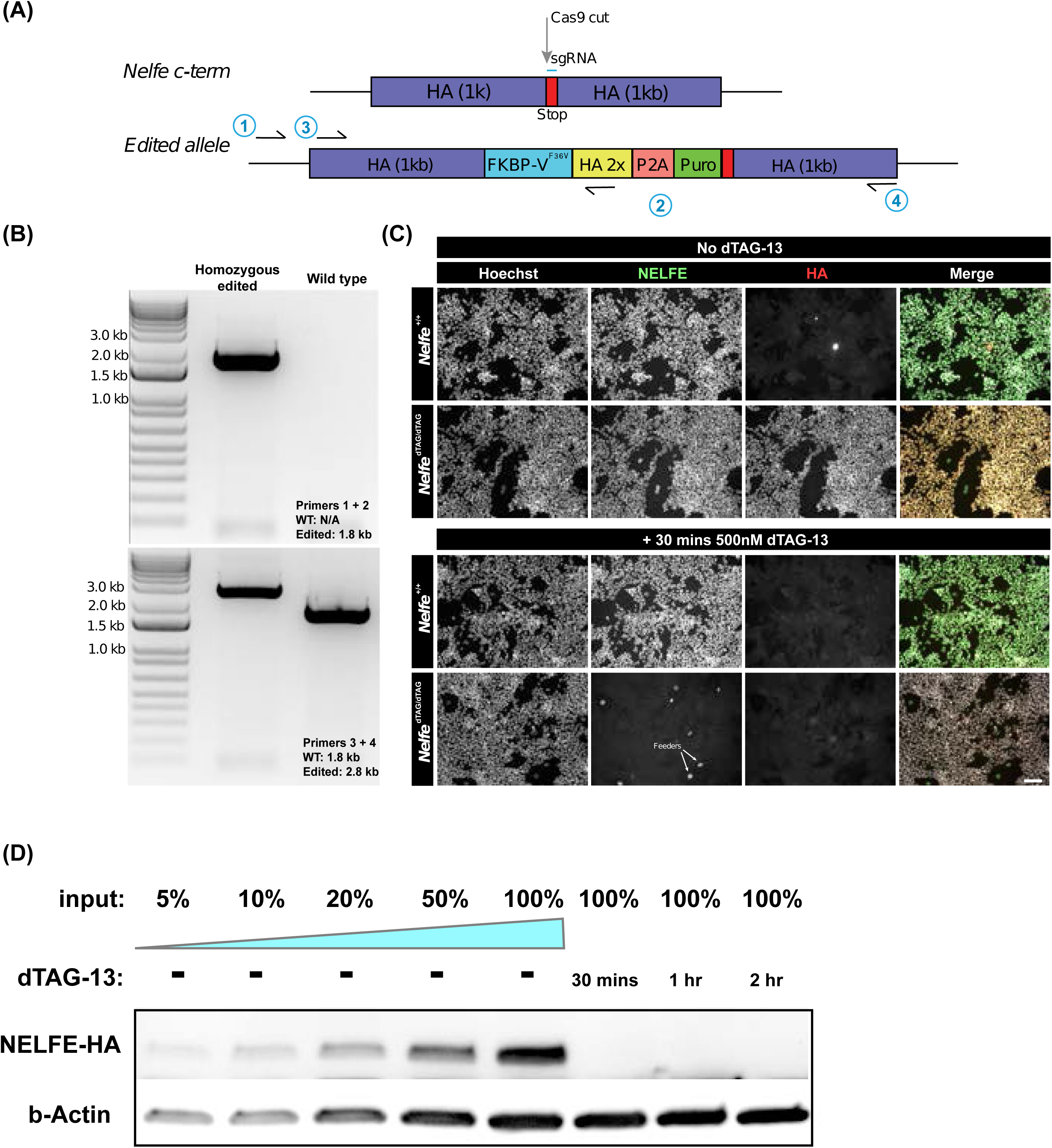
*nelfe-FKBP12* homozygous cell line generation. (A) Schematic of CRISPR design to add the FBBP12 tag at the *nelfe* locus. (B) PCR validation of CRISPR insertion of the FKBP12 tag. (C) Microscopy images evaluating the degradation efficiency before (top) and after a 30min treatment with 500nM dTAG-13 (bottom) in the edited and unedited cell lines. Hoechst was used as a nuclear control, while anti-HA antibodies measure the added tag, and anti-NELFE measures the NELF-E protein level. Arrows point out the presence of Feeder cells. (D) Degradation efficiency of NELF-E as measured by western blotting. b-Actin was used as a loading control, while anti-HA measures the level of NELFE-HA protein. Input denotes the relative amount of total protein loaded.

**Figure S12:**
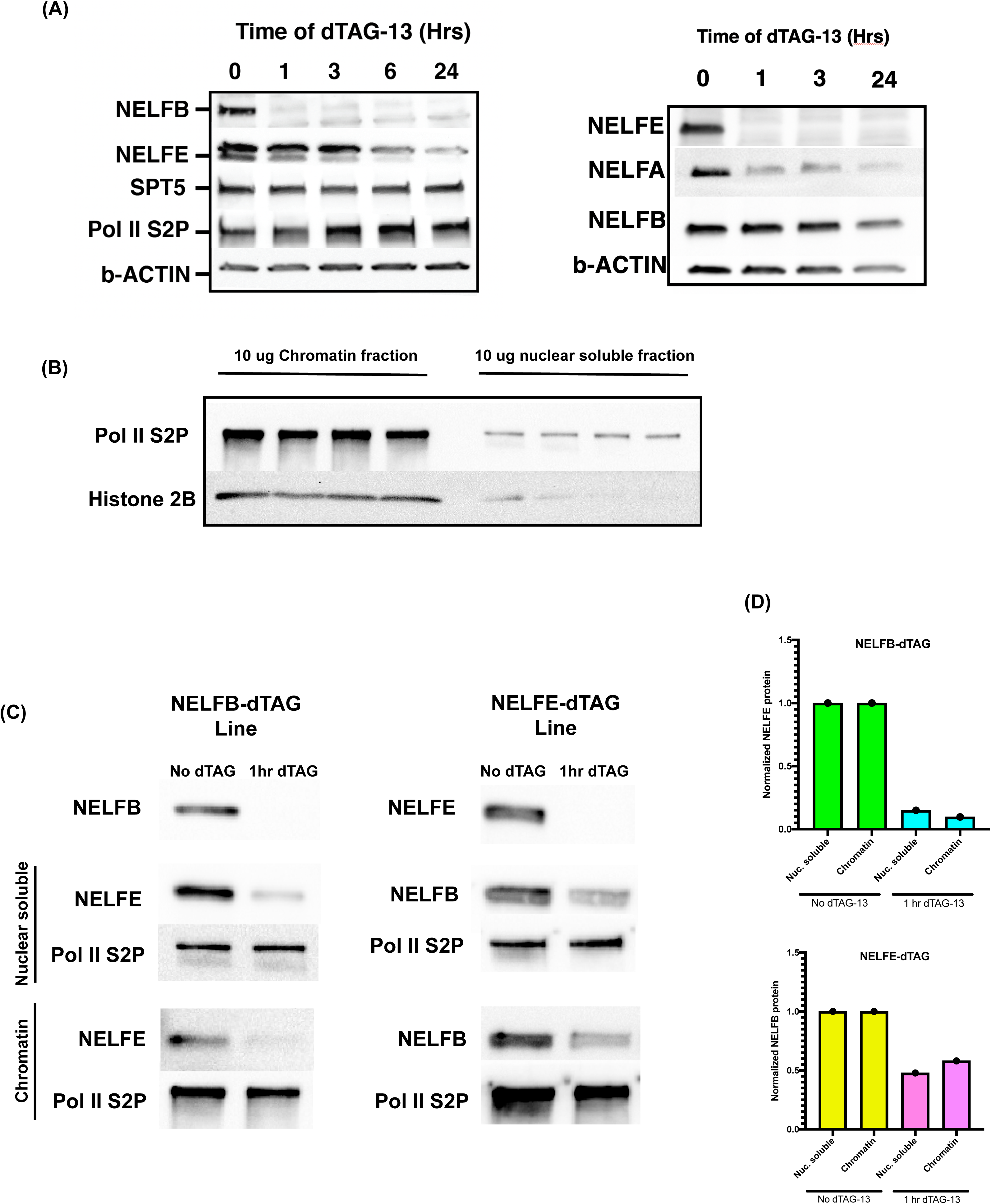
*nelfb and nelfe-FKBP12* homozygous cell line validation. (A) Western blot of whole cells following NELF-B (left) or NELF-E (right) degradation with 500nM dTAG-13 for 0 to 24h of treatment. (B) Western blot validation of chromatin fraction vs nuclear soluble fractionation. (C) Western blot of NELF-B and -E proteins after degradation of either protein for 1h. Both nuclear-soluble and chromatin-bound proteins were analyzed. (D) Quantification of western blot signal in (C) for NELF-E after degradation of NELF-B, and vice-versa.

**Figure S13:**
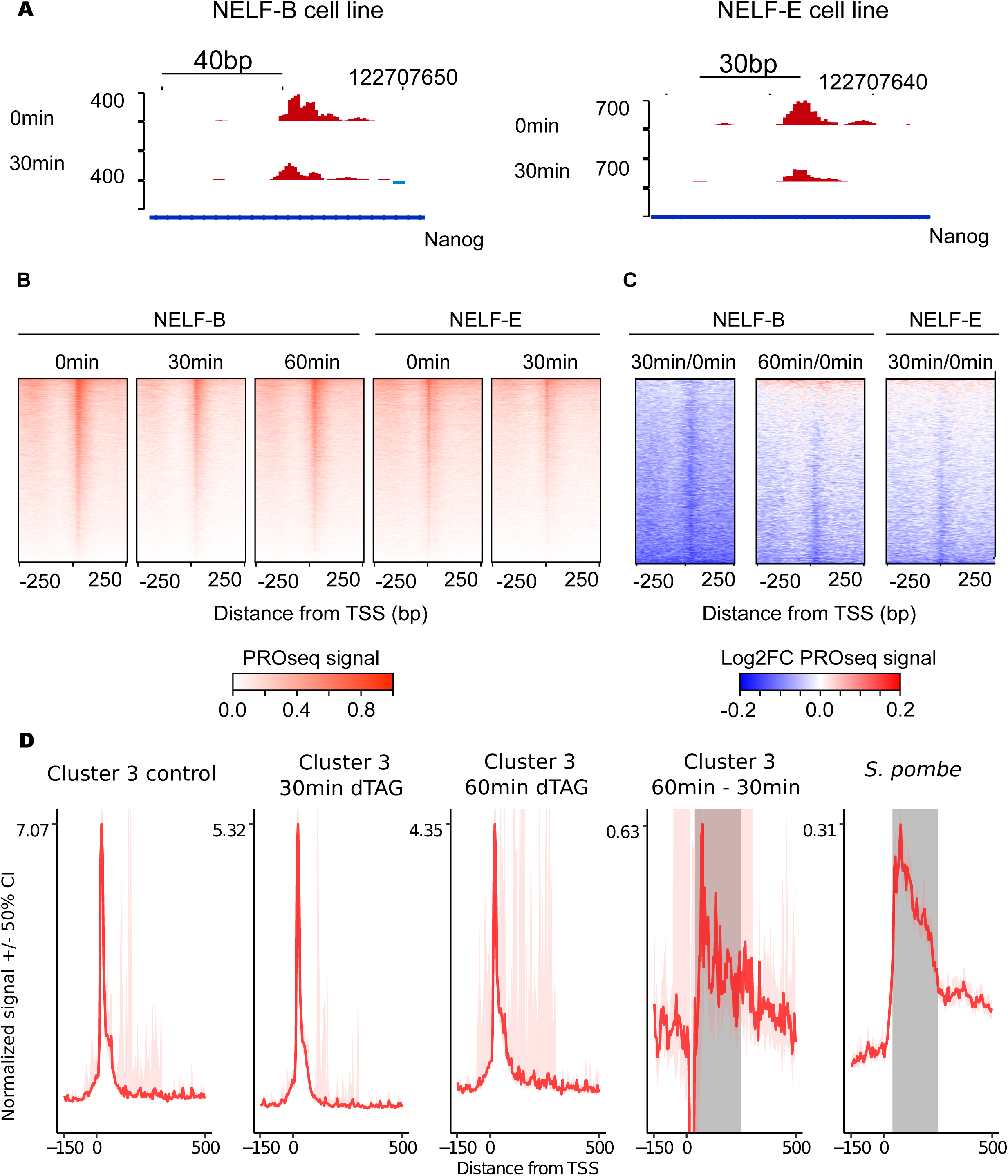
Effect of NELF-B and NELF-E degradation on Pol II distribution. (A) WashU browser shots at the Nanog gene locus before and after NELF-B and -E degradation. (B) Heat maps of spike-in normalized PRO-seq signal (C) Heatmaps of log2 fold changes of normalized PRO-seq signal relative to untreated controls. All heat maps are centered on active TSSs in mESCs. (D) Metaprofiles of cluster 3 genes at each dTAG time point, the recovery of pause-like behavior between 30 and 60 minutes, and the proto-pause observed in *S. pombe* for reference. The proto-pause region is shaded in the two rightmost panels.

**Figure S14:**
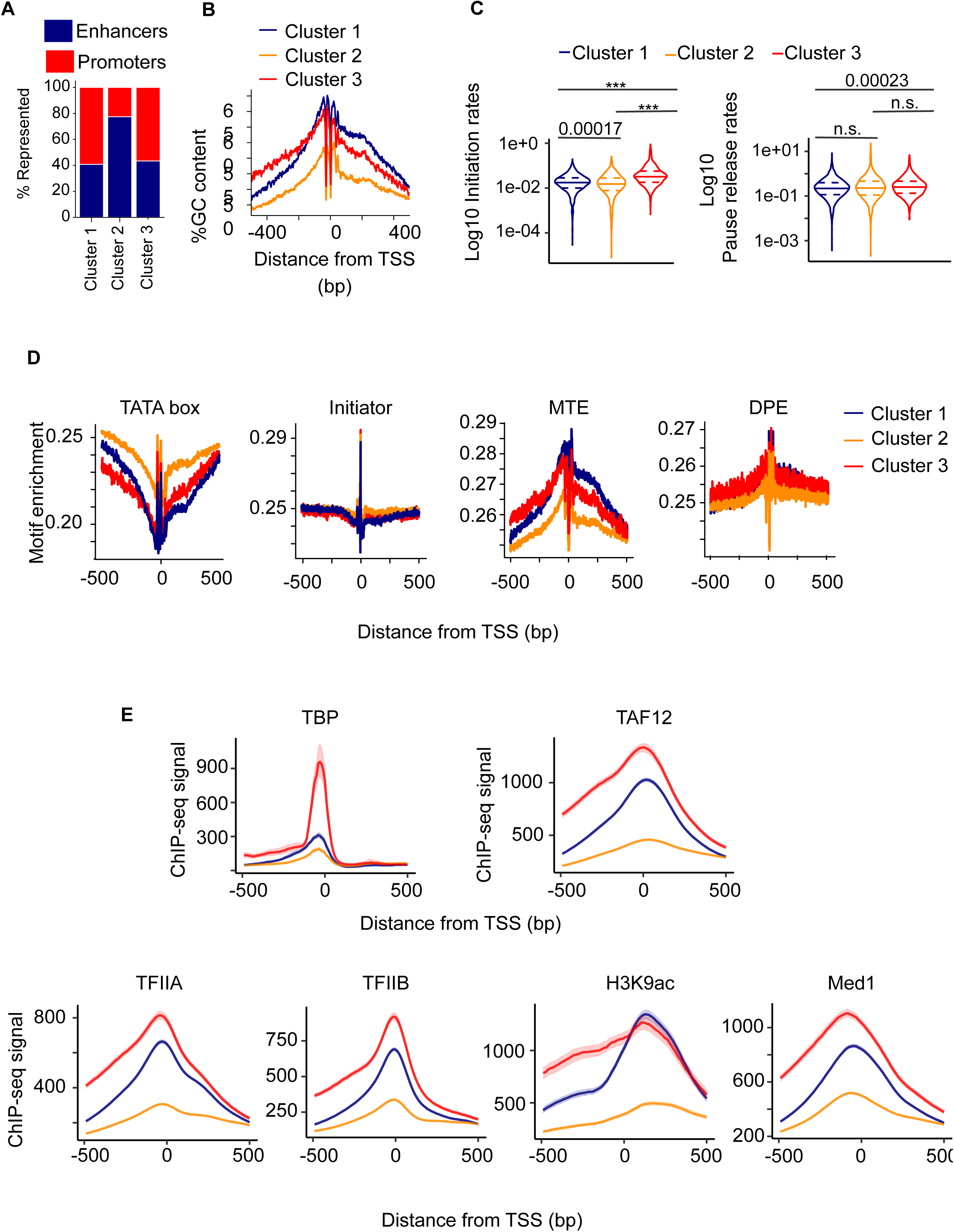
Characterization of transcription recovery clusters after NELF-B degradation. (A) Bar plots depict the percentage of transcribed enhancers and gene promoters in each cluster defined in Figure 3E. (B) Enrichment profiles of the pause motif published in Watts *et al*., Am J Hum Genet (2019) plotted in a 1kb window centered on TSSs found in each of the clusters defined in Figure 3E. (C) Violin plots depict log10 transformed initiation (right) or pause release (left) rates in each cluster. A two-sided Mann-Whitney test was used to compute p-values, where n.s. defines non-significant p-values, and (***) p-values < 2.2e-16. (D) Plots depict the enrichment of the TATA box, Initiator, MTE, and DPE sequence motifs in each cluster in Figure 3E. (E) Meta profiles depict the enrichment of TBP, TAF-12, TFIIA, TFIIB, H3K9ac, and Med1 per cluster.

**Figure S15:**
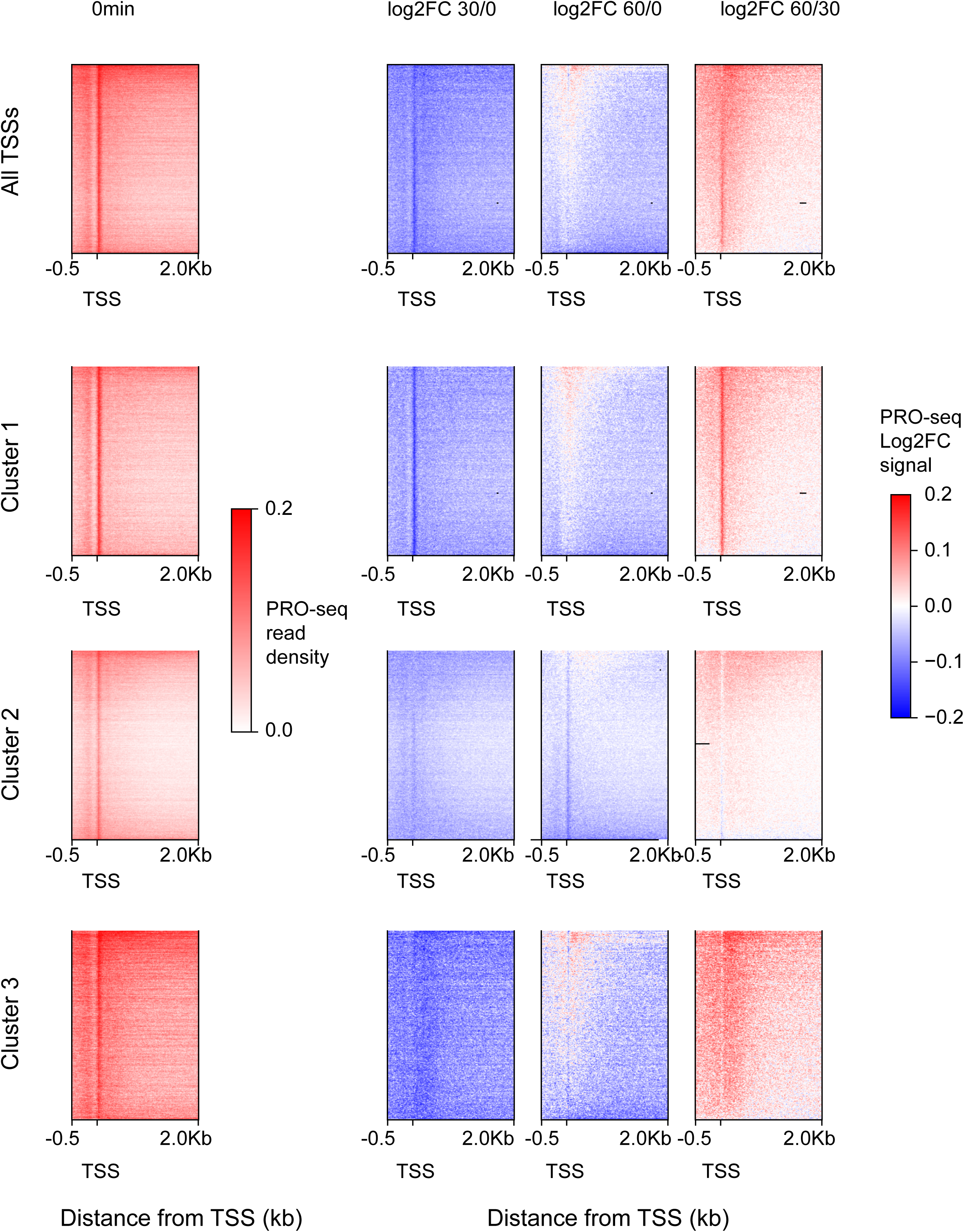
Pol II trickles into gene bodies effect after NELF-B degradation. Heatmaps of log2 fold changes in PRO-seq signal in NELF-B tagged cell lines at all TSSs, Clusters 1, 2, and 3 (in this order from top to bottom rows). The heatmaps depict log2 fold change relative for the following comparisons (from left to right columns): untreated PRO-seq signal, log2 fold change for 30min/0min, 60min/0min, and 60min/30min of dTAG-13 treatment.

**Figure S16:**
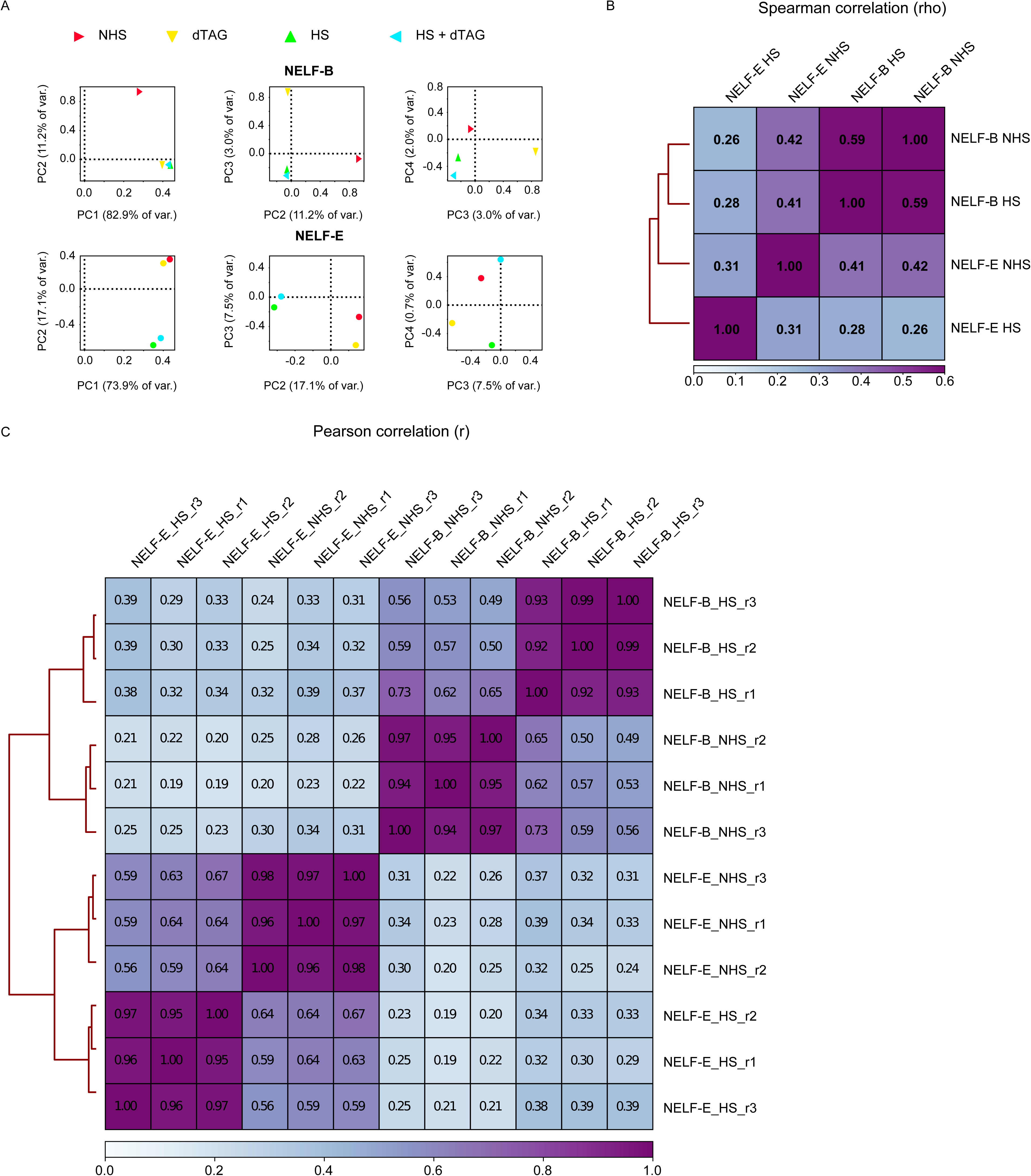
Correlations between heat shock PRO-seq data. (A) Principal component analysis (PCA) of non-heat shock (NHS), dTAG-13 treatment (dTAG), heat shock (HS), and a pre-treatment of dTAG-13 followed by heat shock (HS+dTAG). (B) Clustered dendrogram of spearman correlations (rho) between the NELF-B and NELF-E HS and NHS conditions before any protein degradation. (C) Clustered dendrogram of Pearson correlations (r) between all three independent replicates of HS and NHS in both NELF-B and NELF-E cell lines before degradation. R1, r2, and r3 represent replicate numbers.

**Figure S17:**
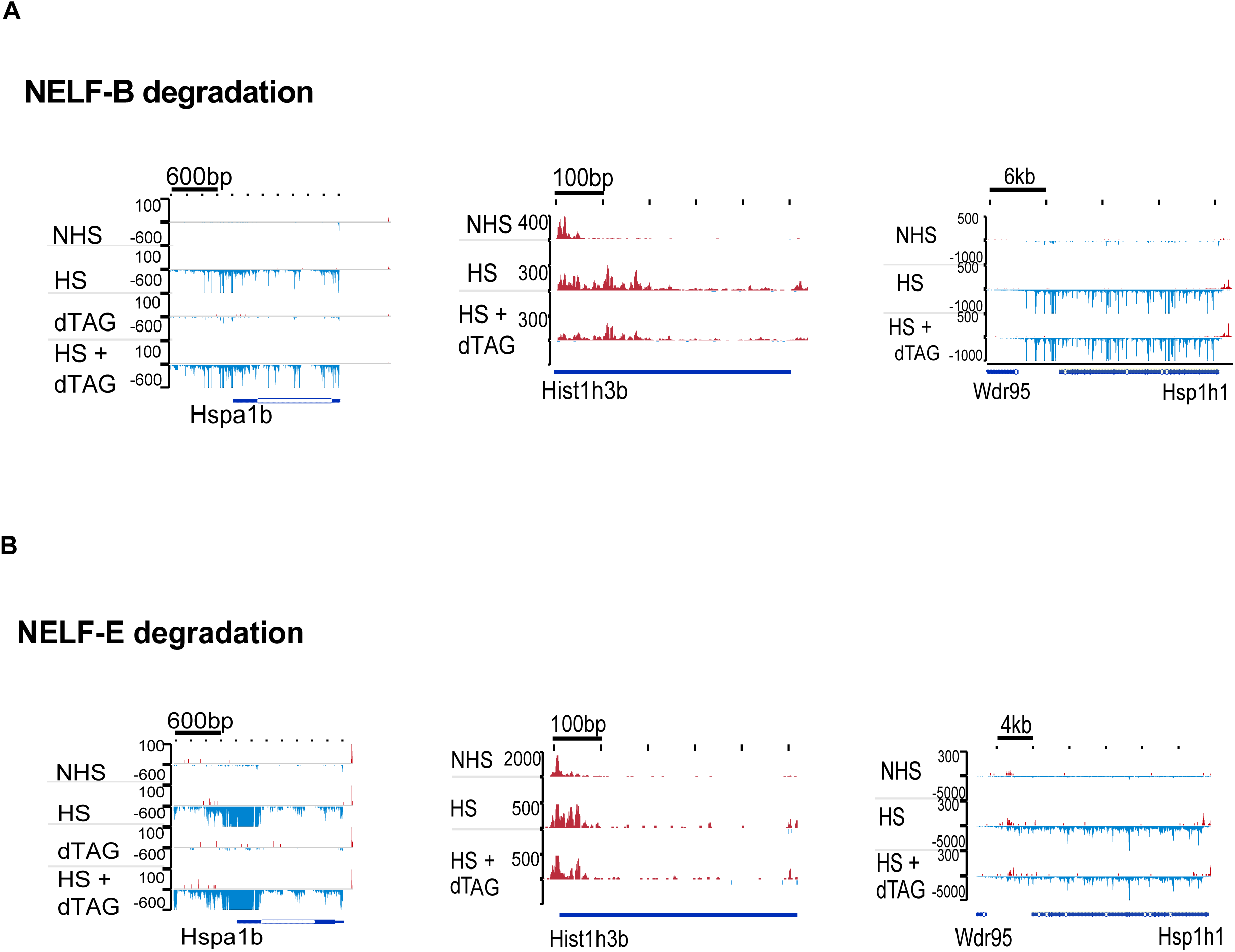
Studying the heat shock response after NELF-B or NELF-E degradation. WashU browser shots at the heat-triggered genes (Hist1h3b, Hsp1h1, Hist1h3b) in the NELF-B (A) and NELF-E (B) edited cell lines.

**Figure S18:**
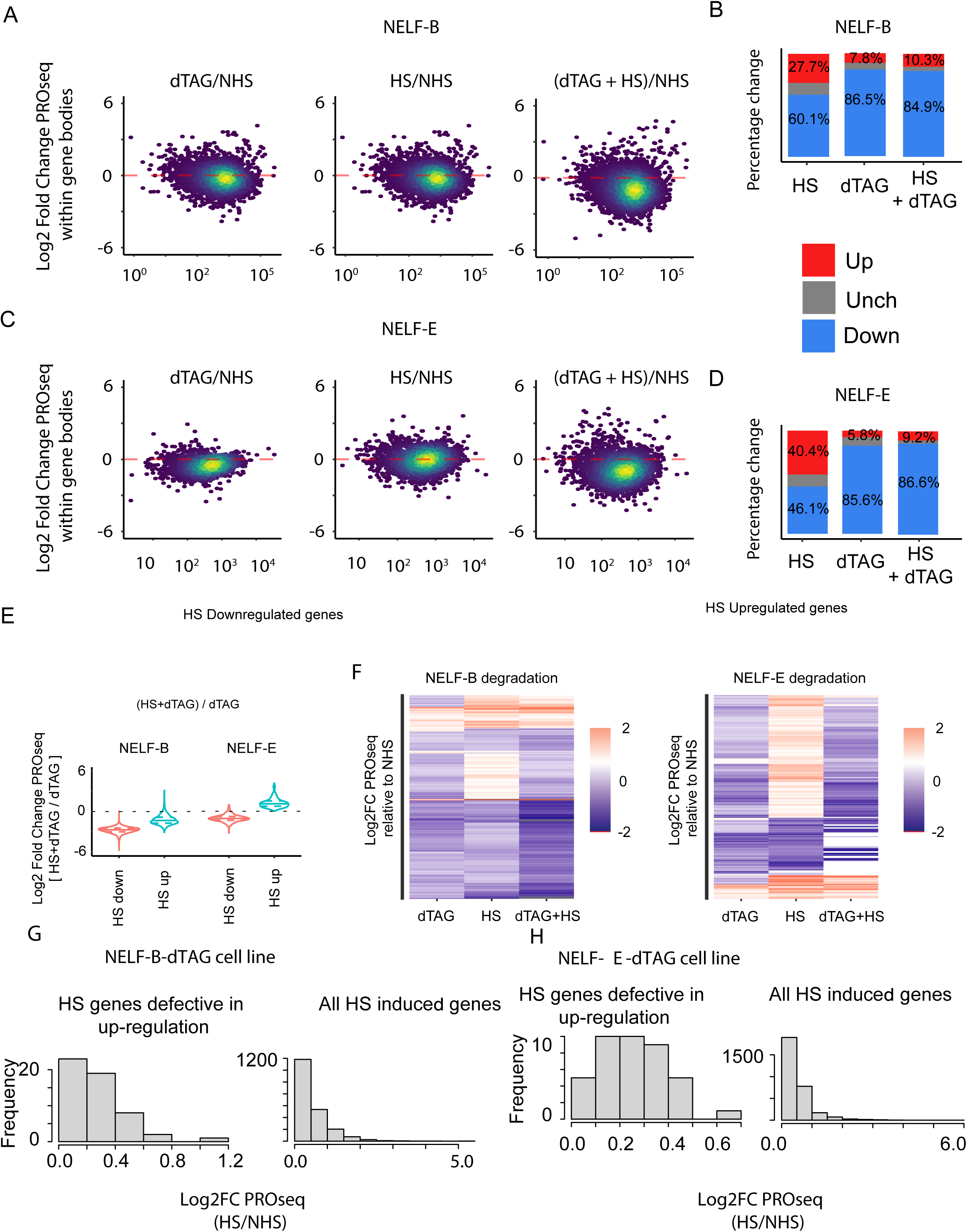
Assessing the heat shock response in *nelfb-fkbp12* and *nelfe-fkbp12* homozygous cell lines. (A) & (C) MA plots show the log2 fold change in gene body PRO-seq signal when comparing dTAG-13 treatment (dTAG) with non-heat shock (NHS), heat shock (HS) with NHS, and the dTAG-13 pre-treatment followed by HS with NHS in the *nelfb-fkbp12* (A) and the *nelfb-fkbp12* (C) cell lines. (B) & (D) Bar plots depict the percentage of upregulated (red), downregulated (blue), and unchanged (gray) genes in the *nelfb-fkbp12* (B) and the *nelfb-fkbp12* (D) cell lines. (E) Violin plots show the log2 fold change in gene body PRO-seq signal when comparing the pre-treatment degradation of either NELF-B (left) or NELF-E (right) at genes known to be upregulated (blue) or downregulated (red) after regular heat stress. (F) Heatmaps of log2 fold changes in PRO-seq signal in NELF-B (left) and NELF-E tagged (right) cell lines. The heatmaps rows depict fold changes relative to NHS for the following treatments: dTAG-13 treatment alone, HS alone, and dual treatment of dTAG-13 and HS. (G-H) Bar graphs at the heat-shock dependent genes that show a defect in up-regulation after NELF depletion. The graphs show the frequency of log2 fold changes in PRO-seq data when comparing the dual treatment of dTAG-13 followed by heat shock with a non-heat shock control. Data is presented for both the *nelfb-FKBP12* (left) and the *nelfe-FKBP12* (right) cell lines.

## References

1. Scholes, C., DePace, A.H., and Sánchez, Á. (2017). Combinatorial Gene Regulation through Kinetic Control of the Transcription Cycle. Cell Syst 4, 97–108.e9.

2. Mikhailov, K.V., Konstantinova, A.V., Nikitin, M.A., Troshin, P.V., Rusin, L.Y., Lyubetsky, V.A., Panchin, Y.V., Mylnikov, A.P., Moroz, L.L., Kumar, S., et al. (2009). The origin of Metazoa: a transition from temporal to spatial cell differentiation. Bioessays 31, 758–768.

3. Arenas-Mena, C. (2010). Indirect development, transdifferentiation and the macroregulatory evolution of metazoans. Philos. Trans. R. Soc. Lond. B Biol. Sci. 10.1098/rstb.2009.0253.

4. Arenas-Mena, C. (2017). The origins of developmental gene regulation. Evol. Dev. 19, 96–107.

5. Core, L., and Adelman, K. (2019). Promoter-proximal pausing of RNA polymerase II: a nexus of gene regulation. Genes Dev. 33, 960–982.

6. Adelman, K., and Lis, J.T. (2012). Promoter-proximal pausing of RNA polymerase II: emerging roles in metazoans. Nat. Rev. Genet. 13, 720–731.

7. Tempo and Mode of Genome Evolution in the Budding Yeast Subphylum (2018). Cell 175, 1533– 1545.e20.

8. Nagy, L.G., Ohm, R.A., Kovács, G.M., Floudas, D., Riley, R., Gácser, A., Sipiczki, M., Davis, J.M., Doty, S.L., de Hoog, G.S., et al. (2014). Latent homology and convergent regulatory evolution underlies the repeated emergence of yeasts. Nat. Commun. 5, 1–8.

9. Vill, A.C., Rice, E.J., De Vlaminck, I., Danko, C.G., and Brito, I.L. (2022). Run-on sequencing reveals nascent transcriptomics of the human microbiome. bioRxiv, 2022.04.22.489220. 10.1101/2022.04.22.489220.

10. Booth, G.T., Wang, I.X., Cheung, V.G., and Lis, J.T. (2016). Corrigendum: Divergence of a conserved elongation factor and transcription regulation in budding and fission yeast. Genome Res. 26, 1010–1011.

11. Rougvie, A.E., and Lis, J.T. (1988). The RNA polymerase II molecule at the 5’ end of the uninduced hsp70 gene of D. melanogaster is transcriptionally engaged. Cell 54, 795–804.

12. Muse, G.W., Gilchrist, D.A., Nechaev, S., Shah, R., Parker, J.S., Grissom, S.F., Zeitlinger, J., and Adelman, K. (2007). RNA polymerase is poised for activation across the genome. Nat. Genet. 39, 1507– 1511.

13. Core, L.J., Waterfall, J.J., and Lis, J.T. (2008). Nascent RNA sequencing reveals widespread pausing and divergent initiation at human promoters. Science 322, 1845–1848.

14. Jonkers, I., Kwak, H., and Lis, J.T. (2014). Genome-wide dynamics of Pol II elongation and its interplay with promoter proximal pausing, chromatin, and exons. Elife 3, e02407.

15. Danko, C.G., Hah, N., Luo, X., Martins, A.L., Core, L., Lis, J.T., Siepel, A., and Kraus, W.L. (2013). Signaling pathways differentially affect RNA polymerase II initiation, pausing, and elongation rate in cells. Mol. Cell 50, 212–222.

16. Zeitlinger, J., Stark, A., Kellis, M., Hong, J.-W., Nechaev, S., Adelman, K., Levine, M., and Young, R.A. (2007). RNA polymerase stalling at developmental control genes in the Drosophila melanogaster embryo. Nat. Genet. 39, 1512–1516.

17. Abuhashem, A., Chivu, A.G., Zhao, Y., Rice, E.J., Siepel, A., Danko, C.G., and Hadjantonakis, A.-K. (2022). RNA Pol II pausing facilitates phased pluripotency transitions by buffering transcription. Genes Dev. 36, 770–789.

18. Williams, L.H., Fromm, G., Gokey, N.G., Henriques, T., Muse, G.W., Burkholder, A., Fargo, D.C., Hu, G., and Adelman, K. (2015). Pausing of RNA polymerase II regulates mammalian developmental potential through control of signaling networks. Mol. Cell 58, 311–322.

19. Buckley, M.S., Kwak, H., Zipfel, W.R., and Lis, J.T. (2014). Kinetics of promoter Pol II on Hsp70 reveal stable pausing and key insights into its regulation. Genes Dev. 28, 14–19.

20. Aoi, Y., Shah, A.P., Ganesan, S., Soliman, S.H.A., Cho, B.-K., Goo, Y.A., Kelleher, N.L., and Shilatifard, A. (2022). SPT6 functions in transcriptional pause/release via PAF1C recruitment. Mol. Cell 82, 3412– 3423.e5.

21. Marshall, N.F., and Price, D.H. (1995). Purification of P-TEFb, a transcription factor required for the transition into productive elongation. J. Biol. Chem. 270, 12335–12338.

22. Li, Q., Price, J.P., Byers, S.A., Cheng, D., Peng, J., and Price, D.H. (2005). Analysis of the Large Inactive P-TEFb Complex Indicates That It Contains One 7SK Molecule, a Dimer of HEXIM1 or HEXIM2, and Two P-TEFb Molecules Containing Cdk9 Phosphorylated at Threonine 186*. J. Biol. Chem. 280, 28819–28826.

23. Booth, G.T., Wang, I.X., Cheung, V.G., and Lis, J.T. (2016). Divergence of a conserved elongation factor and transcription regulation in budding and fission yeast. Genome Res. 26, 799–811.

24. Booth, G.T., Parua, P.K., Sansó, M., Fisher, R.P., and Lis, J.T. (2018). Cdk9 regulates a promoter-proximal checkpoint to modulate RNA polymerase II elongation rate in fission yeast. Nat. Commun. 9, 543.

25. Kwak, H., Fuda, N.J., Core, L.J., and Lis, J.T. (2013). Precise maps of RNA polymerase reveal how promoters direct initiation and pausing. Science 339, 950–953.

26. Kruesi, W.S., Core, L.J., Waters, C.T., Lis, J.T., and Meyer, B.J. (2013). Condensin controls recruitment of RNA polymerase II to achieve nematode X-chromosome dosage compensation. Elife 2, e00808.

27. Hetzel, J., Duttke, S.H., Benner, C., and Chory, J. (2016). Nascent RNA sequencing reveals distinct features in plant transcription. Proc. Natl. Acad. Sci. U. S. A. 113, 12316–12321.

28. Lozano, R., Booth, G.T., Omar, B.Y., Li, B., Buckler, E.S., Lis, J.T., Del Carpio, D.P., and Jannink, J.-L. (2021). RNA polymerase mapping in plants identifies intergenic regulatory elements enriched in causal variants. G3 11. 10.1093/g3journal/jkab273.

29. Choi, J.Y., Platts, A.E., Johary, A., Purugganan, M.D., and Joly-Lopez, Z. (2022). Nascent transcription and the associated cis-regulatory landscape in rice. bioRxiv, 2022.07.06.498888. 10.1101/2022.07.06.498888.

30. Wang, Z., Chivu, A.G., Choate, L.A., Rice, E.J., Miller, D.C., Chu, T., Chou, S.-P., Kingsley, N.B., Petersen, J.L., Finno, C.J., et al. (2022). Prediction of histone post-translational modification patterns based on nascent transcription data. Nat. Genet. 54, 295–305.

31. Zhao, Y., Liu, L., and Siepel, A. Model-based characterization of the equilibrium dynamics of transcription initiation and promoter-proximal pausing in human cells. Preprint, 10.1101/2022.10.19.512929 10.1101/2022.10.19.512929.

32. Lee, C., Li, X., Hechmer, A., Eisen, M., Biggin, M.D., Venters, B.J., Jiang, C., Li, J., Pugh, B.F., and Gilmour, D.S. (2008). NELF and GAGA factor are linked to promoter-proximal pausing at many genes in Drosophila. Mol. Cell. Biol. 28, 3290–3300.

33. Ni, Z., Saunders, A., Fuda, N.J., Yao, J., Suarez, J.-R., Webb, W.W., and Lis, J.T. (2008). P-TEFb is critical for the maturation of RNA polymerase II into productive elongation in vivo. Mol. Cell. Biol. 28, 1161–1170.

34. Wu, C.-H., Yamaguchi, Y., Benjamin, L.R., Horvat-Gordon, M., Washinsky, J., Enerly, E., Larsson, J., Lambertsson, A., Handa, H., and Gilmour, D. (2003). NELF and DSIF cause promoter proximal pausing on the hsp70 promoter in Drosophila. Genes Dev. 17, 1402–1414.

35. Vos, S.M., Pöllmann, D., Caizzi, L., Hofmann, K.B., Rombaut, P., Zimniak, T., Herzog, F., and Cramer, P. (2016). Architecture and RNA binding of the human negative elongation factor. Elife 5. 10.7554/eLife.14981.

36. Narita, T., Yamaguchi, Y., Yano, K., Sugimoto, S., Chanarat, S., Wada, T., Kim, D.-K., Hasegawa, J., Omori, M., Inukai, N., et al. (2003). Human transcription elongation factor NELF: identification of novel subunits and reconstitution of the functionally active complex. Mol. Cell. Biol. 23, 1863–1873.

37. Gilchrist, D.A., Nechaev, S., Lee, C., Ghosh, S.K.B., Collins, J.B., Li, L., Gilmour, D.S., and Adelman, K. (2008). NELF-mediated stalling of Pol II can enhance gene expression by blocking promoter-proximal nucleosome assembly. Genes Dev. 22, 1921–1933.

38. Vos, S.M., Farnung, L., Urlaub, H., and Cramer, P. (2018). Structure of paused transcription complex Pol II-DSIF-NELF. Nature 560, 601–606.

39. Su, B.G., and Vos, S.M. (2024). Distinct negative elongation factor conformations regulate RNA polymerase II promoter-proximal pausing. Mol. Cell. 10.1016/j.molcel.2024.01.023.

40. Werner, F. (2012). A Nexus for Gene Expression—Molecular Mechanisms of Spt5 and NusG in the Three Domains of Life. J. Mol. Biol. 417, 13–27.

41. Ponting, C.P. (2002). Novel domains and orthologues of eukaryotic transcription elongation factors. Nucleic Acids Res. 30, 3643–3652.

42. Nguyen, V.T., Kiss, T., Michels, A.A., and Bensaude, O. (2001). 7SK small nuclear RNA binds to and inhibits the activity of CDK9/cyclin T complexes. Nature 414. 10.1038/35104581.

43. Marz, M., Donath, A., Verstraete, N., Nguyen, V.T., Stadler, P.F., and Bensaude, O. (2009). Evolution of 7SK RNA and its protein partners in metazoa. Mol. Biol. Evol. 26, 2821–2830.

44. S, G.B., Gohil, D.S., and Roy Choudhury, S. (2023). Genome-wide identification, evolutionary and expression analysis of the cyclin-dependent kinase gene family in peanut. BMC Plant Biol. 23, 43.

45. Uehara, T.N., Nonoyama, T., Taki, K., Kuwata, K., Sato, A., Fujimoto, K.J., Hirota, T., Matsuo, H., Maeda, A.E., Ono, A., et al. (2022). Phosphorylation of RNA Polymerase II by CDKC;2 Maintains the Arabidopsis Circadian Clock Period. Plant Cell Physiol. 63, 450–462.

46. Cao, L., Chen, F., Yang, X., Xu, W., Xie, J., and Yu, L. (2014). Phylogenetic analysis of CDK and cyclin proteins in premetazoan lineages. BMC Evol. Biol. 14, 10.

47. Gressel, S., Schwalb, B., and Cramer, P. (2019). The pause-initiation limit restricts transcription activation in human cells. Nat. Commun. 10, 3603.

48. Chou, S.-P., Alexander, A.K., Rice, E.J., Choate, L.A., and Danko, C.G. (2022). Genetic dissection of the RNA polymerase II transcription cycle. Elife 11. 10.7554/eLife.78458.

49. Watts, J.A., Burdick, J., Daigneault, J., Zhu, Z., Grunseich, C., Bruzel, A., and Cheung, V.G. (2019). cis Elements that Mediate RNA Polymerase II Pausing Regulate Human Gene Expression. Am. J. Hum. Genet. 105, 677–688.

50. Tome, J.M., Tippens, N.D., and Lis, J.T. (2018). Single-molecule nascent RNA sequencing identifies regulatory domain architecture at promoters and enhancers. Nat. Genet. 50, 1533–1541.

51. Hendrix, D.A., Hong, J.-W., Zeitlinger, J., Rokhsar, D.S., and Levine, M.S. (2008). Promoter elements associated with RNA Pol II stalling in the *Drosophila* embryo. Preprint, 10.1073/pnas.0802406105 10.1073/pnas.0802406105.

52. Liston, D.R., and Johnson, P.J. (1999). Analysis of a Ubiquitous Promoter Element in a Primitive Eukaryote: Early Evolution of the Initiator Element. Mol. Cell. Biol. 19, 2380.

53. Ngoc, L.V., Cassidy, C.J., Huang, C.Y., Duttke, S.H.C., and Kadonaga, J.T. (2017). The human initiator is a distinct and abundant element that is precisely positioned in focused core promoters. Genes Dev. 31, 6.

54. Inactivation of Yeast Isw2 Chromatin Remodeling Enzyme Mimics Longevity Effect of Calorie Restriction via Induction of Genotoxic Stress Response (2014). Cell Metab. 19, 952–966.

55. Persson, J., Steglich, B., Smialowska, A., Boyd, M., Bornholdt, J., Andersson, R., Schurra, C., Arcangioli, B., Sandelin, A., Nielsen, O., et al. (2016). Regulating retrotransposon activity through the use of alternative transcription start sites. EMBO Rep. 17, 753.

56. Gilchrist, D.A., Dos Santos, G., Fargo, D.C., Xie, B., Gao, Y., Li, L., and Adelman, K. (2010). Pausing of RNA polymerase II disrupts DNA-specified nucleosome organization to enable precise gene regulation. Cell 143, 540–551.

57. Kundaje, A., Kyriazopoulou-Panagiotopoulou, S., Libbrecht, M., Smith, C.L., Raha, D., Winters, E.E., Johnson, S.M., Snyder, M., Batzoglou, S., and Sidow, A. (2012). Ubiquitous heterogeneity and asymmetry of the chromatin environment at regulatory elements. Genome Res. 22, 1735.

58. Lantermann, A.B., Straub, T., Strålfors, A., Yuan, G.-C., Ekwall, K., and Korber, P. (2010). Schizosaccharomyces pombe genome-wide nucleosome mapping reveals positioning mechanisms distinct from those of Saccharomyces cerevisiae. Nat. Struct. Mol. Biol. 17, 251–257.

59. Bintu, L., Ishibashi, T., Dangkulwanich, M., Wu, Y.-Y., Lubkowska, L., Kashlev, M., and Bustamante, C. (2012). Nucleosomal elements that control the topography of the barrier to transcription. Cell 151, 738– 749.

60. DeBerardine, M., Booth, G.T., Versluis, P.P., and Lis, J.T. (2023). The NELF pausing checkpoint mediates the functional divergence of Cdk9. Nat. Commun. 14, 2762.

61. Fant, C.B., Levandowski, C.B., Gupta, K., Maas, Z.L., Moir, J., Rubin, J.D., Sawyer, A., Esbin, M.N., Rimel, J.K., Luyties, O., et al. (2020). TFIID Enables RNA Polymerase II Promoter-Proximal Pausing. Mol. Cell 78, 785–793.e8.

62. Nabet, B., Roberts, J.M., Buckley, D.L., Paulk, J., Dastjerdi, S., Yang, A., Leggett, A.L., Erb, M.A., Lawlor, M.A., Souza, A., et al. (2018). The dTAG system for immediate and target-specific protein degradation. Nat. Chem. Biol. 14, 431–441.

63. Aoi, Y., Smith, E.R., Shah, A.P., Rendleman, E.J., Marshall, S.A., Woodfin, A.R., Chen, F.X., Shiekhattar, R., and Shilatifard, A. (2020). NELF Regulates a Promoter-Proximal Step Distinct from RNA Pol II Pause-Release. Mol. Cell 78, 261–274.e5.

64. Blümli, S., Wiechens, N., Wu, M.-Y., Singh, V., Gierlinski, M., Schweikert, G., Gilbert, N., Naughton, C., Sundaramoorthy, R., Varghese, J., et al. (2021). Acute depletion of the ARID1A subunit of SWI/SNF complexes reveals distinct pathways for activation and repression of transcription. Cell Rep. 37, 109943.

65. Sun, F., Sun, T., Kronenberg, M., Tan, X., Huang, C., and Carey, M.F. (2021). The Pol II preinitiation complex (PIC) influences Mediator binding but not promoter–enhancer looping. Genes Dev. 35, 1175–1189.

66. Nojima, T., Gomes, T., Grosso, A.R.F., Kimura, H., Dye, M.J., Dhir, S., Carmo-Fonseca, M., and Proudfoot, N.J. (2015). Mammalian NET-Seq Reveals Genome-wide Nascent Transcription Coupled to RNA Processing. Cell 161, 526–540.

67. Mahat, D.B., Salamanca, H.H., Duarte, F.M., Danko, C.G., and Lis, J.T. (2016). Mammalian Heat Shock Response and Mechanisms Underlying Its Genome-wide Transcriptional Regulation. Mol. Cell 62, 63–78.

68. Adelman, K., Kennedy, M.A., Nechaev, S., Gilchrist, D.A., Muse, G.W., Chinenov, Y., and Rogatsky, I. (2009). Immediate mediators of the inflammatory response are poised for gene activation through RNA polymerase II stalling. Proc. Natl. Acad. Sci. U. S. A. 106, 18207–18212.

69. Rahl, P.B., Lin, C.Y., Seila, A.C., Flynn, R.A., McCuine, S., Burge, C.B., Sharp, P.A., and Young, R.A. (2010). c-Myc regulates transcriptional pause release. Cell 141, 432–445.

70. Vihervaara, A., Mahat, D.B., Guertin, M.J., Chu, T., Danko, C.G., Lis, J.T., and Sistonen, L. (2017). Transcriptional response to stress is pre-wired by promoter and enhancer architecture. Nat. Commun. 8, 255.

71. Duarte, F.M., Fuda, N.J., Mahat, D.B., Core, L.J., Guertin, M.J., and Lis, J.T. (2016). Transcription factors GAF and HSF act at distinct regulatory steps to modulate stress-induced gene activation. Genes Dev. 30, 1731–1746.

72. Ghosh, S.K.B., Missra, A., and Gilmour, D.S. (2011). Negative elongation factor accelerates the rate at which heat shock genes are shut off by facilitating dissociation of heat shock factor. Mol. Cell. Biol. 31, 4232–4243.

73. Lis, J.T., Mason, P., Peng, J., Price, D.H., and Werner, J. (2000). P-TEFb kinase recruitment and function at heat shock loci. Genes Dev. 14, 792.

74. Peterlin, B.M., and Price, D.H. (2006). Controlling the elongation phase of transcription with P-TEFb. Mol. Cell 23, 297–305.

75. Chen, F., Gao, X., and Shilatifard, A. (2015). Stably paused genes revealed through inhibition of transcription initiation by the TFIIH inhibitor triptolide. Genes Dev. 29, 39–47.

76. Henriques, T., Gilchrist, D.A., Nechaev, S., Bern, M., Muse, G.W., Burkholder, A., Fargo, D.C., and Adelman, K. (2013). Stable pausing by RNA polymerase II provides an opportunity to target and integrate regulatory signals. Mol. Cell 52, 517–528.

77. Versluis, P., Graham, T.G.W., Eng, V., Ebenezer, J., Darzacq, X., Zipfel, W.R., and Lis, J.T. (2024). Live-cell imaging of RNA Pol II and elongation factors distinguishes competing mechanisms of transcription regulation. Mol. Cell 84. 10.1016/j.molcel.2024.07.009.

78. 7SK RNA, a non-coding RNA regulating P-TEFb, a general transcription factor 10.4161/rna.6.2.8115.

79. P-TEFb: The master regulator of transcription elongation (2023). Mol. Cell 83, 393–403.

80. C. Quaresma, A.J., Bugai, A., and Barboric, M. (2016). Cracking the control of RNA polymerase II elongation by 7SK snRNP and P-TEFb. Nucleic Acids Res. 44, 7527–7539.

81. Parfrey, L.W., Lahr, D.J.G., Knoll, A.H., and Katz, L.A. (2011). Estimating the timing of early eukaryotic diversification with multigene molecular clocks. Proc. Natl. Acad. Sci. U. S. A. 108, 13624–13629.

82. Roger, A.J., and Hug, L.A. (2006). The origin and diversification of eukaryotes: problems with molecular phylogenetics and molecular clock estimation. Philos. Trans. R. Soc. Lond. B Biol. Sci. 361, 1039–1054.

83. Scott, T.G., Martins, A.L., and Guertin, M.J. (2022). Processing and evaluating the quality of genome-wide nascent transcription profiling libraries. bioRxiv, 2022.12.14.520463. 10.1101/2022.12.14.520463.

84. Core, L.J., Martins, A.L., Danko, C.G., Waters, C.T., Siepel, A., and Lis, J.T. (2014). Analysis of nascent RNA identifies a unified architecture of initiation regions at mammalian promoters and enhancers. Nat. Genet. 46, 1311–1320.

85. Mahat, D.B., Kwak, H., Booth, G.T., Jonkers, I.H., Danko, C.G., Patel, R.K., Waters, C.T., Munson, K., Core, L.J., and Lis, J.T. (2016). Base-pair-resolution genome-wide mapping of active RNA polymerases using precision nuclear run-on (PRO-seq). Nat. Protoc. 11, 1455–1476.

86. Arenas-Mena, C., Miljovska, S., Rice, E.J., Gurges, J., Shashikant, T., Wang, Z., Ercan, S., and Danko, C.G. (2021). Identification and prediction of developmental enhancers in sea urchin embryos. BMC Genomics 22, 751.

87. Lewis, J.J., Cicconardi, F., Martin, S.H., Reed, R.D., Danko, C.G., and Montgomery, S.H. (2021). The Dryas iulia Genome Supports Multiple Gains of a W Chromosome from a B Chromosome in Butterflies. Genome Biol. Evol. 13. 10.1093/gbe/evab128.

88. Stefanik, D.J., Friedman, L.E., and Finnerty, J.R. (2013). Collecting, rearing, spawning and inducing regeneration of the starlet sea anemone, Nematostella vectensis. Nat. Protoc. 8, 916–923.

89. Chu, T., Rice, E.J., Booth, G.T., Salamanca, H.H., Wang, Z., Core, L.J., Longo, S.L., Corona, R.J., Chin, L.S., Lis, J.T., et al. (2018). Chromatin run-on and sequencing maps the transcriptional regulatory landscape of glioblastoma multiforme. Nat. Genet. 50, 1553–1564.

90. Smith, J.P., Dutta, A.B., Sathyan, K.M., Guertin, M.J., and Sheffield, N.C. (2021). PEPPRO: quality control and processing of nascent RNA profiling data. Genome Biol. 22, 155.

91. DeBerardine, M. (2022). BRGenomics: Tools for the Efficient Analysis of High-Resolution Genomics Data.

92. R Core Team (2022). R: A Language and Environment for Statistical Computing.

93. Lawrence, M., Gentleman, R., and Carey, V. (2009). rtracklayer: an R package for interfacing with genome browsers. Bioinformatics 25, 1841–1842.

94. Lee, S., Cook, D., and Lawrence, M. (2019). plyranges: a grammar of genomic data transformation. Genome Biol. 20, 4.

95. Pagès, H. (2022). BSgenome: Software infrastructure for efficient representation of full genomes and their SNPs.

96. Wickham, H., Averick, M., Bryan, J., Chang, W., McGowan, L., François, R., Grolemund, G., Hayes, A., Henry, L., Hester, J., et al. (2019). Welcome to the tidyverse. J. Open Source Softw. 4, 1686.

97. Zeileis, A., and Grothendieck, G. (2005). zoo: S3 Infrastructure for Regular and Irregular Time Series. J. Stat. Softw. 14, 1–27.

98. Wickham, H. (2016). ggplot2: Elegant Graphics for Data Analysis (Springer).

99. Ramírez, F., Ryan, D.P., Grüning, B., Bhardwaj, V., Kilpert, F., Richter, A.S., Heyne, S., Dündar, F., and Manke, T. (2016). deepTools2: a next generation web server for deep-sequencing data analysis. Preprint, 10.1093/nar/gkw257 10.1093/nar/gkw257.

100. Richter, D.J., Berney, C., Strassert, J.F.H., Poh, Y.-P., Herman, E.K., Muñoz-Gómez, S.A., Wideman, J.G., Burki, F., and de Vargas, C. (2022). EukProt: A database of genome-scale predicted proteins across the diversity of eukaryotes. Peer Community J. 2. 10.24072/pcjournal.173.

101. Parks, D.H., Chuvochina, M., Waite, D.W., Rinke, C., Skarshewski, A., Chaumeil, P.-A., and Hugenholtz, P. (2018). A standardized bacterial taxonomy based on genome phylogeny substantially revises the tree of life. Nat. Biotechnol. 36, 996–1004.

102. Parks, D.H., Chuvochina, M., Chaumeil, P.-A., Rinke, C., Mussig, A.J., and Hugenholtz, P. (2020). A complete domain-to-species taxonomy for Bacteria and Archaea. Nat. Biotechnol. 38, 1079–1086.

103. Finn, R.D., Bateman, A., Clements, J., Coggill, P., Eberhardt, R.Y., Eddy, S.R., Heger, A., Hetherington, K., Holm, L., Mistry, J., et al. (2014). Pfam: the protein families database. Nucleic Acids Res. 42, D222– D230.

104. Li, W., and Godzik, A. (2006). Cd-hit: a fast program for clustering and comparing large sets of protein or nucleotide sequences. Bioinformatics 22, 1658–1659.

105. Fu, L., Niu, B., Zhu, Z., Wu, S., and Li, W. (2012). CD-HIT: accelerated for clustering the next-generation sequencing data. Bioinformatics 28, 3150–3152.

106. Katoh, K., Misawa, K., Kuma, K.-I., and Miyata, T. (2002). MAFFT: a novel method for rapid multiple sequence alignment based on fast Fourier transform. Nucleic Acids Res. 30, 3059–3066.

107. Katoh, K., Kuma, K.-I., Toh, H., and Miyata, T. (2005). MAFFT version 5: improvement in accuracy of multiple sequence alignment. Nucleic Acids Res. 33, 511–518.

108. Katoh, K., and Standley, D.M. (2013). MAFFT multiple sequence alignment software version 7: improvements in performance and usability. Mol. Biol. Evol. 30, 772–780.

109. Eddy, S.R. (2020). HMMer.

110. Capella-Gutiérrez, S., Silla-Martínez, J.M., and Gabaldón, T. (2009). trimAl: a tool for automated alignment trimming in large-scale phylogenetic analyses. Bioinformatics 25, 1972–1973.

111. Pagès H, Aboyoun P, Gentleman R, DebRoy S (2022). Biostrings: Efficient manipulation of biological strings.

112. Ivanek, O.B.A. (2023). seqLogo: Sequence logos for DNA sequence alignments.

113. Letunic, I., and Bork, P. (2007). Interactive Tree Of Life (iTOL): an online tool for phylogenetic tree display and annotation. Bioinformatics 23, 127–128.

114. Adl, S.M., Bass, D., Lane, C.E., Lukeš, J., Schoch, C.L., Smirnov, A., Agatha, S., Berney, C., Brown, M.W., Burki, F., et al. (2019). Revisions to the Classification, Nomenclature, and Diversity of Eukaryotes. J. Eukaryot. Microbiol. 66, 4–119.

115. Ahlmann-Eltze, C., and Patil, I. (2021). ggsignif: R Package for Displaying Significance Brackets for “ggplot2.” PsyArXiv. 10.31234/osf.io/7awm6.

116. Neuwirth, E. (2022). RColorBrewer: ColorBrewer Palettes.

117. Wilke, C.O. (2020). cowplot: Streamlined Plot Theme and Plot Annotations for “ggplot2.”

