## Supplementary material for "Evolution of promoter-proximal pausing enabled a new layer of transcription control": Other supplementary files.: SupplementaryMethods_Leger.docx

**Detailed supplementary methods for pausing machinery component homolog searches**

For all Basic Local Alignment Search Tool (BLAST) searches (BLAST v. 2.13.0; Altschul et al. 1990 https://doi.org/10.1016/S0022-2836(05)80360-2) and profile Hidden Markov Model (HMM)-based searches (HMMer 3.3.2; Eddy et al. 2011 doi: 10.1371/journal.pcbi.1002195), three databases were queried locally:

1) A custom database of eukaryotic sequences generated from the protein sequences contained in the eukaryotic predicted proteome database EukProt v.3 (Richter et al. 2022 doi: 10.24072/pcjournal.173), with all metazoan proteomes removed and replaced with the subset of metazoan proteomes found in The Comparative Dataset (EP00058_Branchiostoma_floridae; EP00059_Ciona_intestinalis; EP00067_Danio_rerio; EP00074_Homo_sapiens; EP00076_Gallus_gallus; EP00081_Caenorhabditis_elegans; EP00090_Calanus_glacialis; EP00096_Caligus_rogercresseyi; EP00099_Drosophila_melanogaster; EP00103_Capitella_teleta; EP00110_Nematostella_vectensis; EP00114_Trichoplax_sp_H2; EP00115_Mnemiopsis_leidyi; EP00118_Oscarella_pearsei; EP00119_Amphimedon_queenslandica;)
2) The archaeal sequences section of the Genome Taxonomy Database (GTDB) release 207 (Parks et al. 2018 <https://doi.org/10.1038/nbt.4229>; Rinke et al. 2021 doi:10.1038/s41564-021-00918-8)
3) The bacterial sequences section of the GTDB release 207

In addition, targeted species-specific HMM searches were conducted against the predicted proteomes of the individual organisms in which PRO-Seq experiments had been conducted, in order to try to identify additional possible homologs:

*Dictyostelium discoideum*: EukProt v.3 EP00023

*Strongylocentrotus purpuratus*: EukProt v.3 EP00057

*Petromyzon marinus*: EukProt v.3 EP00060

*Mus musculus*: EukProt v.3 EP00072

*Homo sapiens*: EukProt v.3 EP00074

*Caenorhabditis elegans*: EukProt v.3 EP00081

*Daphnia pulex*: EukProt v.3 EP00085

*Drosophila melanogaster*: EukProt v.3 EP00099

*Nematostella vectensis*: EukProt v.3 EP00110

*Capsaspora owczarzaki*: EukProt v.3 EP00120

*Sphaeroforma arctica*: (separately) EukProt v.3 EP00125 and Sphaeroforma arctica_v4 PacBio assembly-derived predicted proteome

*Creolimax fragrantissima*: EukProt v.3 EP01134

*Saccharomyces cerevisiae*: EukProt v.3 EP00144

*Schizosaccharomyces pombe*: EukProt v.3 EP00145

*Arabidopsis thaliana*: EukProt v.3 EP00260

*Oryza sativa*: EukProt v.3 EP00265

*Zea mays*: EukProt v.3 EP00267

*Haloferax mediterranei*: GCF_005406325.1 (predicted proteins from genome assembly ASM540632v1)

*Escherichia coli* strain K-12 substrain MG1655: GCF_000005845.2 (predicted proteins from genome assembly ASM584v2)

Because no predicted proteome was available from *Dryas iulia* in public databases, TBLASTN searches were conducted against its genome (Dryas iulia helico3) using the http://blast.lepbase.org/ webserver (Challi et al. 2016 <https://doi.org/10.1101/056994>) with *Drosophila melanogaster* queries and default parameters.

Initial BLASTP searches were conducted using an e-value cut-off of 10; in the case of CDK9, because of the number of hits retrieved with this cutoff value, a BLASTP search with an e-value cutoff of 1e-10 (i.e., 10^-10^) was instead used. All hits were retrieved, and subjected to reciprocal BLASTP searches against the *Homo sapiens* predicted proteome (EukProt v.3 EP00074) with -max_target_seqs 5, and to Pfam (v. 35; Mistry et al. 2021 doi: 10.1093/nar/gkaa913) domain prediction with PfamScan v.1.6.4 (Finn et al. 2014 https://doi.org/10.1093/nar/gkt1223).

***NELF-A***

All sequences with a hit to Homo sapiens NELF-A in the top five hits were retrieved. Because vertebrate NELF-A proteins lack a characteristic Pfam domain, all proteins predicted by PfamScan to include a known Pfam domain were excluded.

All remaining sequences were aligned using MAFFT-L-INS-I 7.475 (Katoh and Toh 2009 doi: 10.1093/bib/bbn013 ; Katoh and Standley 2013 doi: 10.1093/molbev/mst010), and the alignment was trimmed manually in Geneious Prime 2023.2.1 to the edge of a well-aligned region of the alignment (corresponding to residues 20-539 of *Homo sapiens* sequence EP00074_Homo_sapiens_P035196). This trimmed alignment was used to build a custom profile HMM (HMM1) using HMMer 3.3.2 with default parameters.

HMM1 was used to conduct searches against each of the databases listed above with HMMer 3.3.2 (program hmmsearch) with a total e-value cut-off of 0.01 and a domain e-value cut-off of 0.3.

All hits were retrieved and clustered to 80% sequence identity using CD-HIT (Li and Godzik 2006 doi: 10.1093/bioinformatics/btl158) to reduce downstream computational time. This reduced number of sequences was again subjected to BLASTP searches against the *Homo sapiens* predicted proteome and PfamScan domain prediction. From the BLASTP search, it was apparent that the HMMer search was identifying a large number of paralogous sequences in addition to candidate NELF-A sequences; further HMM searches were therefore judged not to be necessary. Aligning all sequences obtained following the first HMM search (MAFFT-L-INS-I) and trimming with trimal (Capella-Gutiérrez et al. 2009 doi:10.1093/bioinformatics/btp348) (-gappyout option) did not yield enough sites for phylogenetic reconstruction. We therefore constructed a phylogeny from those sequences with reciprocal best blast hits to *Homo sapiens* NELF-A: sites were aligned using MAFFT-L-INS-I, trimmed with trimal (-gappyout option), and short or poorly aligned sequences were removed; the remaining sequences were again aligned and trimmed, and a phylogeny was inferred in IQ-TREE (multicore version 2.1.3 COVID-edition for Linux 64-bit built Sep 17 2021; Nguyen et al. 2015 doi: 10.1093/molbev/msu300), using the LG substitution matrix, C20 mixture model with weights optimised to each dataset (--mix-opt option), and gamma rate heterogeneity among sites. Support values were calculated from 1000 ultrafast bootstrap approximation replicates (Minh et al. 2013 doi: 10.1093/molbev/mst024), and the Shimodaira-Hasegawa approximate likelihood ratio test (SH-aLRT; Shimodaira and Hasegawa 1999 Mol. Biol. Evol. 16(8):1114–1116. 1999 ; Guindon et al. 2010 doi: 10.1093/sysbio/syq010) with 1000 replicates. This phylogeny was poorly resolved; however, animal sequences formed a clade with NELF-A-like sequences from close relatives of animals (choanoflagellates, the filasterean *Capsaspora owczarzaki*, and ichthyosporeans). While the support for this clade was not robust, this topology is consistent with orthology; we therefore included these sequences in the set of final candidate NELF-A sequences.

We examined sequence alignment over regions of the *Homo sapiens* NELF-A protein previously shown to affect NELF-C/D binding, and pausing activity (Vos et al. 2018 https://doi.org/10.1038/s41586-018-0442-2). The InterProScan (Jones et al. 2014 https://doi.org/10.1093/bioinformatics/btu031) web server was used to generate domain predictions based on profiles contained in the InterPro combined protein family and domain profile database (Blum et al. 2021 <https://doi.org/10.1093/nar/gkaa977>), in order to identify possible HDAg domains. These domains were predicted only in animal and *Capsaspora* candidate NELF-A sequences; however, additional candidates could be aligned over this region (Fig. XX), and are included in our list of NELF-A-like candidates, though we cannot draw firm conclusions about their likely function.

Sequences annotated in public databases as NELF-A were identified in proteomes of apicomplexan parasites such as *Plasmodium* spp. (including the malaria parasite *Plasmodium falciparum*), *Cryptosporidium* spp. and *Babesia* spp. These sequences aligned over the NELF-C/D binding domain, but were truncated, lacking the NELF-A ‘tentacle’ implicated in PolII binding (Vos et al. 2018 https://doi.org/10.1038/s41586-018-0442-2), and failing to align fully to the *Homo sapiens* NELF-A region required for some pausing activity (Narita et al. 2003 <https://doi.org/10.1128/MCB.23.6.1863-1873.2003> ).

***NELF-B***

All sequences with a hit to Homo sapiens NELF-B in the top five hits were retrieved. The list of predicted Pfam domains was examined by eye to identify any sequences containing domains other than the characteristic COBRA1 domain (PF06209.16), and these sequences were removed from the set of putative NELF-B homologs.

This first set of putative homologs was clustered to 80% sequence identify using CD-HIT 4.8.1 and the resulting set of sequences was aligned using MAFFT-L-INS-I 7.475. The alignment was trimmed manually to the edge of the conserved COBRA1 domain in Geneious Prime 2023.2.1. This trimmed alignment was used to build a custom profile HMM (HMM1) using HMMer 3.3.2 with default parameters.

HMM1 was used to conduct searches against each of the databases listed above with HMMer 3.3.2 (program hmmsearch) with a total e-value cut-off of 0.01 and a domain e-value cut-off of 0.3.

All hits were retrieved and clustered to 80% sequence identity using CD-HIT to reduce downstream computational time. This reduced number of sequences was again subjected to BLASTP searches against the *Homo sapiens* predicted proteome and PfamScan domain prediction.

The resulting dataset was aligned to HMM1 with hmmalign (HMMer 3.3.2), trimming unaligned nonhomologous residues from the result with the --trim option, and the resulting alignment was used to construct HMM2 for an iterative HMM search against databases from all three domains, in order to identify any possible prokaryotic NELF-B homologs. Hits were again subjected to a BLASTP search against the *Homo sapiens* predicted proteome and domain prediction. No NELFB candidates could be identified in prokaryotes. All eukaryotic NELF-B candidates were predicted to include the characteristic COBRA1 domain; in addition, all stramenopile sequences examined were predicted to include a C-terminal Bromodomain (PF00439.28), and 7 of 58 green algal sequences examined were predicted to include an N-terminal MOZART1 domain (PF12554.11) followed by 2-3 COBRA1 domains, rather than a single one. These differences from the domain architecture of animal sequences might reflect functional changes; and in the case of green algal sequences, might indicate that functional changes in a shared ancestor of green algae and land plants preceded secondary loss of the NELF complex in land plants. Other domains were predicted in individual sequences, but were not consistently predicted across a group.

***NELF-C/D***

All sequences with a hit to *Homo sapiens* NELF-C/D in the top five hits were retrieved. The list of predicted Pfam domains was examined by eye to identify any sequences containing domains other than the characteristic TH1 domain (PF04858.16), and these sequences were removed from the set of putative NELF-C/D homologs.

This first set of putative homologs was clustered to 80% sequence identify using CD-HIT 4.8.1 and the resulting set of sequences was aligned using MAFFT-L-INS-I 7.475. The alignment was trimmed manually to the edge of the conserved TH1 domain in Geneious Prime 2023.2.1. This trimmed alignment was used to build a custom profile HMM (HMM1) using HMMer 3.3.2 with default parameters.

HMM1 was used to conduct searches against each of the databases listed above with HMMer 3.3.2 (program hmmsearch) with a total e-value cut-off of 0.01 and a domain e-value cut-off of 0.3.

All hits were retrieved and clustered to 80% sequence identity using CD-HIT to reduce downstream computational time. This reduced number of sequences was again subjected to BLASTP searches against the *Homo sapiens* predicted proteome and PfamScan domain prediction.

The resulting dataset was aligned to HMM1 with hmmalign (HMMer 3.3.2), trimming unaligned nonhomologous residues from the result with the --trim option, and the resulting alignment was used to construct a second HMM (HMM2) for an iterative HMM search. Hits to this sequence were again subjected to a BLASTP search against the *Homo sapiens* predicted proteome and domain prediction. No additional NELF-C/D candidates could be identified. No prokaryotic NELF-C/D candidates could be identified; a single bacterial sequence identified as a possible NELFC/D candidate was judged to be a likely green algal contaminant based on its phylogenetic position and on its similarity (>65% sequence identity) to *Micromonas* spp. sequences in tblastn searches against the nonredundant nt database of GenBank. All eukaryotic NELF-C/D candidates identified contained the characteristic NELF-C/D domain; many stramenopile sequences examined additionally contained a variable number of N-terminal WW domains (PF00397.29), possibly consistent with functional changes in this group.

***NELF-E***

All hits were retrieved and clustered to 80% sequence identity using CD-HIT to reduce downstream computational time. This reduced number of sequences was again subjected to BLASTP searches against the *Homo sapiens* predicted proteome and PfamScan domain prediction. No sequences were removed based on domain predictions. Sequences with reciprocal best blast hits to *Homo sapiens* NELF-E were retained, aligned using MAFFT-L-INS-I as above, trimmed to a well-aligned region corresponding to *Homo sapiens* NELF-E positions 252-370, and used to construct a profile HMM, which was used for HMMer searches as above.

HMMer searches yielded a very large number of paralogous sequences in addition to candidate NELF-E sequences, several of which were already present among the initial blastp search hits; further HMM searches were therefore judged not to be necessary. Instead, hits from the initial blastp searches (clustered to 80% sequence identity using CD-HIT), together with hits from species used in PRO-Seq, were used to construct a maximum likelihood phylogeny as above. NELF-E sequences formed a clade sister to poly-A binding proteins (Supplementary Fig XX), albeit without significant support. These candidate NELF-E sequences were found to be shorter than, and easily distinguishable from, all other sequences in the alignment (Supplementary Fig XX).

Whelk parasite *Piridium sociabile* (Alveolata) sequence P002909 is a likely contaminant from its animal host and was not counted in the set of final NELF-E candidates.

***HEXIM***

All sequences with a hit to *Homo sapiens* HEXIM 1 and/or 2 in the top five hits were retrieved. Sequences were aligned using MAFFT-L-INS-I and trimmed manually to the edge of the conserved HEXIM domain (PF15313.9) in Geneious Prime 2023.2.1. This trimmed alignment was used to build a custom profile HMM using HMMer 3.3.2 with default parameters, which was used in searches against all databases; a second iterative HMM search was performed by constructing a second HMM from the resulting hits aligned to HMM1, as above. Candidate HEXIM-like sequences were identified only in eukaryotes. Following these two HMM searches, sequences were clustered to 95% sequence identity using CD-HIT, sequences from species used in PRO-Seq studies were re-added manually, and a phylogeny was inferred as above. Five parasitic alveolate sequences that branch within the animal clade (*Ancora sagittata* P000394, *Eleutheroschizon duboscqi* P035719, *Oxyrrhis marina* P017039, *Hematodinium* sp. SG-2012 P028650 and P032357) are likely contaminants from animal hosts.

In addition to PfamScan, the InterProScan (Jones et al. 2014 https://doi.org/10.1093/bioinformatics/btu031) web server was used to generate domain predictions based on profiles contained in the InterPro combined protein family and domain profile database (Blum et al. 2021 <https://doi.org/10.1093/nar/gkaa977>). HEXIM domains were predicted in all holozoan sequences, and patchily in other HEXIM-like sequences. Holozoan sequences containing HEXIM domains that form a clade are listed as final HEXIM candidates.

***Cyclin-dependent kinase CDK9***

Based on the reciprocal best BLASTP search against the *Homo sapiens* proteome, it was clear that the initial BLASTP search was identifying several families of paralogous cyclin-dependent kinases in addition to CDK9. Because of this, and because CDK9 lacks a characteristic domain that might differentiate it from closely related CDK paralogs, we concluded that a phylogeny would best clarify which taxa likely possess CDK9 one-to-one orthologs.

Sequences with reciprocal best blast hits to *Homo sapiens* CDK sequences were extracted. To reduce computational time, these sequences were clustered at 50% sequence identity using CD-HIT, and sequences from species used in PRO-Seq studies were re-added manually. Bacterial and archaea sequences retrieved are likely homologs of both CDKs and other eukaryotic kinases, based on diamond searches against the non-redundant protein sequence database of the National Center for Biotechnology Information; however, they were retained as an outgroup for the phylogenetic analysis. Sequences were aligned using MAFFT with default parameters, trimmed using trimAl in --gappyout mode. Short and poorly aligned sequences were removed, sequences were realigned and retrimmed, and a preliminary phylogeny was reconstructed using IQ-TREE as above. A clade of eukaryotic sequences contained *Homo sapiens* CDK9 and its closest paralogs, CDK11, CDK12 and CDK13. In an attempt to increase resolution for CDK9, we extracted the corresponding sequences using TREE2FASTA (Sauvage et al. 2018 <https://doi.org/10.1186/s13104-018-3268-y>) and carried out a separate phylogenetic analysis with these sequences using MAFFT-L-INS-I, trimal and IQ-TREE (as above). Opisthokont sequences formed a well-supported clade that we can confidently deem CDK9 orthologs (Fig XX), consistent with Cao et al. 2014 (doi:10.1186/1471-2148-14-10); however, the topology for other eukaryotic sequences was not well resolved, so that we cannot exclude the possibility that CDK9 might have emerged earlier. Although no *Petromyzon marinus* CDK9 sequence was identified in EukProt, a clear ortholog was identified in GenBank and was added to the file of candidate CDK9 sequences.

***Cyclin-T***

Based on the reciprocal best BLASTP search against the *Homo sapiens* proteome, it was clear that hits from the initial BLASTP search included several paralogous cyclins in addition to Cyclin-T proteins. As with CDK9, Cyclin-T proteins lack a characteristic domain that might differentiate them from closely related cyclins; we therefore reconstructed the phylogeny of the cyclin hits from the initial BLASTP search with the aim of pinpointing the origin of Cyclin-T specifically.

Sequences with reciprocal best blast hits to *Homo sapiens* cyclin sequences were extracted. To reduce computational time, they were clustered at 50% sequence identity using CD-HIT, and sequences from species used in PRO-Seq studies were re-added manually. Sequences were aligned using MAFFT-L-INS-I, trimmed with trimAl in --gappyout mode, and a phylogeny was reconstructed using IQ-TREE as above. For the final phylogeny, a clade of cyclins that included animal cyclins T and K was extracted using TREE2FASTA, and analysed separately (with the same alignment, trimming and phylogeny inference methods as above) in order to better resolve its internal topology. This phylogeny yielded two well-supported clades that each contained animal and choanoflagellate sequences, one of which corresponded to Cyclin-T and one to Cyclin-K. Other holozoan sequences grouped with each clade, but because each of these species contained only one cyclin T/K homolog, we cannot exclude the possibility that these are mutual orthologs of both Cyclin-T and Cyclin-K.

***Paf1***

All sequences with a hit to *Homo sapiens* Paf1 in the top five hits were retrieved. Any sequences containing domains other than the characteristic Paf1 domain (PF03985.16) were removed from the set of putative Paf1 homologs.

This first set of putative homologs was clustered to 80% sequence identify using CD-HIT 4.8.1 and the resulting set of sequences was aligned using MAFFT-L-INS-I 7.475. The alignment was trimmed manually to the edge of the conserved Paf1 domain in Geneious Prime 2023.2.1. This trimmed alignment was used to build a custom profile HMM (HMM1) using HMMer 3.3.2 with default parameters.

Two iterative profile HMM searches were conducted as above in an attempt to identify possible Paf1 homologs in Archaea or Bacteria, but no candidates were identified. Eukaryotic sequences with the Paf1 domain and best blast hits to *Homo sapiens* Paf1 identified following the first HMM search were retained as a set of candidate Paf1 sequences; other domains were predicted for only three of these sequences, likely misassembled sequences, and these were excluded from the final set of candidates.

***Spt4***

All sequences with a hit to Homo sapiens Spt4 in the top five hits were retrieved. Any sequences lacking the characteristic Spt4 domain (PF06093.16) were removed from the set of putative Spt4 homologs.

This first set of putative homologs was clustered to 80% sequence identify using CD-HIT 4.8.1 and the resulting set of sequences was aligned using MAFFT-L-INS-I 7.475. The alignment was trimmed manually to the edge of the conserved Spt4 domain in Geneious Prime 2023.2.1. This trimmed alignment was used to build a custom profile HMM (HMM1) using HMMer 3.3.2 with default parameters, which was used in targeted species-specific searches. A second, iterative profile HMM search was conducted as above for possible Spt4 homologs in Bacteria, but no candidates could be identified.

In the absence of a known close paralog of Spt4, sequences containing an Spt4 domain, and with reciprocal best blast hits to *Homo sapiens* Spt4, were retained as a set of candidate Spt4 sequences; no other domains were predicted in these sequences.

***Spt5***

All sequences with a hit to Homo sapiens Spt5 in the top five hits were retrieved. Any sequences lacking the characteristic NGN (PF03439.16), Spt 5 N-terminal (PF11942.11) and/or KOW motif (PF00467.32) domains were removed from the set of putative Spt5 homologs.

This first set of putative homologs was clustered to 80% sequence identify using CD-HIT 4.8.1 and the resulting set of sequences was aligned using MAFFT-L-INS-I 7.475. The alignment was trimmed manually to the N-terminal edge of the Spt5 N-terminal domain and the C-terminal edge of the NGN domain in Geneious Prime 2023.2.1. This trimmed alignment was used to build a custom profile HMM (HMM1) using HMMer 3.3.2 with default parameters. This HMM was used to search all databases as above.

In the absence of a known close paralog of Spt5, sequences containing an NGN domain, and with reciprocal best blast hits to *Homo sapiens* Spt5, were retained as a set of candidate Spt5 sequences. Spt5 N-terminal domains were identified only in eukaryotic sequences, while the NGN domain and KOW motif were identified in all three domains.

***Supplementary files***

Datasets containing examples of confidently predicted candidate orthologs as well ass profile HMM files, trimmed and untrimmed alignments, final domain predictions, and final phylogenies are provided as supplementary materials.

Supplementary material folder contents (files have been renamed for ease of organization):

- *alleval10_taxatrimmed_80.linsi.fasta or *alleval10_taxatrimmed.linsi.fasta: untrimmed alignment used to construct the first profile HMM
- *alleval10_taxatrimmed_80_aligntrimmed.fasta or *alleval10_taxatrimmed_aligntrimmed.fasta: trimmed alignment used to construct the first profile HMM
- *.hmm: first profile HMM
- *alleval10hmm1.trimhmmalign.fasta: newly identified candidate orthologs aligned and trimmed to the first profile HMM, used to construct the second profile HMM
- *.hmm2.hmm: second profile HMM
- Final*candidates_renamed.fasta: file of confidently identified candidate sequences. Because these sequences are derived from clustered data, they are merely representative and not exhaustive lists of all candidates contained in all of the databases searched. Taxonomic information has been added to the original sequence names. Because *Dryas iulia* sequences were identified from searches against genomic scaffolds, with uncertainty remaining around intron boundaries, these sequences were not included in phylogenies.
- Final*candidates_renamed.Pfam35: predicted Pfam35 domains for the sequences contained in each Final*candidates_renamed.fasta file.

***NELF-A, NELF-E, HEXIM, CDK9, CyclinT:***

- *.linsi.fasta or *mafft.fasta: untrimmed alignment used in phylogenetic reconstruction
- *.linsi.trimalgappyout.phy or *mafft.trimalgappyout.phy: trimmed alignment used in phylogenetic reconstruction
- *linsi.gappyout.LGC20G.* or *mafft.gappyout.LGC20G.*: IQ-TREE output files
- *.linsi.gappyout.LGC20G_renamed.tre or *.mafft.gappyout.LGC20G_renamed.tre: final phylogeny, with SH-aLRT and ultrafast bootstrap support values, with taxonomic information added to the original sequence names.
